# A beta-glucosidase of an insect herbivore determines both toxicity and deterrence of a dandelion defense metabolite

**DOI:** 10.1101/2021.03.22.436380

**Authors:** Meret Huber, Thomas Roder, Sandra Irmisch, Alexander Riedel, Saskia Gablenz, Julia Fricke, Peter Rahfeld, Michael Reichelt, Christian Paetz, Nicole Liechti, Lingfei Hu, Ye Meng, Wei Huang, Christelle A. M. Robert, Jonathan Gershenzon, Matthias Erb

## Abstract

Gut enzymes can metabolize plant defense metabolites and thereby affect the growth and fitness of insect herbivores. Whether these enzymes also influence herbivore behavior and feeding preference is largely unknown. We studied the metabolization of taraxinic acid β-D-glucopyranosyl ester (TA-G), a sesquiterpene lactone of the common dandelion (*Taraxacum officinale*) that deters its major root herbivore, the common cockchafer larva (*Melolontha melolontha)*. We demonstrate that TA-G is rapidly deglycosylated and conjugated to glutathione in the insect gut. A broad-spectrum *M. melolontha* β-glucosidase, Mm_bGlc17, is sufficient and necessary for TA-G deglycosylation. Using plants and insect RNA interference, we show that Mm_bGlc17 reduces TA-G toxicity. Furthermore, Mm_bGlc17 is required for the preference of *M. melolontha* larvae for TA-G deficient plants. Thus, herbivore metabolism modulates both the toxicity and deterrence of a plant defense metabolite. Our work illustrates the multifacteted roles of insect digestive enzymes as mediators of plant-herbivore interactions.

## INTRODUCTION

Plants produce an arsenal of toxic secondary metabolites, many of which protect them against phytophagous insects by acting as toxins, digestibility reducers, repellents and deterrents [1]. Insect herbivores commonly metabolize defense metabolites, with important consequences for the toxicity of the compounds [2, 3]. Recent studies identified a series of enzymes that metablize plant defense metabolites and thereby benefit herbivore growth and fitness [4–6]. However, to date, the behavioral consequences of insect metabolism of plant defense metabolites is little understood, despite the importance of behavioral effects of plant defenses for plant fitness and evolution in nature [1,7–9].

Insect enzymes that were identified to metabolize plant defense compounds belong mainly to a few large enzyme classes including the cytochrome P450 monooxygenases, UDP-glycosyltransferases and glutathione S-transferases [10–14]. However, members of other enzyme groups can participate in detoxification, some of which are also involved in primary digestive processes for the breakdown of carbohydrates (β-glucosidases), proteins (proteases) and lipids (lipases). For instance, a *Manduca sexta* β-glucosidase deglycosylates the *Nicotiana attenuata* diterpene glycoside lyciumoside IV, thus alleviating its toxicity [6]. Similarly, the Mexican bean weevil (*Zabrotes subfasciatus*) expresses a protease that degrades α-amylase inhibitors from its host, the common bean (*Phaseolus vulgaris*) [15]. Finally, several insects degrade antinutritional plant protease inhibitors through intestinal proteases [16, 17]. Together, these studies suggest that families of typical digestive enzymes should be examined more carefully for possible roles in the detoxification of plant chemicals.

Enzymes involved in carbohydrate digestion may play a particular role in processing plant defense glycosides. Such compounds are typically considered protoxins, non-toxic, glycosylated precursors that are brought into contact with compartmentalized plant glycosidases upon tissue damage to yield toxic aglycones [18]. Both plant and insect glycosidases may activate plant defense glycosides [3]. The alkaloid glycoside vicine in fava beans for instance is hydrolyzed to the toxic aglucone divicine in the gut of bruchid beetles [19]. Similarly, phenolic glycoside toxins are hydrolysed rapidly by *Papilio glaucus*, the eastern tiger swallowtail. *Papilio glaucus* subspecies adapted to phenolic glycoside containing poplars and willow show significantly lower hydrolysis of these metabolites [20]. Finally, iridoid glycosides from *Plantago* species are hydrolysed and thereby activated by herbivore derived β-glucosidases, and β-glucosidase activity is negatively correlated with host plant adaptation both within and between species [21, 22]. These studies show that herbivore-derived enzymes may cleave plant protoxins and so may be a target of host plant adaptation. However, the genetic basis of protoxin activation by herbivores and the biological consequences of this phenomenon for insect behavior and performance are poorly understood.

Although the deglycosylation of plant defense metabolites is commonly assumed to be disadvantageous for the herbivore, a recent study in *Manduca sexta* showed that deglycosylation of a plant glycoside may decrease rather than increase toxicity [6]. Silencing *M. sexta* β-glucosidase 1 resulted in developmental defects in larvae feeding on *Nicotiana attenuata* plants producing the diterpene glycoside lyciumoside IV (Lyc4), but not in larvae feeding on Lyc4-deficient plants, suggesting that deglycosylation detoxifies rather than activates Lyc4. Although Lyc4 is an atypical defensive glycoside that carries several different sugar moieties and is only partially deglycosylated by *M. sexta*, these results bring up the possibility that defensive activation by glycoside hydrolysis does not necessarily increase the toxicity of these compounds, but may be a detoxification strategy. Clearly more research on how glycoside hydrolysis by digestive enzymes impacts herbivores is needed to understand the role of this process in plant-herbivore interactions [3,6,23].

The herbivore toxins derived from glycoside protoxins have often been investigated for their defensive roles in connection with herbivore growth and development [6,19–22] rather than feeding deterrence, despite the fact that the latter is a well-established mechanism for plant protection in this context [24]. For example, the maize benzoxazinoid glucoside HDMBOA-Glc reduces food intake by *Spodoptera* caterpillars as soon as the glucoside moiety is cleaved off by plant β-glucosidases [25]. Similarly, the deterrent effect of cyanogenic glucosides in *Sorghum* towards *Spodoptera frugiperda* is directly dependent on a functional plant β-glucosidase that releases cyanide upon tissue disruption [26]. Furthermore, different glucosinolate breakdown products have been shown to affect oviposition and feeding choices by *Pieris rapae* and *Trichoplusia ni* [27–29]. However, whether protoxin activation by herbivore-derived enzymes influences herbivore behavior and host plant choice remains unknown.

All protoxin activating enzymes that have been characterized so far in insect herbivores are β-glucosidases, which cleave β-D-glucosides and release free glucose [3]. The primary role of β-glucosidases in insect digestion is to function in the last steps of cellulose and hemicellulose breakdown by converting cellobiose to glucose [30]. Most insect β-glucosidases however also accept other substrates, including various di-and oligosaccharides, glycoproteins and glycolipids, which may help herbivores to obtain glucose from various sources and enable the further breakdown of glycosylated proteins and lipids [31–34]. However, the broad substrate specificity of insect β-glucosidases for plant glucosides with an aryl or alkyl moiety may also result in the activation of defense metabolites, as discussed above [35]. Thus, investigating the substrate specificity and the biochemical function of insect β-glucosidases is important to understand the ecology and evolution of insect mediated protoxin activation.

Known plant protoxins include glucosinolates, salicinoids, and cyanogenic, iridoid and benzoxazinoid glycosides. Plants produce many other types of glycosides that may also be protoxins, but most of these have not yet been carefully investigated for their toxicity or metabolic stability in herbivores. Among these potential protoxins are the bitter-tasing sesquiterpene lactone glycosides. Sesquiterpene lactones form a large group of over 2000 plant defense compounds found principally in the Asteraceae, with glycosides especially common in the latex-producing tribe Cichorieae, which enters the human diet through lettuce, endive and chicory [36]. These substances have a long appreciated role in defense against insect herbivores [37], but it is not clear if glycosylated sesquiterpene lactones should be considered as protoxins that are activated by plant damage.

Here, we studied the metabolism of a sesquiterpene lactone glycoside during the interaction between the common dandelion *Taraxacum officinale* agg. (Asteraceae, Chicorieae) and the larvae of the common cockchafer, *Melolontha melolontha* (Coleoptera, Scarabaeidae) [38, 39]. *Melolontha melolontha* larvae feed on roots of different plant species including members of the Poaceae, Brassicaceae, Salicaceae and Asteraceae, which can contain glycosylated defense compounds such as benzoxazinoids, glucosinolates, and salicinoids, as well as sesquiterpene lactone glycosides [40–43]. The alkaline gut pH of *M. melolontha* (pH = 8.0 – 8.5) possibly facilitates its polyphagous feeding habit by inhibiting the often acidic activating glucosidases of plant protoxins [3, 44]. In the third and final instar, *M. melolontha* prefers to feed on *T. officinale*, which produces large quantities of latex in its roots [41, 45]. The most abundant latex compound, the sesquiterpene lactone glycoside taraxinic acid β-D-glucopyranosyl ester (TA-G), deters *M. melolontha* feeding and thereby benefits plant fitness [7,8,45].

To understand the interaction between TA-G and *M. melolontha*, we first investigated whether TA-G is deglycosylated during insect feeding and whether plant or insect enzymes are involved. We then identified *M. melolontha* β-glucosidases that might hydrolyze TA-G through a comparative transcriptomic approach and narrowed down the list of candidate genes through *in vitro* characterization of heterologously expressed proteins. Finally, we silenced TA-G hydrolysing β-glucosidases in *M. melolontha* through RNA interference (RNAi) and determined the effect of these enzymes on TA-G hydrolysis, toxicity and deterrence *in vivo*. Taken together, our results reveal that β-glucosidases modify the effects of plant defense metabolites on both herbivore performance and behavior, with potentially important consequences for the ecology and evolution of plant-herbivore interactions.

## RESULTS

### TA-G is deglucosylated and conjugated to GSH during M. melolontha feeding

To test if TA-G is hydrolyzed during *M. melolontha* feeding, we analyzed larvae that had ingested defined amounts of TA-G containing *T. officinale* latex. The aglycone TA was not detected in the latex itself, but was present in substantial amounts in the regurgitant and gut of latex-fed larvae. TA-G on the other hand disappeared as soon as the latex was ingested by the larvae (Fig 1A). TA-glutathione (TA-GSH) and TA-cysteine (TA-Cys) were also identified in latex-fed larvae based on mass spectral and NMR data with the Cys sulfhydryl moiety being conjugated to TA at the exocyclic methylene group of the α-methylene-γ-lactone moiety (Figs 1B-C, Figs S1-2). Lower amounts of TA-Cys-Glu and TA-Cys-Gly were also present (Fig S1). No TA-G-GSH or TA-G-Cys conjugates were detected in this experiment. Based on current knowledge of the GSH pathway in insects [46], it is likely that TA is first conjugated to GSH and then cleaved sequentially to form the other metabolites, although some conjugation to GSH prior to deglycosylation may also occur (Fig 1C). Quantitative measurements showed that approximately 25% of the ingested TA-G was converted to GSH conjugates and derivatives (Fig 1D), with TA-Cys accounting for 95% of all identified compounds (Fig 1E). Thus, the deglucosylation and GSH conjugation of TA is a major route for metabolism of this sesquiterpene lactone in *M. melolontha*. The majority of the conjugates accumulated in the anterior midgut (Fig 1D). In contrast to the different body parts, the frass only contained a small fraction of TA conjugates and was dominated by trace quantities of intact TA-G (Figs 1D-E).

**Fig. 1.**
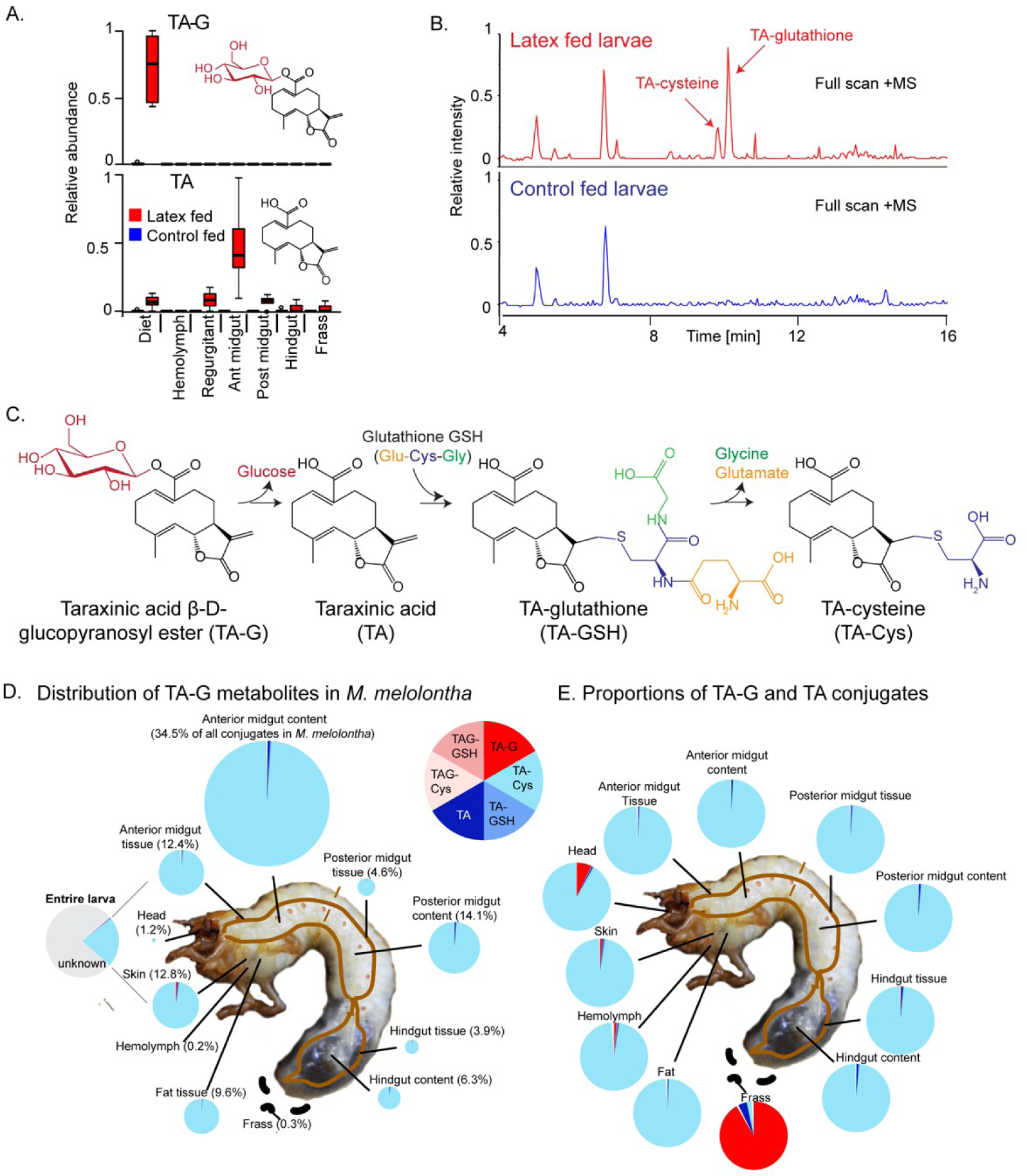
Taraxinic acid β-D-glucopyranosyl ester (TA-G) is rapidly deglucosylated and conjugated to glutathione (GSH) upon ingestion by *Melolontha melolontha*. A. Relative abundance of TA-G and its aglycone taraxinic acid (TA) in diet enriched with *Taraxacum officinale* latex and in *M. melolontha* larval gut, hemolymph and frass after feeding on latex-containing and control diets. Ant = anterior; post = posterior. N = 5. **B.** HPLC-MS full scan (positive mode) of the anterior midgut of *M. melolontha* larvae fed latex-containing and control diets. **C.** Schematic illustration of proposed TA-G metabolism in *M. melolontha*. **D.** Distribution of the total deglucosylated and conjugated metabolites of TA-G in *M. melolontha* larvae that consumed 100 µg TA-G within 24 h. The size of the circles is relative to the total amount of conjugates. Values denote the percentage of metabolites found in each body part and are the mean of 8 replicates. **E.** Relative proportions of TA-G metabolites in quantities from panel D. Values denote the mean of 8 replicates.

### Insect rather than plant enzymes catalyze TA-G deglucosylation in M. melolontha

TA-G deglucosylation may be mediated by plant or insect enzymes or a combination of both. TA-G in *T. officinale* latex incubated at different pH levels at room temperature was readily enzymatically deglucosylated to TA at a pH of 4.6 and 5.4, but not at lower or higher pH values (Fig 2A). As the midgut pH of *M. melolontha* is above 8 (Fig 2B) [44], the deglucosylation of TA-G by plant-derived enzymes is likely inhibited. To test whether TA-G is hydrolyzed by *M. melolontha* enzymes, various *M. melolontha* gut sections were dissected and extracted. Strong deglycosylation activity was detected in the proximal parts of the gut, especially in the anterior midgut (Fig 2B, table S1). TA-G hydrolysis also occurred when larvae were fed with diet containing heat-deactivated latex, which no longer hydrolyzes TA-G itself (Fig 2A and Fig 2C, table S2). Therefore, insect-derived enzymes are sufficient for TA-G deglycosylation in *M. melolontha*.

**Fig. 2.**
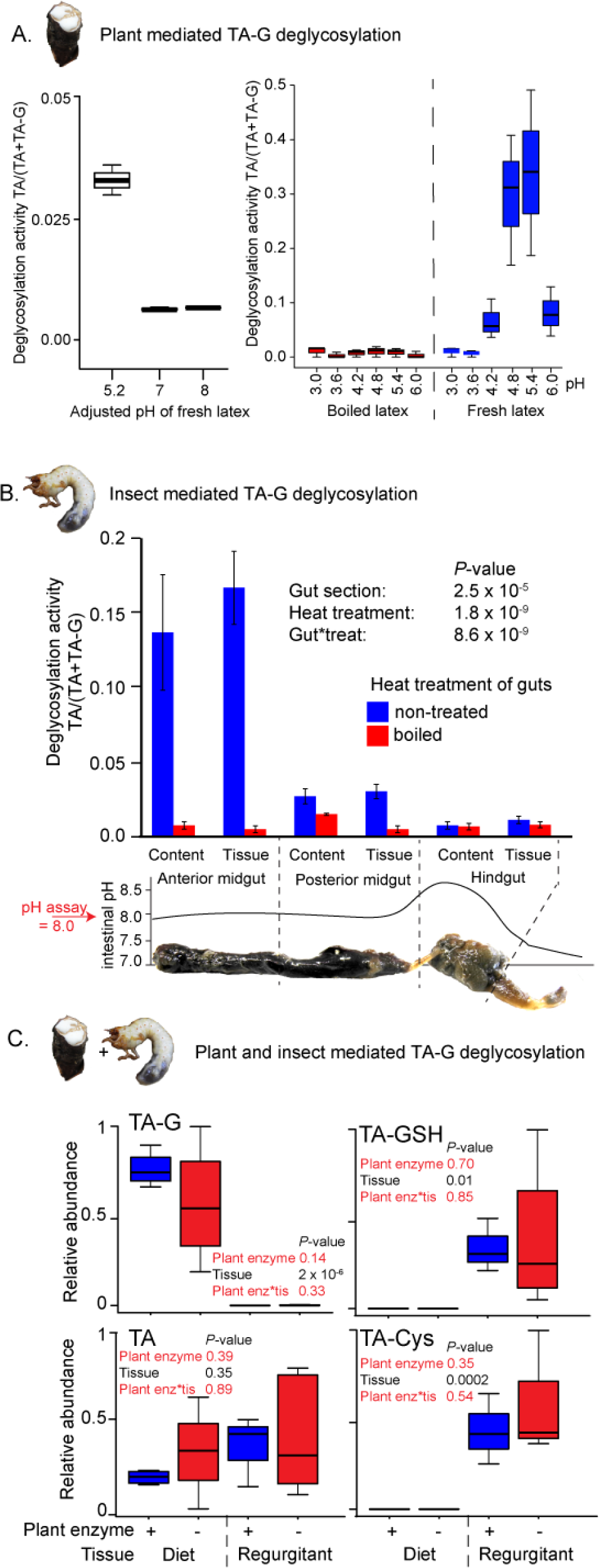
Insect rather than plant enzymes deglucosylate TA-G. **A.** Left and right panel: plant-mediated enzymatic deglycosylation of TA-G at pH 3-8. *Taraxacum officinale* latex was collected from wounded roots and incubated in buffers adjusted to different pH values. N = 3. **B.** Deglucosylation activity of untreated and boiled extracts of *M. melolontha* gut content and gut tissue incubated at pH 8.0 with boiled latex extracts. The *P*-values of a two-way ANOVA are shown. N = 6. Error bars = SEM. The intestinal pH of *M. melolontha* is shown for comparative purposes (data from [44]). **C.** Relative abundance of TA-G and its metabolites in the diet and regurgitant of larvae feeding on carrot slices coated with either intact (+) or heat-deactivated (-) *T. officinale* latex. Heat deactivation of latex did not significantly affect the deglucosylation of TA-G in *M. melolontha*. *P*-values refer to two-way ANOVAs. N = 4. TA-G: taraxinic acid β-D-glucopyranosyl ester; TA = taraxinic acid; GSH = glutathione; Cys = cysteine. Peak area was normalized across all treatments based on the maximal value of each metabolite.

### TA-G hydrolysis is catalyzed by M. melolontha β-glucosidases

As the glucose moiety of TA-G is attached through an ester rather than a glycoside linkage, carboxylesterases or glucosidases may deglycosylate TA-G. TA-G deglycosylation by *M. melolontha* midgut protein extracts was inhibited by the addition of the α-and β-glucosidase inhibitor castanospermine in a dose-dependent manner, but not by the α-glucosidase inhibitor acarbose or the carboxylesterase inhibitor bis(p-nitrophenyl)phosphate (Fig S3). This suggests that β-glucosidases rather than carboxylesterases catalyze TA-G deglucosylation in *M. melolontha*.

### Identification of gut-expressed M. melolontha β-glucosidases

In order to identify TA-G hydrolyzing β-glucosidases, we separately sequenced 18 mRNA samples isolated from anterior and posterior midguts of larvae that had been feeding on diet coated with crude latex, TA-G enriched extracts or water. Putative *M. melolonth*a β-glucosidases were identified based on amino acid similarity to known β-glucosidases from *Tenebrio molitor* and *Chrysomela populi*. 19 sequences with similarity to β-glucosidases had an expression profile matching the observed pattern of high TA-G deglucosylation activity in the anterior midgut. Partial sequences were extended using rapid-amplification of cDNA ends (RACE PCR), resulting in 12 full length β-glucosidases sharing between 55 and 79% amino acid similarity (Fig 3A, text S1 and S2). The remaining seven transcripts could not be amplified or turned out to be fragments of the other candidate genes. All amplified sequences contained an N-terminal excretion signal and possessed the ITENG and NEP motifs characteristic of glucosidases (text S1)[47–49]. Expression levels of the candidate genes were 37 – 308 fold higher in the anterior than posterior midgut samples (*Padj* < 10^-5^, exact tests, n = 3), thus matching the differences in TA-G deglucosylation rate between these gut compartments (Fig 3B). Average expression of the transcripts did not differ among *M. melolontha* larvae fed water, TA-G or latex (Fig 3B, *Padj* > 0.50, exact tests, n = 3).

**Fig. 3.**
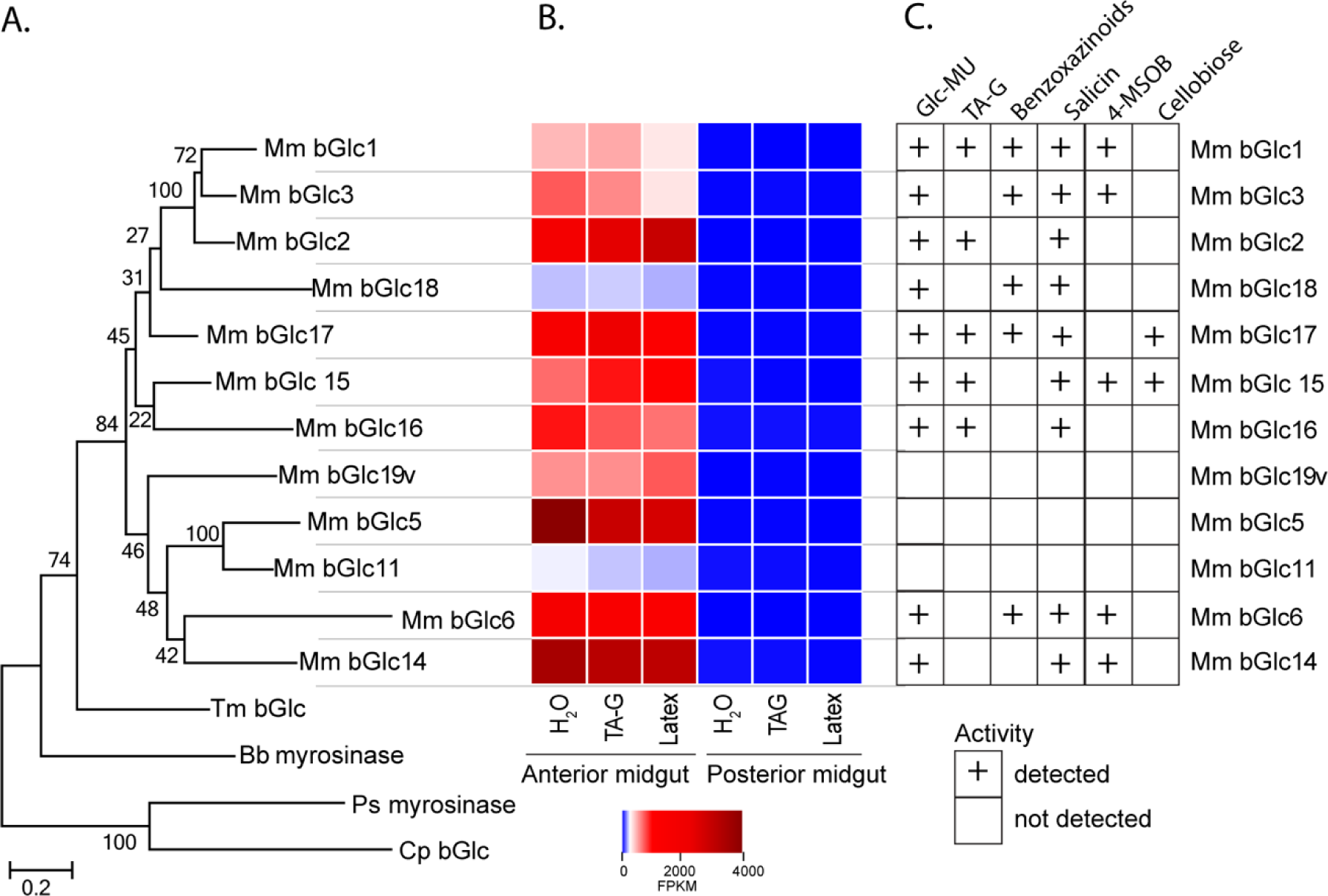
*Melolontha melolontha* midgut β-glucosidases hydrolyze TA-G and other plant defensive glycosides. **A.** Phylogeny of newly identified *M. melolontha* β-glucosidases and previously reported β-glucosidases of *Tenebrio molitor* (Tm bGlc, AF312017.1) and *Chrysomela populi* (Cp bGlc, KP068701.1), and myrosinases (thioglucosidases) of *Phyllotreta striolata* (Ps myrosinase, KF377833.1) and *Brevicoryne brassicae* (Bb myrosinase, AF203780.1) based on amino acid similarities using maximum likelihood method. Bootstrap values (n=1000) are shown next to each node. **B.** Heat map of average (n = 3) gene expression levels of *M. melolontha* β-glucosidases in the anterior and posterior midgut of larvae feeding on diet supplemented with water, TA-G or *T. officinale* latex containing diet. FPKM = Fragments per kilobase of transcript per million mapped reads. **C.** Activity of heterologously expressed *M. melolontha* β-glucosidases with TA-G, a mixture of maize benzoxazinoids, the salicinoid salicin, 4-methylsulfinylbutyl glucosinolate (4-MSOB), cellobiose and the fluorogenic substrate 4-methylumbelliferyl-β-D-glucopyranoside (Glc-MU). Glucosidase activities of three consecutive assays with excreted proteins from insect High Five^TM^ cells were measured. Negative controls (buffer, non-transformed wild type cells and cells transformed with green fluorescent protein) did not hydrolyze any defense metabolite.

### Five M. melolontha β-glucosidases exhibit TA-G hydrolyzing activity

The amplified *M. melolontha* β-glucosidases were heterologously expressed in an insect cell line and assayed with a variety of plant glycosides, including TA-G, benzoxazinoids, a salicinoid and a glucosinolate as well as the disaccharide cellobiose. Nine of the 12 β-glucosidases were active with the standard fluorogenic substrate, 4-methylumbelliferyl-β-D-glucopyranoside, and hydrolyzed at least one of the plant metabolites (Fig 3C, Fig S4). For the three remaining enzymes, we did not observe hydrolysis of any substrate, which could be due to a lack of catalysis or low expression and secretion by the cell line. All tested substrates were deglucosylated by at least one

*M. melolontha* glucosidase (Fig 3C) in agreement with the hydrolysis activity of crude midgut extracts (Fig S4). Five heterologously expressed proteins deglucosylated TA-G (Fig 3C) with the highest TA aglycone formation found for Mm_bGlc17 (Fig S4). Apart from TA-G, Mm_bGlc17also deglycosylated benzoxazinoids, salicin and cellobiose. These data suggest that Mm_bGlc17 and up to four other gut-expressed β-glucosidases may play a role in TA-G metabolism in *M. melolontha*.

### The M. melolontha β-glucosidase Mm_bGlc17 hydrolyzes TA-G in vivo

To test whether *M. melolontha* β*-*glucosidases contribute to TA-G deglucosylation, we silenced two β-glucosidases with TA-G deglycosylation activity, Mm_bGlc16 and Mm_bGlc17, as well as one β-glucosidase without TA-G activity, Mm_bGlc18, by injecting dsRNA targeting a 500 bp fragment of each gene into the second segment of anesthetized *M. melolontha* larvae (Fig S5). After seven days, a stable and specific reduction of the target mRNAs had occurred (Fig S5). TA-G deglucosylation was reduced by 75% in gut extracts of larvae which were silenced in *Mm_bGlc17* (Fig 4A, table S3). Silencing of *Mm_bGlc16* and *Mm_bGlu18* did not significantly reduce TA-G deglucosylation activity compared to GFP controls (Fig 4A). These results confirm that *M. melolontha* derived β-glucosidases hydrolyze TA-G and demonstrate that Mm_Gluc17 accounts for most of the TA-G deglucosylation *in vivo*.

**Fig. 4.**
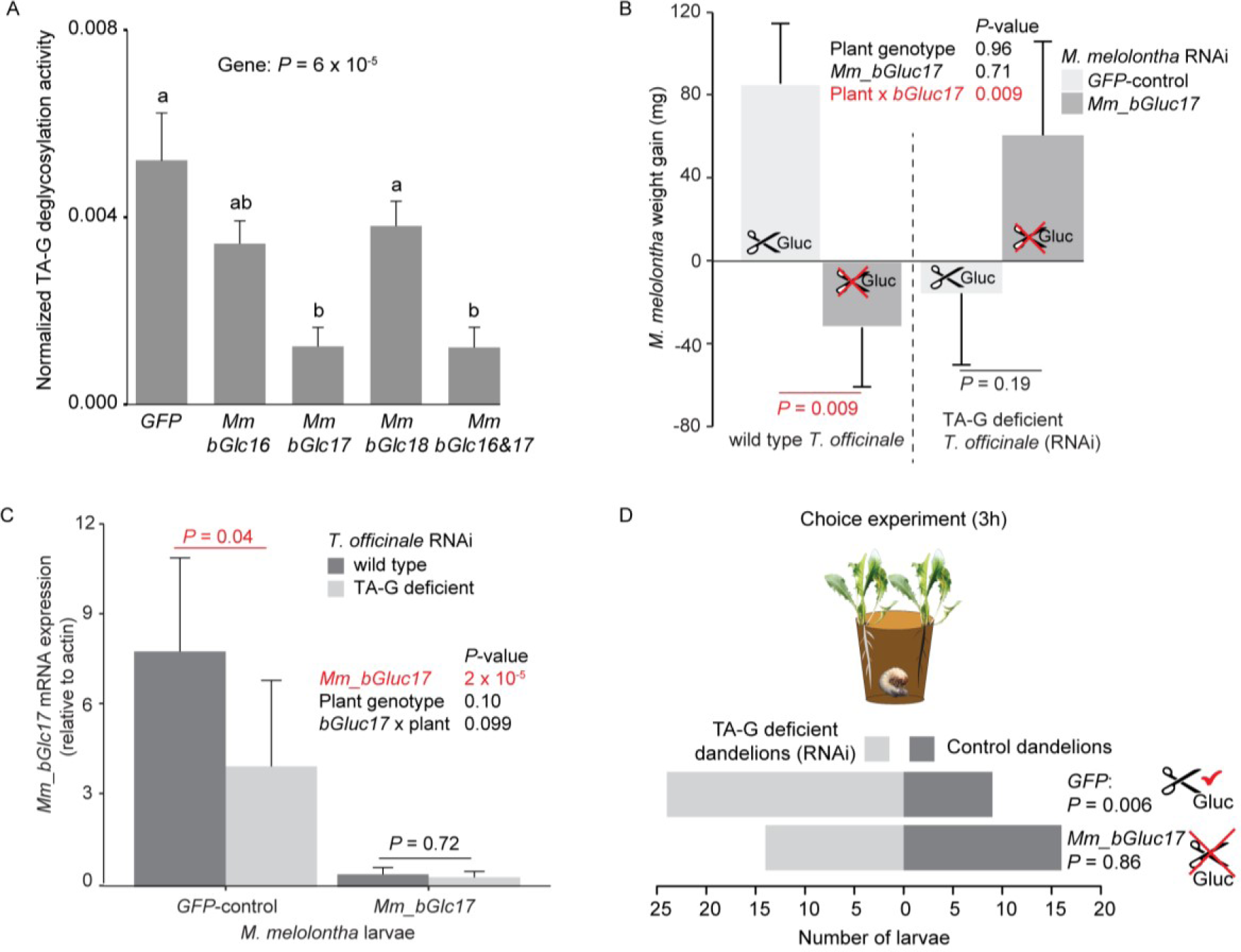
Silencing of *Mm_bGlc17* reduces TA-G deglucosylation and modifies the impact of TA-G on larval growth and behavior. **A.** TA-G deglycosylation activity (TA/(TA+TA-G)) of gut extracts from *M. melolontha* larvae in which different β-glucosidases were silenced through RNA interference (RNAi). Silencing of *Mm_bGlc17* significantly reduced hydrolysis of TA-G by gut extracts. A *GFP* dsRNA construct was used as a negative control. *Mm_bGlc16&17* treated larvae received a 50:50 (v/v) mixture of both dsRNA species. Deglucosylation activity was normalized to that of boiled control samples to correct for the background of non-enzymatic hydrolysis. N = 9-10. *P*-value of a one-way ANOVA is shown. Different letters indicate a significant difference according to a Tukey’s honest significance test. Error bars = SEM. **B.** Weight gain of *Mm_bGlc17*-silenced and *GFP*-control *M. melolontha* larvae growing on transgenic TA-G deficient or control *T. officinale* lines. N = 11-15. *P*-values refer to a two-way ANOVA and Student’s *t*-tests. Error bars = SEM. **C.** Gene expression (relative to actin) of *Mm_bGlc17*-silenced and *GFP*-control *M. melolontha* larvae feeding on transgenic TA-G deficient or control *T. officinale* lines. N = 12-14. *P*-values refer to a two-way ANOVA (log-transformed data) and Kruskal-Wallis rank sum tests (non-transformed values). **D.** Choice of *Mm_bGlc17*-silenced and *GFP*-control larvae between transgenic TA-G deficient and control *T. officinale* lines. Silencing of *Mm_bGlc17* abolished the choice of control larvae for TA-G deficient lines. *P*-values refer to binomial tests.

### Mm_bGluc17 benefits M. melolontha growth on TA-G containing plants

To test whether Mm_bGluc17 modulates the impact of TA-G on larval performance, *Mm_bGlc17* silenced and *GFP*-control larvae were allowed to feed on either TA-G producing wild type or TA-G deficient transgenic dandelions. The interaction of *Mm_bGlc17* silencing and plant genotype significantly affected larval growth (Fig 4B, *P* (*Mm_bGlc17* x TA-G) = 0.009, two-way ANOVA). On TA-G containing plants, *Mm_bGlc17* silencing reduced larval growth, with *GFP*-control larvae gaining 4.5 % body weight and *Mm_bGlc17-*silenced larvae losing 1.4 % body weight (Fig 4B, *P*= 0.009, Student’s *t*-test). By contrast, on TA-G deficient plants *Mm_bGlc17* silencing did not affect larval weight gain (*P* = 0.19, Student’s *t*-test). *GFP*-control *M. melolontha* larvae had higher growth on TA-G containing than TA-G lacking plants (*P* = 0.035, Student’s *t*-test, Fig S6), while the reversed pattern was found in tendency for *Mm_bGlc17* silenced larvae (*P* = 0.099, Student’s *t*-test, Fig S6). The experiment was repeated twice with similar results (Fig S7).

As Mm_bGlc17 benefited larval growth in the presence of TA-G, we investigated whether the expression of this gene is induced by TA-G, and whether its expression correlates with insect performance of *GFP*-larvae on wild type *T. officinale* plants. *Mm_bGlc17* gene expression increased by 95 % on TA-G containing compared to TA-G lacking plants (Fig 4C, *P* = 0.04, Kruskal-Wallis rank sum test). Furthermore, *Mm_bGlc17* expression was positively correlated with *M. melolontha* growth on wild type *T. officinale* plants (*P* = 0.006, linear model, Fig S8), although this pattern was influenced by two data points with high Mm_bGlc17 activity (*P* = 0.20 without outliers, linear model, Fig S8). In contrast, *Mm_bGlc17* expression was not correlated with *M. melolontha* growth on TA-G deficient *T. officinale* plants (*P* = 0.94, linear model, Fig S8). Taken together, these data show that *Mm_bGlc17* expression is induced by TA-G and increases larval performance in the presence of TA-G.

### Mm_bGluc17 expression is required for the deterrent effect of TA-G towards M. melolontha

As TA-G in *T. officinale* latex was previously found to deter *M. melolontha* larvae (24), we tested whether TA-G hydrolysis influences the deterrent properties of TA-G. *Mm_bGlc17* silenced and *GFP*-control larvae were allowed to choose between TA-G producing wild type and TA-G deficient transgenic dandelions. *GFP*-silenced control larvae were deterred by TA-G, with over 60% of the larvae feeding on TA-G deficient plants and 30% on the wild type (Fig 4D, *P* (3h) = 0.006, binomial test; Fig S9). By contrast, *Mm_bGlc17* silenced larvae did not show any preference for TA-G deficient over TA-G producing wild type plants: 44% of the larvae fed on wild type plants, while 42% fed on TA-G deficient plants (Fig 4D, *P* (3h) = 0.86, binomial test). Both patterns were constant over time (Fig S9). *Mm_bGlc17* silencing did not significantly affect the total percentage of larvae that made a choice (86% Mm_bGlc17 vs. 91% GFP). These results demonstrate that *Mm_bGlc17* expression is required for the deterrent effect of TA-G towards *M. melolontha*.

## DISCUSSION

Herbivore enzymes are well known to modify plant defense metabolites, but how these modifications feed back on herbivore performance is often unclear. Furthermore, the behavioral effects of plant defense metabolizations are not understood. Here, we show that an herbivore β-glucosidase deglycosylates a plant secondary metabolite, which modifies both its toxic and deterrent properties and thereby determines the interaction between a plant and its major root-feeding natural enemy.

Metabolization of plant defense metabolites is considered central for the ability of species to overcome chemical defeses of their host plants [2], and recent papers established direct molecular evidence for this concept [4–6]. As the transformation of defense metabolites by insect enyzmes occurs in the gut, metabolization products are considered unlikely to be tasted via frontal sensory structures of insect herbivores, it is thus commonly assumed that there is no direct impact of this process on herbivore behavior [3, 50]. By contrast, transformation of defense metabolites by plant enzymes that are activated by tissue disruption are well accepted to have a strong behavioral impact on insect herbivores, which is in line with the rapid and early formation of plant defense catabolites [25–29]. Here, we find that the insect β-glucosidase Mm_bGlc17, which deglycosylates a defensive sesquiterpene lactone (TA-G) in the insect gut, is also required to elicit the deterrent effect of this metabolite. Our early work on TA-G showed that, in a community context, the capacity of dandelions to produce the glycosylated sesquiterpene lactone reduces *M. melolontha* attack and its negative effect on plant growth and fitness [8], resulting in the selection of high TA-G genotypes under high *M. melolontha* pressure [7]. As these effects are likely the result of the deterrent, rather than the toxic properties of TA-G, they are likely also directly dependent on the presence of Mm_bGlc17. Thus, the metabolism of *M. melolontha* may not only drive the feeding preferences of the herbivore, but also the ecology and evolution of dandelions in their natural habitat. Insect detoxifying enzymes may thus not only shape plant defense evolution by reducing the toxicity of defense compounds, but also by modulating herbivore behavior and host plant choice.

Many plant defensive metabolites are glycosides, which are typically non-toxic themselves but are deglycosylated upon herbivore damage forming toxic products. Both plant-and herbivore-derived β-glucosidases can mediate deglycosylation in the insect gut, but their relative contribution is often unclear [3,19–21]. Here we provide several parallel lines of evidence to demonstrate that the deglucosylation of TA-G, a glycosylated secondary metabolite in the latex of *T. officinale*, depends primarily on β-glucosidases from *M. melolontha* rather than on plant enzymes: First, *T. officinale* TA-G hydrolase activity has an acidic pH optimum (4.8 – 5.4), and activity is very low at the alkaline pH (8.0) found in the gut of *M. melolontha*. Second, TA-G is deglycosylated by *M. melolontha* gut extracts in the absence of plant material, and the presence of plant extract with active TAG glycohydolases does not increase TA-G deglycosylation. Third, *M. melolontha* expresses several β-glucosidase with TA-G – hydrolyzing activity as demonstrated in *in vitro* assays. Fourth, silencing the *M. melolontha* TA-G β-glucosidase Mm_bGlc17 reduces TA-G deglycosylation activity in larval gut extracts. Together, these results demonstrate that insect rather than plant β-glucosidases hydrolyze ingested TA-G in *M. melolontha*.

A long list of plant glycosides are protoxins that are activated by deglycosylation including glucosinolates, benzoxazinoids, salicinoids, alkaloid glycosides, cyanogenic glycosides and iridoid glycosides [1,3,18]. But, until now nothing was known about whether sesquiterpene lactone glycosides are also protoxins. Sesquiterpene lactone aglycones are much more potent than their corresponding glycosides in pharmacological studies of cytotoxicity and anti-cancer activity [51, 52]. However, the consequences of sesquiterpene lactone deglycosylation for herbivore behavior and performance have not been previously investigated [45,53,54]. Our experiments show that deglycosylation of TA-G is associated with an increase rather than a decrease in larval growth on TA-G producing plants. This suggests that the cleavage of TA-G to TA reduces rather than enhances the toxicity of this sesquiterpene lactone. Several explanations for this phenomenon are possible. First, GSH may be more rapidly conjugated by TA than TA-G, and thus deglycosylation is a step towards detoxification. Second, if the target site of TA-G lies in a hydrophilic compartment (such as the gut lumen), deglycosylation may block its activity. Third, the glucose liberated by TA-G deglycosylation may enhance the nutritional quality of dandelion roots for the larvae. When we compared dandelion roots exposed to different plant neighbors in a previous study, we found both positive and negative correlations between root glucose levels and larval growth [55], suggesting a high degree of context dependency. In summary, our results provide evidence that deglycosylation of plant defenses may reduce negative impacts on herbivores. Deglycosylation of a diterpene glycoside of *Nicotiana attenuata* was also found to reduce its toxicity, but in this case the product still contained two other glycoside moieties and thus differs little from its substrate in terms of polarity compared to the differences between TA and TA-G [6].

While Mm_bGlc17 improves larval performance on TA-G producing plants, the enzyme also prompts *M. melolontha* larvae to avoid TA-G. We propose two mechanisms that may be responsible for these counterintuitive results. First, the recognition of TA-G through deglycosylation may guide the *M. melolontha* larva to good feeding sites independently of the toxicity of TA-G. Exploitation of plant secondary metabolites and sugars to locate nutritious tissue has been reported for instance for the specialist root herbivore *Diabrotica virgifera virgifera* feeding on maize roots [56–58]. *Melolontha melolontha* larvae preferentially feed on side roots of dandelions, which contain lower TA-G and higher soluble protein levels than main roots and also may be more nutritious as they are actively growing [8]. Thus, the larvae may not be avoiding TA-G because of its toxicity, but because avoiding high TA-G levels guides them to nutritious roots, with the avoidance behavior being facilitated by *Mm_bGlc17*. A second explanation for the observed patterns may be that herbivore growth by itself gives an incomplete picture regarding the costs of TA-G consumption and metabolism. It has been shown for instance that plant secondary metabolites can enhance larval weight gain, but at the same time increase larval mortality, suggesting that growth is not always beneficial [59, 60]. Furthermore, TA may change the susceptibility of the larvae to parasites and pathogens, as has been shown for the plant volatile indole in maize [61]. Thus, it is possible that under natural conditions, Mm_bGlc17-dependent cleavage of TA-G reduces rather than enhances *M. melolontha* fitness, thus leading to Mm_bGlc17-mediated TA-G avoidance. A more detailed understanding of the role of Mm_bGlc17 and TA-G under natural conditions and over the full 3-year life cycle of *M. melolontha* would help to shed light on these hypotheses.

Interestingly, besides TA-G, Mm_bGluc17 deglycosylates other substrates, including cellobiose and salicinoid and benzoxazinoid defense compounds. The ability of this enzyme to hydrolyze benzoxazinoids seems counterintuitive from the insect’s perspective since benzoxazinoid hydrolysis increases both feeding deterrence as well as toxicity [25,27–29,62], raising the possibility that some plants can co-opt insect enzymes to activate their own defenses. On the other hand, insects are known to have evolved some resistance to plant glycosidic protoxins by inhibiting the activating glycosidases of plants and down-regulating their own activating glycosidases [3,19,20]. The fact that Mm_bGluc17 catalyzes the hydrolysis of a range of glucosides plus the glucose ester TA-G is also unusual. There are only a few previous reports of enzymes with this versatility [63, 64].

The ability of Mm_bGluc17 to mediate hydrolysis of cellobiose, a disaccharide derived from cellulose, suggests its evolutionary origin as a digestive enzyme that was later recruited for processing plant defenses. The relatively large number of β-glucosidases in many insect herbivores [3,6,65] and their species-specific phylogenetic clustering [65] indicate that in addition to contributing to the digestion of cell wall carbohydrates– which are mostly shared among plant species – many β-glucosidases also act on a variety of specialized metabolites, such as plant defense componds. Thus, plant defenses may play an underestimated role in the evolution of β-glucosidases in insect herbivores. Other herbivore digestive enzymes may also interact with plant defenses leading to changes in herbivore performance and behavior, which likely modulate the ecology and evolution of plants and their consumers.

## METHODS

### Plant material

*Taraxacum officinale* plants used for extraction of latex and TA-G were grown in 0.7 – 1.2 mm sand and watered with 0.01 – 0.05% fertilizer with N-P-K of 15-10-15 (Ferty 3, Raselina, Czech Republic) in a climate chamber operating under the following conditions: 16 h light 8 h dark; light supplied by a sodium lamp (EYE Sunlux Ace NH360FLX, Uxbridge, UK); light intensity at plant height: 58 µmol m^2^ s^-1^; temperature: day 22 °C; night 20 °C; humidity: day 55%, night 65%. Depending on the availability, three to five month-old wild type plants of the European A34, 6.56 or 8.13 accession were used unless otherwise indicated [66]. Plants used for the choice experiments were germinated on seedling substrate and transplanted into individual pots filled with potting soil (5 parts ‘Landerde’, 4 parts peat and 1 parts sand) after 2-3 weeks and grown in a climate chamber operating under the following conditions: 16 h light 8 h dark, light supplied by arrays of Radium Bonalux Super NL 39W/840 white lamps; light intensity at plant height: 250 µmol m2 s-1; temperature: day 22°C; night 18°C; humidity 65%. Plant used for the performance experiments were germinated on seedling substrate, transplanted to individual pots filled with a homogenized mixture of 2/3 seedling substrate (Klasmann-Deilmann, Switzerland) and 1/3 landerde (Ricoter, Switzerland) and cultivated in a greenhouse operating under the following conditions: 50%–70% relative humidity, 16/8 hr light/dark cycle, and 24°C at day and 18°C at night, without extrernal light source. The TA-G deficient line RNAi-1 and the control line RNAi-15 were used for these experiments [8].

### Insects

*Melolontha melolontha* larvae were collected from meadows in Switzerland and Germany. Larvae were reared individually in 200 ml plastic beakers filled with a mix of potting soil and grated carrots in a climate chamber operating under the following conditions: 12 h day, 12 h night; temperature: day 13 °C, night 11 °C; humidity: 70%; lighting: none, except for the RNAi experiment, for which the day and night temperature was 4 °C during rearing. All experiments were performed in the dark with larvae in the third larval instar.

### Statistical analysis

All statistical analyses were performed in R version 3.1.1 [67]. Pairwise comparisons were performed with the Agricolae package [68]. Results were displayed with gplots, ggplot2 and RColorBrewer [69–71]. Differential gene expression was analyzed using DeSeq2 and edgeR [72, 73]. Details on the statistical procedure are given in the individual sections.

*Isolation and identification of TA-G metabolites in* M. melolontha *larvae*

In order to test whether TA-G is deglycosylated during digestion in *M. melolontha*, we screened for TA-G, TA and other TA-G metabolites in larvae that fed on either diet supplemented with latex or water. Ten *M. melolontha* larvae were starved for 10 days at room temperature before offering them approximately 0.35 cm^3^ boiled carrot slices that were coated with either main root latex or water. Larvae were allowed to feed for 4 h inside 180 ml plastic beakers covered with a moist tissue paper, after which the frass and regurgitant were collected in 1 ml methanol. Regurgitant was collected by gentle prodding of the larvae. Left-over food was frozen at -80 °C until extraction. The larvae were cooled for 10 min at -20 °C and subsequently dissected on ice to remove the anterior midgut, posterior midgut, hindgut and hemolymph, which were collected in 1 ml methanol. All larval samples were homogenized by vigorously shaking with 2-3 metal beads for 4 min in a paint shaker (Fluid Management, Wheeling, IL, USA), centrifuged at 4 °C for 10 min at 17,000 g and the supernatant was stored at -20 °C until analysis. Left-over food was ground in liquid nitrogen to a fine powder of which 100 mg was extracted with 1 ml methanol by vortexing for 30 s. The samples were subsequently centrifuged at room temperature for 10 min at 17,000 g and the supernatant was stored at -20 °C until analysis. Methanol samples were analyzed on a high pressure liquid chromatograph (HPLC 1100 series equipment, Agilent Technologies, Santa Clara, CA, USA), coupled to a photodiode array detector (G1315A DAD, Agilent Technologies) and an Esquire 6000 ESI-Ion Trap mass spectrometer (Bruker Daltonics, Bremen, Germany). Metabolite separation was accomplished with a Nucleodur Sphinx RP column (250 x 4.6 mm, 5 µm particle size, Macherey–Nagel, Düren, Germany). The mobile phase consisted of 0.2% formic acid (A) and acetonitrile (B) utilizing a flow of 1 ml min^-1^ with the following gradient: 0 min, 10% B, 15 min: 55% B, 15.1 min: 100% B, 16 min: 100% B, followed by column reconditioning [45]. To search for unknown metabolites of TA-G, we visually compared the chromatograms of the anterior midgut of latex and control-fed larvae and subsequently performed MS^2^ experiments using AutoMS/MS runs on the Esquire 6000 ESI-Ion Trap MS to obtain structure information. Using QuantAnalysis (Bruker Daltonics), TA-G, TA and the putative TA-GSH conjugates were quantified based on their most abundant ion trace: TA-G: 685 [M+[M-162]], negative mode, retention time RT = 12.2 min; TA: 263 [M+H], positive mode, RT = 16.8 min); TA-GSH: 570 [M+H], positive mode, RT = 10.1 min ; TA-Cys-glycine: 441 [M+H], positive mode, RT = 9.4 min; TA-Cys-glutamate: 513 [M+H], positive mode, RT = 10.4 min, TA-Cys: 384 [M+H], positive mode, RT = 9.8 min.

### NMR analysis of TA conjugates from M. melolontha midgut extract

In order to identify the structures of the putative TA conjugates, we allowed 15 *M. melolontha* larvae to feed for one month on *T. officinale* plants. Larvae were then recovered and dipped for 2 s in liquid nitrogen before dissecting them on ice. The entire midgut was homogenized in 1 ml methanol by shaking the samples for 3 min with 3 metal beads in a paint shaker. The samples were centrifuged at room temperature for 10 min at 17,000 g, passed through a 0.45 µm cellulose filter, and subsequently purified by HPLC. NMR analyses were conducted using a 500 MHz Bruker Avance HD spectrometer equipped with a 5 mm TCI cryoprobe. Capillary tubes (2 mm) were used for structure elucidation in MeOH-*d4*. The analysis revealed the presence of TA-Cys by comparison with a synthesized standard (see below, Fig S2). Other TA conjugates identified by HPLC-MS were below the detection threshold of NMR.

### Synthesis of TA-G metabolite standards for identification and quantification

In order to characterize and quantify the TA-G metabolites, we isolated and synthesized TA-G, TA-G-GSH, TA-G-Cys, TA, TA-GSH and TA-Cys. TA-G was purified from *T. officinale* latex methanol extracts as described in [8]. TA was obtained by incubating 50 mg purified TA-G with 25 mg β-glucosidase from almonds (Sigma Aldrich) in 2.5 ml H2O at 25 °C for two days. The sample was centrifuged at room temperature for 5 min at 17,000 g and supernatant was discarded. The TA-containing pellet was dissolved in 100 µl dimethylsulfoxide (DMSO) and diluted in 1.9 ml 0.01 M TAPS buffer (pH=8.0). Subsequently, solid phase extraction was performed with a 500 mg HR-X Chromabond cartridge (Macherey-Nagel). The cartridge was washed and conditioned with two volumes of methanol and H2O, respectively. Separation was accomplished using one volume of each H2O, 30% methanol, 60% methanol, and two volumes of 100% methanol. TA eluted in the first 100% methanol fraction, in which no impurities were detected on an Esquire 6000 ESI-Ion Trap-MS. Samples were evaporated under N2 flow at room temperature to almost complete dryness, and 1 ml H2O was added before freeze-drying. To obtain TA-GSH and TA-Cys conjugates, the most abundant TA conjugates in the LC-MS chromatograms, we dissolved 5 mg isolated TA in 5 µl DMSO in two separate Eppendorf tubes and added 1.6 ml 0.01 M TAPS buffer (pH 8.0) and a 75-fold molar excess of either GSH or Cys to the tubes. Similarly, to obtain TA-G-GSH and TA-G-Cys conjugates, we dissolved 5 mg TA-G in 1 ml 0.01 M TAPS (pH=8.0) in two separate Eppendorf tubes and added a 75 molar excess of either GSH or Cys. TA-GSH, TA-G-GSH and TA-G-Cys samples were incubated for 2 days and TA-Cys for 7 days in the dark at 25 °C, after which most of the TA and TA-G had spontaneously conjugated. All samples were stored at -20 °C until purification by semi-preparative HPLC.

Semi-preparative HPLC was accomplished using an HPLC-UV system coupled to a fraction collector (Advantec SF-2120) using a Nucleodur Sphinx RP column (250 x 4.6 mm, 5 µm particle size, Macherey-Nagel). The mobile phase consisted of 0.01% formic acid (A) and acetonitrile (B). Flow rate was set to 1 ml min^-1^ with the following gradient: 0 min: 15% B, 5 min: 30% B, 9 min: 54% B, 9.01 min: 100% B, followed by column reconditioning. Compounds were monitored with a UV detector at 245 nm. As the synthesis resulted in the formation of several isomers that differed in retention times, the conjugates with the same retention times as found in *M. melolontha* larvae were collected. The elution time of the compounds were: TA-G-GSH: 6.9 min; TA-G-Cys: 6.4 min; TA-GSH: 8.6 min; TA-Cys: 8.3 min. The fractions were concentrated under nitrogen flow at 30 °C and subsequently lyophilized. The final yields of the conjugates were: TA-G-GSH: 2.1 mg; TA-G-Cys: 0.38 mg; TA-GSH: 1.47 mg; TA-Cys: 0.23 mg. Purified fractions were analyzed by NMR spectroscopy for structure verification. Structures with chemical shifts are depicted in Fig S2. Standard curves of the conjugates were prepared using 100 µg of the respective compounds in 100% methanol on an Agilent 1200 HPLC system (Agilent Technologies,) coupled to an API 3200 tandem mass spectrometer (Applied Biosystems, Darmstadt, Germany) equipped with a turbospray ion source operating in negative ionization mode. Injection volume was 5 μl. Metabolite separation was accomplished on a ZORBAX Eclipse XDB-C18 column (50 x 4.6mm, 1.8 μm; Agilent Technologies). The mobile phase consisted of 0.05% formic acid (A) and acetonitrile (B) using a flow of 1.1 ml min^-1^ with the following gradient: 0 min: 5% B, 0.5 min: 5% B, 4 min: 55% B, 4.1 min: 90% B, 5 min: 90% B, followed by column reconditioning. The column temperature was kept at 20 °C. The ion spray voltage was maintained at -4.5 keV. The turbo gas temperature was set at 600 °C. Nebulizing gas was set at 50 psi, curtain gas at 20 psi, heating gas at 60 psi and collision gas at 5 psi. Multiple reaction monitoring (MRM) in negative mode monitored analyte parent ion → product ion: m/z 423 → 261 (collision energy (CE) -14 V; declustering potential (DP) -40 V) for TA-G; m/z 730 → 143, (CE -66V; DP -80V) for TA-G-GSH; m/z 544 → 382 (CE -26V; DP -80V) for TA-G-Cys; m/z 261 → 217 (CE -14V; DP -30V) for TA; m/z 568 → 143 (CE -44V; DP -50V) for TA-GSH; m/z 382 → 120 (CE -30V; DP -45V) for TA-Cys; m/z 568 → 143 (CE -44V; DP -50V) for loganic acid. Both Q1 and Q3 quadrupoles were maintained at unit resolution. Analyst 1.5 software (Applied Biosystems) was used for data acquisition and processing. Weight-based response factors of TA-G, TA and their conjugates were calculated relative to loganic acid (Extasynthese, Genay, France). The weight based response factors were as follows: TA-G: 2.8; TA-G-GSH: 2.5, TA-G-Cys: 1.9; TA: 0.3; TA-GSH: 1.9; TA-Cys: 1.1.

### Quantification of M. melolontha TA-G metabolism

In order to quantify the deglucosylation of TA-G and conjugation to GSH, we performed a Waldbauer assay in which we analyzed the TA-G metabolites in *M. melolontha* larvae after consumption of a fixed amount of TA-G. Eight larvae were starved for 7 days before offering them 100 mg artificial diet [8] supplemented with 100 µg purified TA-G, obtained as described above. Larvae were allowed to feed in the dark for 24 h in a 180 ml plastic beaker covered with a moist tissue paper, after which the larvae had completely consumed the food. Frass was collected in 500 µl methanol containing 1 µg*ml^-1^ loganic acid as an internal standard. Subsequently, larvae were dipped for 2 s in liquid nitrogen and the anterior midgut, posterior midgut, hindgut content and tissue, hemolymph and fat tissue removed by dissection. For the gut samples, gut content was collected separately from the gut tissue. All samples were homogenized in 500 µl methanol containing 1 µg*ml^-1^ loganic acid by vigorously shaking the tubes for 2 min with 2-3 metal beads in a paint shaker. All samples were centrifuged at room temperature for 10 min at 17,000 g. Supernatants were analyzed by LC-MS on the API 3200 triple quadrupole mass spectrometer as described above using a 5 µl injection volume. Metabolites were quantified based on loganic acid as an internal standard using Analyst 1.5 software.

### pH dependent hydrolysis of TA-G in T. officinale latex

In order to test whether TA-G is hydrolyzed by plant enzymes, we analyzed the hydrolysis of TA-G in latex that was extracted in buffers that covered the pH range present in the plant vacuole (pH 5), plant cytosol (pH 7) and *M. melolontha* gut (pH 8) [44]. We cut the main roots of *T. officinale* plants 0.5 cm below the stem-root junction and collected the exuding latex of an entire plant in 1 ml 0.05 M MES buffer (pH 5.2), 0.05 M TRIS-HCl buffer (pH 7.0) or 0.05 M TRIS-HCl (pH 8.0), with three replicates for each buffer. Samples were kept at room temperature for 5 min before stopping the reaction by boiling the samples for 10 min at 98 °C, during which TA-G was found to be stable. Samples were centrifuged at room temperature for 10 min at 17,000 g and the supernatant was analyzed by an HPLC 1100 series instrument (Agilent Technologies), coupled to a photodiode array detector (G1315A DAD, Agilent Technologies). Metabolite separation was accomplished as described in [45]. Peak areas for TA-G and its aglycone TA were integrated at 245 nm. As the absorption spectrum of TA-G and TA do not differ, we expressed the deglycosylation activity as the ratio of the peak area of TA/(TA + TA-G). pH-dependent difference in the deglucosylation activity was analyzed using a Kruskal-Wallis rank sum test.

To investigate the precise pH optimum of the plant hydrolases, and to test for spontaneous hydrolysis of TA-G at acidic pH, we extracted *T. officinale* latex in buffers with a pH range of 3-6. Main root latex was collected as described above, extracted in 2 ml H2O containing 20% glycerol and 200 µl extract was immediately suspended in equal volumes of a series of 0.1 M citrate buffers adjusted to pH 3.0, 3.6, 4.2, 4.8, 5.4, and 6.0. Half of the latex-buffer solution was immediately incubated for 10 min at 95 °C to block enzymatic reaction. The remaining samples were kept at room temperature for 15 min to allow enzymatic reaction and subsequently heated for 10 min at 95 °C. Samples were centrifuged at room temperature at 17,000 g and the supernatant was analyzed on HPLC-UV as described above. The peak area of TA-G and TA was integrated at 245 nm and the deglycosylation activity was expressed as TA/(TA + TA-G).

### In vitro deglycosylation of TA-G by M. melolontha gut enzymes

In order to test for the presence of TA-G deglycosylating enzymes in *M. melolontha*, we analyzed the formation of TA in crude extracts of the anterior midgut, posterior midgut and hindgut. Six *M. melolontha* larvae were starved for one week after which they were cooled for 10 min at -20° C before dissection. Larvae were dissected into anterior and posterior midgut and hindgut, with the gut content separated from the gut tissue. Gut samples were weighed and homogenized in 0.01 M TAPS buffer (pH 8.0) containing 10% glycerol with 10 μl per mg tissue using a plastic pistil. For the deglycosylation assay, 30 µl gut samples that had either been kept on ice or boiled for 10 min at 95 °C were incubated with 30 µl latex extract (prepared as described below) for 20 min at 25° C, after which the reaction was stopped by heating the samples for 10 min at 95 °C. Samples were centrifuged at 17,000 g at room temperature for 10 min, after which the supernatant was diluted 1:1 in 0.01M TAPS buffer (pH 8.0) and stored at -20 °C until chemical analysis. Latex extract was obtained by extracting the entire main root latex of six *T. officinale* plants in 6 ml 0.01 M TAPS buffer (pH = 8.0), after which the samples were immediately heated for 10 min at 95 °C. The latex samples were centrifuged for 20 min at 17,000 g and filtered through a 0.45 µm cellulose filter. HPLC-UV analysis and quantification of TA-G and TA was carried out as described above. Deglucosylation activity was expressed as the ratio of TA/(TA+TA-G). Differences between the deglucosylation activity of the gut extract and heat treatment were analyzed with a two-way ANOVA.

### Deglucosylation of TA-G by M. melolontha in vivo in the absence and presence of plant hydrolases

To test whether *M. melolontha* enzymes are sufficient to deglucosylate TA-G, we fed larvae with TA-G supplemented diet that contained *T. officinale* latex extracts that had been left intact or heat deactivated. Eight larvae were starved for two weeks before offering them approximately 0.35 cm^3^ boiled carrot slices coated with 50 µl of intact or heat-deactivated latex extract. Latex extracts were obtained by cutting the main roots of *T. officinale* plants 0.5 cm below the tiller and collecting the latex of an entire plant in 100 µl of either ice-cooled (for intact extracts) or 95 °C (for heat deactivated extracts) H2O. *Melolontha melolontha* larvae were allowed to feed in the dark inside 180 ml beakers covered with soil for four hours. Subsequently, regurgitant was collected in 1 ml methanol by gently prodding of the larvae. Left-over food was frozen in liquid nitrogen, ground to a fine powder and 50 mg ground tissue was extracted with 500 µl methanol by vortexing the samples for 30 s. All samples were centrifuged at room temperature for 10 min at 17,000 g and supernatant analyzed by LC-MS on an Esquire 6000 ESI-Ion Trap-MS (Bruker Daltonics) as described above. TA-G, TA, TA-GSH and TA-Cys were integrated as described above using QuantAnalysis. Statistical differences in the metabolite abundance between the sample type (food, regurgitant) and the presence of active plant enzymes were analyzed with two-way ANOVAs for each metabolite separately.

### Inhibition of TA-G deglycosylation by M. melolontha in vitro

To test whether glucosidases or carboxylesterases mediate the deglucosylation of TA-G, we measured this activity in *M. melolontha* gut extracts in the presence of either carboxylesterase or glucosidase inhibitors. Bis(p-nitrophenyl)phosphate was used as a carboxylesterase inhibitor, whereas castanospermine was deployed as a glucosidase inhibitor that reduces the activity of both α-and β-glucosidases. Six larvae were starved for 12 days before dissection. The anterior midgut content was extracted in 0.01 M TAPS buffer (pH 8.0) containing 10% glycerol using 10 µl per mg gut material. To obtain TA-G as a substrate for the deglycosylation assay, the entire main root latex of each of 15 *T. officinale* plants was collected in 150 µl 0.1 M TAPS (pH 8.0) and samples were immediately heated for 10 min at 95 °C. The samples were centrifuged at room temperature for 10 min at 17,000 g, and the supernatants were pooled and diluted 1:10 in H2O. The enzymatic assay was performed by incubating 10 µl of the diluted latex TAPS extract with 20 µl gut extract and 30 µl 0, 0.002 or 0.2 mM carboxylesterase or glucosidase inhibitor for one hour at room temperature. As a negative control, half volumes of the 0 mM inhibitor samples were immediately incubated at 95 °C to stop the enzymatic reaction. Samples were centrifuged at room temperature for 10 min at 17,000 g and the supernatant was analyzed on an HPLC-UV as described above. TA-G and TA were quantified by integrating the peak area at 245 nm. Deglucosylation activity was expressed as the ratio of TA/(TA + TA-G).

To investigate whether α-or β-glucosidases mediate the hydrolysis of TA-G, we measured deglucosylation activity in *M. melolontha* midgut extracts in the presence of acarbose, a specific α-glucosidase inhibitor, or castanospermine, which inhibits both α-and β-glucosidases. Three L3 *M. melolontha* larvae were starved for 5 days, dipped for 2 s in liquid nitrogen, dissected and the anterior midgut content and tissue were extracted in 10 µl 0.15 M NaCl per mg material. Samples were homogenized with a plastic pistil and centrifuged at 4 °C for 10 min at 17,000 g. Then 20 µl of the supernatant were incubated with 20 µl boiled latex TAPS extract (obtained as described above) and 0.002, 0.2 or 20 mM acarbose or castanospermine (added in 40 µl) for one hour at room temperature. The reaction was stopped by heating for 10 min at 95 °C. Samples were centrifuged at room temperature for 10 min at 17,000 g and the supernatant was analyzed on an HPLC-UV as described above. The peak areas of TA-G and TA were integrated at 245 nm. Deglucosylation activity was expressed as the ratio of TA/(TA + TA-G).

### Transcriptome sequencing and analysis

In order to identify putative *M. melolontha* β-glucosidases, we sequenced 18 anterior and posterior midgut transcriptomes (3 treatments, 2 gut tissues, 3 replicates of each) from larvae feeding on control, TA-G enriched or latex containing diet using Illumina HiSeq 2500. Fifteen *M. melolontha* larvae were starved for 10 days. For three consecutive days, larvae were offered 0.35cm^3^ boiled carrot slices that were coated with either (i) 50 µl water (“control”), (ii) 50 µl latex water extract that contained heat-deactivated latex of the main root of one *T. officinale* plant (“TA-G enriched”) or (iii) the entire main root latex from one *T. officinale* plant (“latex enriched”). The latex water extract was obtained by collecting the main root latex of 15 *T. officinale* plants in a total of 1.5 ml 95 °C hot water. After 15 min incubation at 95 °C, the sample was centrifuged at room temperature for 10 min at 17,000 g and the supernatant was stored at -20 °C. Food was replaced every day. All larvae consumed at least 95% of the offered food during the entire period of the experiment. On the third day, the larvae were dissected four hours after being fed. Larvae were dipped in liquid nitrogen for 2 s and subsequently anterior and posterior midguts were removed by dissection. The gut tissue was cleaned from the gut content, immediately frozen in liquid nitrogen, and stored at -80 °C until RNA extraction. For RNA extraction, gut tissue was ground to a fine powder using plastic pistils. RNA was extracted from 10-20 mg ground tissue using innuPREP RNA Mini Kit (Analytik Jena, Jena, Germany) following the manufacture’s protocol. On column digestion was performed with the innuPREP DNAse Digest Kit (Analytik Jena). TrueSeq compatible libraries were prepared and PolyA enrichment performed before sequencing the transcriptomes on an Illumina HiSeq 2500 with 17 Mio reads per library of 100 base pairs, paired-end. Reads were quality trimmed using Sickle with a Phred quality score of >20 and a minimum read length of 80. De novo transcriptome assembly was performed with the pooled reads of all libraries using Trinity (version Trinityrnaseq_r20131110) running at default settings. Raw reads were archived in the NCBI Sequence Read Archive (SRA) [number to be inserted at a later stage]. Transcript abundance was estimated by mapping the reads of each library to the reference transcriptome using RSEM [74] with Bowtie (version 0.12.9) [75] running at default settings. Differential expression analysis was performed with Wald test in DeSeq2 in which low expressed genes were excluded. GO terms were retrieved using Trinotate and GO enrichment analysis of the up-regulated genes (Benjamini-Hochberg adjusted *P*-value < 0.05) in the anterior midgut of the control and TA-G enriched samples, as well as the control and latex enriched samples, were performed using the hypergeometric test implemented in BiNGO using the Benjamini – Hochberg adjusted *P*-value of < 0.01.

### Identification, phylogenetic and expression analysis of M. melolontha β-glucosidases

In order to identify putative *M. melolontha* β-glucosidases, we performed tBLASTn analysis using the known β-glucosidases from *Tenebrio molitor* (AF312017.1) and *Chrysomela populi* (KP068701.1) as input sequences [34, 76]. We retained transcripts with a BitScore larger than 200, an average FPKM value (all samples) larger than two and an at least two fold higher average FPKM value in the anterior than posterior midguts of the control samples to match the *in vitro* deglycosylation activity. Through this analysis, 19 sequences were selected of which 11 appeared to be full length genes and 8 were gene fragments.

In order to verify the gene sequences, RNA was isolated from *M. melolontha* anterior midgut samples (three biological replicates) using the RNeasy Plant Mini Kit (Qiagen) and single-stranded cDNA was prepared from 1.2 µg of total RNA using SuperScript^TM^ III reverse transcriptase and oligo d(T12-18) primers (Invitrogen, Carlsbad, CA, USA). Rapid-amplification of cDNA ends PCR (‘SMARTer^TM^ RACE cDNA Amplification Kit’ Clontech, Mountain View, CA, USA) was used to obtain full length genes (table S4 for primer information). In the end 12 full length open reading frames of putative β-glucosidases could be amplified from *M. melolontha* cDNA (see table S4 for primer information, text S2 for *M. melolontha* β-glucosidase nucleotide sequences) a reduction from the 19 originally-selected sequences due to lack of amplification of some gene fragments, merging of others, and assembly errors in the transcriptome. Signal peptide prediction of the resulting 12 candidate genes was performed with the online software TargetP (http://www.cbs.dtu.dk/services/TargetP/) [77]. We aligned the amino acid sequences of the 12 candidate sequences, as well as of the known glucosidases of *T. molitor* (AF312017.1), *C. populi* (KP068701.1), *Brevicoryne brassicae* (AF203780.1) and *Phyllotreta striolata* (KF377833.1) [65, 78] using the MUSCLE algorithm (gap open, -2.9; gap extend, 0; hydrophobicity multiplier, 1.2; clustering method, upgmb) implemented in MEGA 5.05 [79], and visualized the alignment in BioEdit version 7.0.9.0 [80]. The alignment was used to compute a phylogeny with a maximum likelihood method (WAG model; gamma distributed rates among sites (5 categories); Nearest-Neighbor-Interchange heuristic method; sites with less than 80% coverage were eliminated) as implemented in MEGA 5.05. A bootstrap resampling strategy with 1000 replicates was applied to calculate tree topology.

In order to estimate the expression levels of the putative β-glucosidases, we replaced the previously identified β-glucosidase sequences in the transcriptome with the confirmed full-length genes and estimated transcript abundance by mapping the trimmed short reads of each library to the corrected reference transcriptome as implemented in the Trinity pipeline using RSEM and Bowtie. For differential expression analysis, all contigs that had an average count value of > 1 per library were retained. To test whether TA-G or latex affected the expression of the β-glucosidases, differential expression analysis was accomplished by pairwise comparisons of the control and TA-G enriched anterior midgut samples, and the control and latex-enriched anterior midgut samples using an exact test in edgeR [72]. The significance level of 0.05 was adjusted for multiple testing using the Benjamini-Hochberg false discovery rate method. To test whether the expression level of β-glucosidases differed between anterior and posterior midgut samples, a pairwise comparison between the control samples of the anterior and posterior midgut was performed as described above. Averaged FPKM values of each treatment and gut section were displayed with a heat map. *Cloning and heterologous expression of* M. melolontha *β-glucosidases* In order to characterize the isolated *M. melolontha* β-glucosidase genes, they were heterologously expressed in a line of *Trichoplusia ni*-derived cells (High Five Cells, Life Technologies, Carlsbad, CA, USA) as described in Rahfeld et al. [76]. Briefly, genes were cloned into the pIB/V5-His TOPO vector (Life Technologies). After sequence verification, these vector constructs were individually used with the FuGeneHD-Kit to transfect insect High Five Cells according to the manufacturer’s instructions (Promega. Madison, WI, USA). After one day of incubation at 27 °C, the cultures were supplied with 60 mg*ml^-1^ blasticidin (Life Technologies) to initiate the selection of stable cell lines. Afterwards, the insect cells were selected over three passages. The cultivation of the stable cell lines for protein expression was carried out in 75 cm^3^ cell culture flasks, containing 10 ml Express Five culture medium (Life Technologies), 20 mg*ml^-1^ blasticidin, one x Protease Inhibitor HP Mix (SERVA Electrophoresis, Heidelberg, Germany). After three days of growth, the supernatant was collected by centrifugation (4000 g, 10 min, 4 °C), concentrated using 10.000 Vivaspin 4 (Sartorius) and desalted (NAP-5, GE Healthcare, Munich, Germany) into assay buffer (100 mM NaPi, pH 8).

### Enzymatic assays of recombinant proteins

In order to test the TA-G hydrolyzing activity and substrate specificity of the *M. melolontha* glucosidases, the heterologously expressed proteins were assayed with the plant defensive glycosides TA-G, a mixture of maize benzoxazinoids (BXDs), salicin and 4-methylsulfinylbutyl glucosinolate (4-MSOB), as well as the disaccharide cellobiose, which were obtained as described below. The standard fluorogenic substrate, 4-methylumbelliferyl-β-D-glucopyranoside, served as a positive control. Non-transformed insect cells (WT) and cells transformed with GFP served as negative controls. For the enzymatic assays, 97 µl concentrated and desalted supernatant of the heterologous expression culture was incubated with 3 µl 10 mM substrate for 24 h at 25 °C, after which the reaction was stopped with an equal volume of methanol. Due to a very rapid deglycosylation of TA-G, incubation time was shortened to 10 s for this compound. After assays, all samples were centrifuged at 11,000 g for 10 min at room temperature and the supernatant analyzed with a different method for each substrate as described below:

TA-G was purified as described in [8]. Deglycosylation activity was measured based on the concentration of the aglycone TA on an HPLC-UV and quantified at 245nm as described above.

BXDs were partially purified from maize seedlings (cultivar Delprim hybrid). Seeds were surface-sterilized and germinated in complete darkness. After 20 days, leaves from approximately 60 seedlings were ground under liquid nitrogen to a fine powder and extracted with 0.1% formic acid in 50% methanol with 0.25 ml per 100 mg tissue. Methanol was evaporated under nitrogen flow at 40 °C. BXDs were enriched using 500 mg HR-X Chromabond solid phase extraction cartridges (Macherey-Nagel) with elution steps (5ml) of water, 30% (aq.) methanol and 100% methanol. Two ml water was added to the 100% methanol fraction, which contained the BXDs. Subsequently, methanol was completely evaporated from this fraction under nitrogen flow at 40 °C, and after freeze-drying, the freeze-dried material (∼5 mg) was dissolved in 1 ml H2O. This enriched BXD solution contained a mixture of different BXD glucosides, with DIMBOA-glucoside as the major compound. To test for the deglycosylation of the BXDs, the formation of the aglycone MBOA (a spontaneous degradation product of the DIMBOA aglycone) was monitored on a an Agilent 1200 HPLC system coupled to an API 3200 tandem mass spectrometer (Applied Biosystems) equipped with a turbospray ion source operating in negative ionization mode. Injection volume was 5 μl using a flow of 1 ml*min^-1^. Metabolite separation was accomplished with a ZORBAX Eclipse XDB-C18 column (50 x 4.6mm, 1.8 μm; Agilent Technologies) using the following gradient of 0.05% formic acid (A) and methanol (B): 0 min: 20% B, 9min: 25% B, 10 min: 50% B, 12 min: 100% B, followed by column reconditioning. The column temperature was kept at 20 °C. MRM was used to monitor analyte parent ion → product ion: m/z 164 → 149 (CE -20 V; DP -24 V) for MBOA. Analyst 1.5 software (Applied Biosystems) was used for data acquisition and processing. Salicin (Alfa Aeser) was purchased and its deglycosylation was quantified based on the formation of the deglycosylation product salicyl alcohol, which was analyzed on an HPLC-UV using the same procedure as described for TA-G. The peak of salicyl alcohol (elution time = 9.3 min) was integrated at 275 nm.

4-MSOB was isolated from 50 g of broccoli seeds (Brokkoli Calabraise, ISP GmbH, Quedlingburg, Germany), which were homogenized in 0.3 L of 80 % aqueous methanol and centrifuged at 2500 g for 10 min, and the supernatant separated on a DEAE-Sephadex A25 column (1 g). After the supernatant was loaded, the column was washed 3 times with 5 mL formic acid + isopropanol + water (3 : 2 : 5 by volume) and 4 times with 5 mL water. Intact glucosinolates were eluted from the DEAE Sephadex with 25 mL of 0.5 M K2SO4 (containing 3 % isopropanol) dropped into 25 mL of ethanol [81]. The collected solution was centrifuged to spin down the K2SO4 and the supernatant was dried under vacuum. The residue was resuspended in 3 mL of water and 4-MSOB was isolated by HPLC as described in [46]. Purification was performed on a Agilent 1100 series HPLC system using a Supelcosil LC-18-DB Semi-Prep column (250x10 mm, 5 µm, Supelco, Bellefonte, PA, USA) with a gradient of 0.1 % (v/v) aqueous trifluoroacetic acid (solvent A) and acetonitrile (solvent B). Separation was accomplished at a flow rate of 4 ml min^-1^ at 25 °C as follows: 0-3% B (6 min), 3-100% B (0.1 min), a 2.9 min hold at 100% B, 100-0% B (0.1 min) and a 3.9 min hold at 0% B, and the fraction containing 4-MSOB was collected with a fraction collector. The fraction was dried under vacuum and resuspended in 10 mL methanol to which 40 mL ethanol was added to precipitate the glucosinolate as the potassium salt. The flask was evaporated under vacuum to remove the solvents and the residue was recovered as a powder. The identity and purity of the isolated 4-MSOB was checked by LC-MS (Bruker Esquire 6000, Bruker Daltonics, Bremen) and 1H NMR (500 MHz model; Bruker BioSpin GmbH, Karlsruhe, Germany). The deglycosylation of 4-MSOB was quantified based on the formation of the 4-MSOB isothiocyanate with an API 3200 LC-MS as described above operating in positive ionization mode.

Injection volume was 5 μl using flow of 1.1 ml*min^-1^. Metabolite separation was accomplished with ZORBAX Eclipse XDB-C18 column (50 x 4.6mm, 1.8 μm; Agilent Technologies) using the following gradient of 0.05% formic acid (A) and acetonitrile (B): 0 min: 3% B, 0.5 min: 15% B, 2.5 min: 85% B, 2.6 min: 100% B, 3.5 min: 100% B, followed by column reconditioning. The column temperature was kept at 20 °C. MRM was used to monitor analyte parent ion → product ion: m/z 178 → 114 (CE -13 V; DP -26 V). Analyst 1.5 software (Applied Biosystems) was used for data acquisition and processing.

Cellobiose (Fluka) was purchased and its deglycosylation was quantified based on the decrease of substrate on an API 3200 LC-MS as described above operating in negative ionization mode. Injection volume was 5 μl using flow of 1 ml*min^-1^. Metabolite separation was accomplished with an apHera NH2 column (15 cm x 4.6 mm x 3 μm) using the following gradient of H2O (A) and acetonitrile (B): 0 min: 20% A, 0.5 min: 20% A, 13 min: 45% A, 14 min: 20% A, followed by column reconditioning. The column temperature was kept at 20 °C. MRM was used to monitor analyte parent ion → product ion: m/z 341 → 161 (CE -10 V; DP -25 V). Analyst 1.5 software was used for data acquisition and processing.

4-methylumbelliferyl-β-D-glucopyranoside (Sigma Aldrich), a fluorogenic substrate served as a rapid positive control for the presence of β-glucosidases. Hydrolysis of 4-methylumbelliferyl-β-D-glucopyranoside was scored visually by the presence of fluorescence in samples excited with UV light at 360 nm using a gel imaging system (Syngene).

Activity of the heterologously expressed β-glucosidases was categorized into presence and absence based on the formation of the respective aglycones of TA-G, BXDs, salicin and 4-MSOB, and the decrease of substrate for cellobiose. For the secondary metabolites, activity was accepted if the aglycone concentration was three fold higher than the mean aglycone concentration of the controls (GFP, WT; except only WT for TA-G). For cellobiose, activity was scored as positive if the cellobiose concentration after the assay was lower than 30% of the cellobiose concentration of the controls (GFP, WT). The enzymatic assays were performed three times (except TA-G only twice) with freshly harvested recombinant proteins within two weeks, which gave similar results (Fig S4). The averaged categorization results are displayed in Fig 3C.

### M. melolontha gut enzymatic assays with plant defensive glycosides

In order to test whether *M. melolontha* gut proteins deglucosylate BXDs, 4-MSOB and salicin, we tested glucohydrolase activity of crude extracts of the anterior midgut *in vitro*. Ten *M. melolontha* larvae were starved for 24 h, after which the larvae were dipped for 2 s in liquid nitrogen and subsequently anterior midgut tissue and gut content were removed by dissection. The samples were extracted with 10 µl ice-cooled 0.1 M TAPS (pH 8.0) per mg material as described above. All samples were centrifuged at 17,000 g for 5 min at 4 °C and the supernatant stored at -20 °C until the enzymatic assay. Deglucosylation activity was measured by incubating 20 µl gut extract that had been either kept on ice or boiled for 10 min at 95 °C with a 6 mM mixture of BXDs, salicin or 4-MSOB (substrates were obtained as described above added in a 20 µl volume) in 0.01 M TAPS (pH 8.0) for 1 h at room temperature, after which the reaction was stopped by the addition of an equal volume of methanol. All samples were centrifuged at 3220 g for 5 minutes at room temperature and the supernatant stored at -20 °C until analysis. For the BXDs, salicin and 4-MSOB, the formation of the aglycone was quantified using HPLC-MS and HPLC-UV as described above. Deglycosylation activity was standardized by dividing the peak area of the aglycone of each sample by the maximal peak area of all samples (“relative aglycone formation”). Differences in the relative aglycone formation between boiled and non-boiled gut samples, as well as between anterior midgut content and tissue samples were analyzed with two-way ANOVAs.

### Development of RNA interference (RNAi) methodology for M. melolontha larvae

In order to establish RNAi in *M. melolontha*, we injected different doses of dsRNA targeting *tubulin* and *GFP* (negative control) into the larvae. As a template for dsRNA synthesis we chose an approximately 500 bp fragment of each gene. The fragments were amplified using the Q5® High-Fidelity DNA Polymerase (New England Biolabs, Ispwich, MA, USA) according to the manufacturer’s procedure and the specific primer combinations Mm-tubulin-fwd and Mm-tubulin-rev for *tubulin*, as well as GFP-RNAi_fwd and GFP-RNAi_rev for *GFP* (table S4). Isolated and purified *M. melolontha* cDNA served as a template for *tubulin*. Plasmid pGJ 2648, which encodes for the emerald variant for *GFP* and was kindly supplied by Dr. Christian Schulze-Gronover, served as a template for *GFP*. Amplified fragments were separated by agarose gel electrophoresis and purified using GeneJET Gel Extraction Kit (Thermo Fisher Scientific, Waltham, MA, USA) according to the manufacturer’s procedure. An A-tail was added using DreamTaq DNA Polymerase (Thermo Fisher Scientific) and the A-tailed fragments were then cloned into T7 promoter sequence containing pCR®2.1-TOPO® plasmids (Life Technologies) according to the manufacturer’s instructions. Plasmids with the insert in both orientations with regard to the T7 promoter were identified by sequencing.

DsRNA was synthesized using the MEGAscript® RNAi Kit (Thermo Fisher Scientific) according to the manufacturer’s procedure. The above described tubulin and GFP plasmid templates were linearized downstream of the insert using the restriction enzymes BamHI (New England Biolabs). Sense and antisense single stranded (ss) RNA was synthesized in separate reactions. The complementary RNA molecules were then annealed and purified using MEGAscript® RNAi Kit according to the manufacturer’s instructions (Thermo Fisher Scientific).

In order to investigate the required dsRNA concentration and duration of the silencing, we injected 2.5 and 0.25 µg dsRNA of *tubulin* or *GFP* per g larva into M*. melolontha*. The larvae were anesthetized under CO2, after which approximately 50 µl tubulin or *GFP* dsRNA (100 ng*µl^-1^ for 2.5 µg per g larva and 10 ng*µl^-1^ for 0.25 µg per g larva) was injected with a sterile syringe (Ø 0.30 x 12 mm) into the hemolymph between the second and third segment of 9 *M. melolontha* larvae per concentration. Every second day, the larvae were weighed. Five days after injection, the larvae received fresh carrots to feed on. Two, five and ten days after injection, three larvae per concentration were frozen in liquid nitrogen. The entire larvae were ground to a fine powder using mortar and pistil under liquid nitrogen and stored at -80°C until RNA extraction. Total RNA was isolated using the GeneJET Plant RNA Purification Kit following the manufacturer’s instructions. On-column RNA digestion was performed with RNase free DNase (Qiagen, Netherlands). cDNA synthesis was performed using SuperScript™ II Reverse Transcriptase (Thermo Fisher) and oligo (dT21) (Microsynth, Switzerland) according to the manufacturer’s instructions. Consequently, the qPCR reaction was performed with the KAPA SYBR® FAST qPCR Kit Optimized for LightCycler® 480 (Kapa Biosystems, Wilmington, MA. USA) in a Nunc™ 96-Well plate (Thermo Fisher Scientific) on a LightCycler® 96 (Roche Diagnostics, Switzerland) with one technical replicate per sample. *Tubulin* gene expression was quantified relative to actin using the qPCR primers qPCR_Mm_Tubulin_fwd and qPCR_Mm_Tubulin_rev for *tubulin*, as well as qPCR_Mm_actin_fwd and qPCR_Mm_actin_rev for *actin* (table S4). Differences in the relative expression of *tubulin* to *actin* and between *tubulin* and *GFP* dsRNA treated larvae were analyzed with a Student’s *t*-test.

### Synthesis of double strand RNA (dsRNA) for RNA interference (RNAi)

In order to test whether Mm_bGlc17 accounts for the TA-G deglycosylation *in vivo*, we silenced *Mm_bGlc16*, *Mm_bGlc17* and *Mm_bGlc18* in *M. melolontha* using RNAi and analyzed TA-G deglycosylation activity *in vitro* using anterior midgut extracts. *Melolontha melolontha* in which a *dsGFP* fragment was injected served as a control. *GFP* dsRNA was synthesized as described above. To obtain dsRNA for the glucosidase genes, we chose approximately 500 bp fragments of *Mm_bGlc16*, *Mm_bGlc17* and *Mm_bGlc18* cDNA as templates for dsRNA synthesis that showed maximal sequence divergence with other *M. melolontha* β-glucosidases as well as among each other (text S3). The fragments were amplified using the Q5® High-Fidelity DNA Polymerase (New England Biolabs) according to the manufacturer’s procedure and specific primer combinations of which one primer was fused to the T7 promoter sequence. The plasmids obtained from the heterologous expression were used as PCR templates (see above). For each β-glucosidase, we performed two PCR reactions to yield two dsRNA templates that are identical except for a single T7 promoter sequence at opposite ends. For *Mm_bGlu16* fragment amplification, the primer combinations Mm_bGlc_16_fwd_T7 and Mm_bGlc_16_rev as well as Mm_bGlc_16_fwd and Mm_bGlc_16_rev_T7 were used. For the amplification of *Mm_bGlc17* and *Mm_bGlc18* fragments, the respective primers were deployed. Amplified fragments were separated by agarose gel electrophoresis and purified using GeneJET Gel Extraction Kit (Thermo Fisher Scientific) according to the manufacturer’s procedure. An A-tail was added using DreamTaq DNA Polymerase (Thermo Fisher Scientific) and the A-tailed fragments were then cloned into pIB/V5-His-TOPO® plasmids. DsRNA was synthesized and linearized as described above using the restriction enzymes Xhol for the glucosidase genes and BamHI for GFP (New England Biolabs). The DsRNA was synthesized using the MEGAscript® RNAi Kit (Thermo Fisher Scientific) according to the manufacturer’s procedure. The above described *M. melolontha* β-glucosidase and *GFP* plasmid templates were linearized downstream of the insert using restriction enzymes XhoI and BamHI (New England Biolabs), respectively, and annealed and purified as described above. *TA-G deglycosylation activity in RNAi silenced* M. melolontha *larvae*

To silence *M. melolontha* glucosidases *in vivo*, we injected dsRNA of the respective glucosidases or *GFP* as a control into *M. melolontha* larvae as described above using 50 µl of a 10 ng*µl^-1^ *Mm_bGlc16*, *Mm_bGlc17*, *Mm_bGlc18* or *GFP* dsRNA. In addition, we performed a co-silencing of *Mm_bGlc16* and *Mm_bGlc17 (Mm_bGlc16&17)*, for which 25 µl 10 ng*µl^-1^ *Mm_bGlc16* and *Mm_bGlc17* were injected. Larvae were kept at room temperature for 7 days, after which the larvae were dissected as described above. The anterior midgut content was extracted with 10 µl 0.01 M TAPS (pH 8.0) per mg material and centrifuged at 17,000 g for 10 min at 4 °C. For the enzymatic reaction, 10 µl supernatant that was either kept at 4 °C or had been boiled for 1 h at 98 °C was incubated with 40 μl 0.01 M TAPS (pH 8.0) and 50 μl 2 mM latex water extract. After 3 h, the reaction was stopped by adding equal volumes of methanol. The samples were centrifuged at 17,000 g for 10 min at room temperature and the supernatant analyzed on a Waters ACQUITY UPLC series equipment coupled to an ACQUITY photodiode array and an ACQUITY QDa mass detector. Metabolite separation was accomplished using a CQUITY UPLC column with 1.7 μm BEH C18 particles (2.1 x 100 mm). The mobile phase consisted of 0.05 % formic acid (A) and acetonitrile (B) utilizing a flow of 0.4 ml*min^-1^ with the following gradient: 0min: 5% B, 1.5min: 20% B, 2.5min: 40% B, 3min: 95% B, 5min: 95% B, followed by column reconditioning. The peak area of TA and TA-G were integrated at 245 nm using Waters MassLynx™^49^. Deglycosylation activity was expressed as the ratio of TA/(TA+TA-G). In addition, to account for the spontaneous deglycosylation of TA-G, the deglycosylation activity was normalized by subtracting the average TA/(TA+TA-G) of the boiled samples from each non-boiled sample (“normalized deglycosylation activity”). Difference in the normalized and non-normalized deglycosylation activities between the RNAi silenced larvae was analyzed with one-way ANOVAs, and significant differences between the groups were determined using Tukey’s Honest Significance test.

### Mm_bGlc 17 silencing efficiency and specificity

To test for the silencing efficiency and specificity of the *Mm_bGlc17* dsRNA injection, we injected *M. melolontha* with 0.25 µg dsRNA of *Mm_bGlc17* or *GFP* dsRNA per g larva as described above. Non-injected larvae were set as controls. After injections, larvae were kept at room temperature for 2 days, after which the larvae were dissected and the individual midguts were isolated. Then, total RNA of the midgut was extracted using RNeasy Lipid Tissue Mini Kit (QIAGEN), coupled with on-column DNA digestion following the manufacturer’s instructions. One microgram of each total RNA sample was reverse transcribed with SuperScript® III Reverse Transcriptase (Invitrogen™). The RT-qPCR assay (n=7-8) was performed on the LightCycler® 96 Instrument (Roche) using the KAPA SYBR FAST qPCR Master Mix (Kapa Biosystems). The actin gene was used as an internal standard to normalize cDNA concentrations. The relative gene expressions of Mm_bGlc16, Mm_bGlc17, and Mm_bGlc18 to actin were calculated with 2^−ΔΔCt^ method. Primers (qPCR_Mm _bGlc_16_fwd, qPCR_Mm _bGlc_16_rev, qPCR_Mm _bGlc_17_fwd, qPCR_Mm_bGlc_17_rev, qPCR_Mm _bGlc_18_fwd, qPCR_Mm _bGlc_18_rev, qPCR_Mm _actin-fwd, qPCR_Mm _actin-rev) are listed in table S4.

### Effects of Mm_bGlc17 silencing on M. melolontha performance

In order to test whether *Mm_bGlc17* activity affects the performance of *M. melolontha* larvae in the presence and absence of TA-G, we assessed the growth of *Mm_bGlc17* and control (*GFP*) silenced larvae on TA-G deficient and control *T. officinale* plants. *Taracacum officinale* seeds were germinated on seedling substrate. After 15 days, plants were transplanted into 1-L rectangular pots (18×12×5 cm, length×width×height) filled with a homogenized mixture of 2/3 seedling substrate (Klasmann-Deilmann, Switzerland) and 1/3 landerde (Ricoter, Switzerland). Each pot consisted of four plants in two parallel rows of two plants which were arranged along the short edges of the pots. Rows were spaced 9 cm apart and had a distance of 4.5 cm from the short edges, and plants within each row were grown 4 cm apart from each other. After 50 days of growth, half of the pots (N=15 per genotype) were randomly selected to examine the performance of *Mm_bGlc17*-silenced larva, and the second half of the pots (N=15 per genotype) was used for *GFP*-control larva. dsRNA was synthesized as described above. Larvae were treated with 0.25 µg dsRNA of *Mm_bGlc17* or *GFP* dsRNA per g larva as previously described. One pre-weighed larva was added into a hole (4-cm depth, 1-cm diameter) in the center of the pots and covered with moist soil. After three weeks of infestation, larvae were recovered from the pots, reweighted and the midgut was extracted for subsequent RNA extraction following the above mentioned protocol. To reduce the possible effects of environmental heterogeneity within the greenhouse, the position and direction of the pots were randomly re-arranged weekly. Total RNA of the midgut was extracted using RNeasy Lipid Tissue Mini Kit (QIAGEN), coupled with on-column DNA digestion following the manufacturer’s instructions. One microgram of each total RNA sample was reverse transcribed with SuperScript® III Reverse Transcriptase (Invitrogen™). The RT-qPCR assay was performed on the LightCycler® 96 Instrument (Roche) using the KAPA SYBR FAST qPCR Master Mix (Kapa Biosystems). The actin gene was used as an internal standard to normalize cDNA concentrations. The relative gene expressions to actin were calculated with 2−ΔΔCt method.

Differences in *M. melolontha* weight gain between larval and plant RNAi treatment were analyzed with a two-way ANOVA. Differences in larval weight gain between *Mm_bGlc17*-silenced and *GFP*-control larvae were analyzed with Student’s *t*-tests for larvae grown on wild type and TA-G deficient plants separately. Differences in larval weight gain on TA-G containing and TA-G lacking *T. officinale* plants were analyzed with Student’s *t*-tests for the *Mm_bGluc17*-silenced and *GFP*-control larvae separately. A two-way ANOVA was applied to analyze differences in relative *Mm_bGluc17* expression between larval and plant RNAi treatment.

Relative *Mm_bGluc17* expression was thereto log-transformed to improve model assumptions. Differences in relative *Mm_bGluc17* expression between larvae growing on TA-G containing and TA-G lacking plants were analyzed with Kruskal-Wallis rank sum tests based on untransformed data for *Mm_bGluc17*-silenced and *GFP*-control larvae separately. The correlation between relative *Mm_bGluc17* expression and *M. melolontha* weight gain was analyzed with linear regressions for the wild type and TA-G deficient *T. officinale* plants separately. To analyze the robustness of the linear regression on wild type plants, the two values with relative *Mm_bGluc17* expression > 20 were excluded in a separate model.

To repeat the above described experiment*, T.officinale* seeds of TA-G deficient and control plants were cultivated in the greenhouse as previously described, with some slight modifications. Seedlings were germinated on seedling substrate and transplanted into individual pots (11 x 11 x 11 cm) after 21 days of growth (N=40 per line). After 70 days of growth, Larvae were treated with 0.25 µg dsRNA of *Mm_bGlc17* or *GFP* dsRNA per g larva as described above. 4 days later, for each *T. officinale* line half of the plants were infested with one pre-weighted *Mm_bGlc17*-silenced larva and the other half was infested with one pre-weighted *GFP*-control larva. After 3 weeks of infestation, larvae were carefully recaptured form the pots, weighted and added into the pots again. 5 weeks later, larvae were recaputerd again and weighted.

Differences in *M. melolontha* weight gain between larval and plant RNAi treatment were analyzed with two-way ANOVAs for three time periods (three weeks, three-eight weeks, and eight weeks after start of experiment) separately. Differences in larval weight gain between *Mm_bGlc17*-silenced and *GFP*-control larvae in these three time periods were analyzed with Student’s *t*-tests for wild type and TA-G deficient monocultures separately.

### Effects of Mm_bGlc17 silencing on deterrence of TA-G

In order to test whether *M.* melolontha glucosidase activity affects the deterrence of TA-G, we assessed the choice of Mm_bGlc17 and control (GFP) silenced larvae between TA-G deficient and control *T. officinale* plants. *Melolontha melolontha* larvae were injected with 0.025 µg*g-1 *Mm_bGlc17* or *GFP* dsRNA as described above. 1 week after ds RNA injection, the larvae were starved for three days and placed individually into the center of 250 ml plastic beakers filled with vermiculite. 5 week-old TA-G deficient and control *T. officinale* seedlings were embedded into the vermiculite-filled beaker at opposite edges with 37 replicated beakers for each of the *Mm_bGlc17* and *GFP* treatment. The feeding site was scored visually three hours after start of the experiment by inspecting the beakers from outside. Differences in the choice between TA-G deficient and control *T. officinale* plants were analyzed with binomial tests for the *Mm_bGlc17* and *GFP* silenced larvae separately.

### Data availability

Raw reads from transcriptome sequencing were deposited NCBI Sequence Read Archive (SRA) [number to be inserted at a later stage]. The data that supports the finding of the study are available from the corresponding authors upon request.

## ACKNOWLEDGEMENTS

We would like to thank Daniel Giddings-Vassão, Verena Jeschke, Nataly Wilsch, Ilham Sbaiti, Katharina Lüthy, Zohra Aziz and Ines Cambra for supporting the experimental procedures, as well as Tobias Köllner for help in the sequence analyses. The project was funded by the Max-Planck Society, the German Research Foundation (project number 422213951), the Swiss National Science Foundation (Grant No. 153517), the Seventh Framework Programme for Research and Technological Development of the European Union (FP7 MC-CIG 629134) and the University of Münster.

## AUTHOR CONTRIBUTIONS

Conceived and designed the experiment: MH, ME, SI, PR; performed the experiments: MH, TR, SG, SI, AR, JF, CP, LH, YM, WH, CAMR; analyzed data: MH, SI, CP, MR, TR, NL, ME; contributed reagents/material/analysis tools: JG, ME; wrote the paper: MH, ME, SI, JF. The authors declare that no competing interests exists.

