## Supplemental Information for "A beta-glucosidase of an insect herbivore determines both toxicity and deterrence of a dandelion defense metabolite"

SUPPLEMENTARY INFORMATION

### Supplemental texts

### SI Text

**Text S1.** Alignment of β-glucosidases of *M. melolontha* and other insect species. The catalytic glutamates of the glucosidase-characteristic NEP and ITENG motifs are indicated with red triangles.

```
Mm_bGlc1      -MKHL-LLFLIL-------------AVFANAQNY-----S--FPDGFLFGVSTASYQVEGAWNISGKGENIWDRLVHSTPERVVDMSNGDVACDAYHKTE  78
Mm_bGlc11     -MKVQ-VVLILC-------------IYLCNAQEY-----E--FPEGFIFGVGTASYQIEGAWNESDKGENIWDYLTHNYPEFVMDSSNGDVASDGYHKLD  78
Mm_bGlc14     -MRRI-IFLLLL-------------GCLCKAQDG-----DRLFPEDFKFGVATASYQIEGAWQADGKGENIWDYLTHNKDTLIDDGSNGDVACNSYNKVD  80
Mm_bGlc15     --MKL-VIFALI-------------LTYVSAQDT-----R--FPDGFKFGVATASYQIEGAWDANGKGENIWDRLTHTRPELIADGSNGDIACDAYHKTE  77
Mm_bGlc16     -MKVL-VILCTV-------------IYLCKAQAY-----E--FPDDFLFGVATAAYQIEGAWDTNGKGENIWDNLTHTYPHLVVDGSNGDVACDAYSHTA  78
Mm_bGlc17     -MKRL-ILIFVL-------------ACLAKAQDY-----Q--FPEGFKFGVATASYQVEGAWNENGKGENIWDRLTHTQPDLIADNSNGDIACDAYHKTE  78
Mm_bGlc18     -MGYFEPLIVFL-------------ISLWRAQSY-----V--FPEEFLFGVATSTYQVEGAWNLSGKGVNIWDHLTHTNPEFTADGSNGDVACDAYHKTK  79
Mm_bGlc19v    -MSSY-VYKNIIETKTIDEDFVPEGFAACSSSEY-----T--FPDGFIFGVATSAYQVEGAWEDDGKGENIWDHLTHTRPTAITDESNGDIACDTYHKTV  91
Mm_bGlc2      -MKPIGLIVLVF-------------LAAADAQNY-----S--FPDDFIFGVATASYQVEGAWNLNGKGENIWDRLTHSTPELIADNSNGDVACDTYHKTE  79
Mm_bGlc3      -MKSL-ILILVL-------------AVSVNGQIY-----S--FPEDFQFGVATASYQVEGAWNVSGKGENIWDRLTHSTPERIVDNSNGDVACDAYHKTA  78
Mm_bGlc5      -MKAI-IILALL-------------CSVTKAQDY-----E--FPQDFLFGVATASYQIEGAWNESDKGENIWDYLTHNDPDFVTNRDNGDDACKAYYKTD  78
Mm_bGlc6      -MRRV-LILILL-------------ACLCAAQDEGEETDNTVFPDDFQFGVATAAYQTEGGWDDGGKGENIWDYLLHNTDNFTVDGSSGDDACKSYYKTE  85
B. brassicae  ---------------------------------MDY-----K--FPKDFMFGTSTASYQIEGGWNEDGKGENIWDRLVHTSPEVIKDGTNGDIACDSYHKYK  62
C. populi     -MRTV-FLIVLT-------------SYGVFGHNG-----T--FPPNFLFGVAHSAYQVEGAWNIDGKKDSIWDHFVHQNRSRIHNRDTGDVACDSYDLYK  78
P. striolata  MQQTIAFVLVLL-------------QFAFNADGALKNNGQ--FPRNFLFGAATASYQIEGAWNEDGKGQSTWDEFTHRIPSPVTNNDTGDIACDHYHRYK  85
T. molitor    -ARGA-ILLVICAST----------ITLADVPDY-----Y--FPDGFVFGAATAAYQVEGGWDEDGKGESIWDRGTHEHADWVADNSNGDIACDSYHKYK  81

Mm_bGlc1      KDVQLLKNLNVNFYRFSISWSRILPSGYVNVINPDGIRYYNELIDELLANGIEPFVTMYHWDLPQPLQ-EIG-GWANPLISDIFADYAELLYSQFGDRVK  176
Mm_bGlc11     QDIQLMKNVGVDFYRFSLSWSRILPTGHINKVSDDGVRYYDELIDKLLANDILPYVTIFHWDLPQPLL-EIG-GWPNPALADIFANYADLVFRLFGDRVK  176
Mm_bGlc14     DDVELLVELRVQYYRFSISWSRILPTGHANEVNEAGVQYYSDLIDKLLENGITPLVTMYHWDLPQPLQ-EIG-GWTSPVLVDLFVDYANVLFTSFGDRVT  178
Mm_bGlc15     EDVQLLKGLGVDFYRFSISWSRILPTGYADEINEDGIRYYNELIDELLANDITPYVTMYHWDLPQPLQ-EIG-GWLNSSLADIFVDYADVLYEQFGDRVK  175
Mm_bGlc16     EDVQLLKNLGVDFYRFSFSWSRVLPTGKTDYINPDGIRYYNELIDTLLENNIEPMATMYHFDLPQPLE-DEG-GFLNIVIADYFEDYAEVLYENFGDRIK  176
Mm_bGlc17     EDVQLLKNLGVNFYRFSISWSRILPTGYTNVVNEDGIRYYNALIDALLENGITPLVTMFHWDLPQPLQ-EIG-GWPNPLLADIFADYADILYREFGDRVK  176
Mm_bGlc18     EDVQLMVDMGLDVYRFSLSWTRILPNGYNNYTNPDGVRYYNELIDTLLANGIKPLVTMFHWDIPQVFQDDYG-GWLGSDMVDIFVDYADVAFSLFGDRVK  178
Mm_bGlc19v    EDVQLLNTLGVDFYRFSLSWSRLLPTGHANTVNPLGVAYYNELIDELIANDITPLVTLFHWDLPQPLQ-EIG-GWPNPLLVDLYADYANIVFSLFGDRVK  189
Mm_bGlc2      DDVQLLNNIGVNFYRFSISWSRILPTGYVNEINSDGIRYYNELIDELLANDIQPLVTMFHWDLPQPLQ-EIG-GWTTPLISDLYADYADILYSEFGDRVK  177
Mm_bGlc3      EDVQLLKDIGVDFYRFSISWSRILPTGYVNEINPDGIRYYNELIDELLANNIVPLVTMFHWDLPQSLQ-DIG-GWPNPLISDLFADYAEILYSQFGDRVA  176
Mm_bGlc5      EDVQMIKDVGAHFYRFSLSWSRLLPTGHINKVSEDGVAYYNDLIDQLLANDIIPYVTIFHWDLPQPLL-ELG-GWPNPALADIYAAYANFVFNEFGDRVQ  176
Mm_bGlc6      EDVELLKNIGVNFYRFSISWSRILPTGHSYSVNQVGVNYYNDLINRLVANGIEPMVTMYHWDLPQPMQ-ELG-GWPNPRLAEYFVDYADVLFRLFGDRVK  183
B. brassicae  EDVAIIKDLNLKFYRFSISWARIAPSGVMNSLEPKGIAYYNNLINELIKNDIIPLVTMYHWDLPQYLQ-DLG-GWVNPIMSDYFKEYARVLFTYFGDRVK  160
C. populi     EDIRLAAALGVNMYRFSISWPRVLPNGDTTVINKRGLRYYERILNEVRRHDMVPVVTIYHWDLPQALM-KRGIDWTNPKLVNYFVDFARIVIRHL-RHVG  176
P. striolata  EDIKMAADLGLQAYRFSISWPRVLPNGYADTINEKGMQFYKDLVDEIVKYGMVPVCTLFHWDLPLKLY-ENGIDWRNEKIIDIFVAYSRLMIQNL-PKVG  183
T. molitor    EDVQMLKTLGVNFYRFSIAWSRVLPTGKADEVNQAGIDYYNNLIDELLANDIEPYVTMFHWDLPQPLQ-DEG-GWPDRKLADYFVDYARVLFENFGDRIK  179

Mm_bGlc1      NWITFNEPPVVC-SGYA-GIMPPGYDQGGIGDYLCAHHLLISHGKAYRIYDEEYRNAQQGRVGITIDTSWYEP-ESDSDLDI--SERVLQMKYGLYVHPI  271
Mm_bGlc11     NWITINEPYQICEEGFSEGILAPGYAQHGIGGYLCGHTVLLAHAKAFKIYNDKYREEQDGRVGIVVHGAWAEP-SSESEEDKSAAETWQQMNFGWFLHPI  275
Mm_bGlc14     NWITFNEPYQICQQGYSEGTKAPAYTQDGVGGYLCGHTLLLAHAYTYHLYHDDYAGT-DGRIGITVNGVWAEA-ASEDSVDQTAADDYIEFHFGWFFNPI  276
Mm_bGlc15     DWITFNEPIQVCEAGYSNAGKAPAYTMAGVGGYLCGHTLLIAHGKTYRLYNEKYRESQQGRVGITIDAGWYEP-VSDSDTDIEAAERSMQMNYGWYAHPI  274
Mm_bGlc16     QWITFNEPSPICENGYGADSMAPLANQPGVGGYICSRTLLVAHAKAYHLYNDRYKGSQGGRVGITINNFWYEP-YADTTD--SAVELALQVAFGWFTHPI  273
Mm_bGlc17     DWLTFNEPTPICIGGYSEGWMAPAYELQGVGGYLCGHTLLIAHGKAYRLYNEKYRDTQEGRVGITIDSGWYEP-ASESEEDIQAAEDSVHIKYGWMVHPI  275
Mm_bGlc18     DWITFNELHIFCELGHSMDIMAPALGLSGVGGYLCAHNILIAHGKTYKLYNEKYRDIQKGRIGITIDGEWKEP-ASTCPEDIETAERALQMEFGWIAHPI  277
Mm_bGlc19v    DWITFNEPYQICQEAYSEANKAPAYNQDGTGGYLCAYTVLLAHARAYRLYETTYKEAQQGRVGITVHGIWAEP-ETATDADIAAAESYQQFHFGIYLHPI  288
Mm_bGlc2      NWITFNEPNAVC-AGYA-GSMAPGYYIEGIGDYLCAHHLLISHGKAYQIYNEKYRDAQQGIVGITVYTPWYEP-ASDSEDDIQAAERAMQMTYGIYVHPI  274
Mm_bGlc3      NWITFNEPPVIC-GGYA-GGLAPGYNQDGIGDYLCAHNLLISHARAYRIYEEKYKDVQEGRVGITVNTRWYEP-ASDSAEDAAAAERAIQMIYGIYIHPI  273
Mm_bGlc5      NWITFNEPYQICEQGFSDGSLAPGYAQQGIGGYLCGHTVLLAHSKAYHIYDDLYRETQGGRVGIVVHGAWGEP-ESDSDEDAQAAENFIQMNFGWFLHPI  275
Mm_bGlc6      TWVTINEPYEFCHRGYGTGGLAPGYTQDGIGDYLCAYTVLLAHARTYRMYQENYLEDQGGRIGISLNSDWYEG-ADVNAEDTAAAETSNQMMLGWFAHPI  282
B. brassicae  WWITFNEPIAVC-KGYSIKAYAPNLNLKTTGHYLAGHTQLIAHGKAYRLYEEMFKPTQNGKISISISGVFFMPKNAESDDDIETAERANQFERGWFGHPV  259
C. populi     YWVTINEPHTFCRLGYGLGIFAPGIESSGRDEYKCIYTMMKAHAAVYRMYKREFPHY-KGKMSLVYDMQWYLPKNASSAKDVEAVRRGLDFDAGLYVHPV  275
P. striolata  YWSTINEPRVHCLRSYGDGKHAPGIAESGIADYQCSYVILKAHAKAYRMYKKEFPHYKAPF-GIVIDCQWYVP-ATNSSEDIAAAERYYEFECGMYFHPI  281
T. molitor    YWMTFNEIMQICEAGYSGGSFAPYISNPGVGGYECTHTVLLAHGRTYRLYDSDFRAEQNGQIGIAIDSYWHEPNYADRETDQEASEVDMQLNYGWFVNPF  279

Mm_bGlc1      FSETGDYPPILRKRVDDLSVEQGYARSRLPHFTPDEIEFLHGSSDFLGLNHYTSYLCSSSSYD------LLPPSQWADTGAVCYQSDEW-ESSSSTWLKV  364
Mm_bGlc11     YSESGDYPSIMKERIEDLSELEGFPRSRLPVFSQEEIELLKDSSDFLGLNHYGSFLCKPLEES------QIIPSHENDIGSECYLSDDW-EPSASPWFSV  368
Mm_bGlc14     Y--IGDYPQIMKDNIQARSELEGFPDTRLPEFTETEIALLRSSSDFLGLNHYTTYQCTPLEDV----DSLEQPSFKVDSGVNCWSPTDW-EGGASSWLKV  369
Mm_bGlc15     FSETGDYPEIMKERVAELSELEGFETSRLPSFTQEEIEFIRGSSDFLGLNHYSSSLCTSIPEE----WALTGPSQWIDVGSLCLPSPEW-EVAASSWLYV  369
Mm_bGlc16     YSKEGDFPPAMKERIARLSEEEGFSASRLPELTTEEIELIKGSSDFFGLNHYTTHFCSESGID-----SISRPSHNYDMGVTCMPDYSY-EVAGSFWLSV  367
Mm_bGlc17     YSETGDYPPVLRERVDALSAEEGFARSRLPIFTEEEIELIKGSSDFLGLNHYTTNLCTPIPEE----WGVVGPSHYVDSGANCYQDPSW-EGSGSSWLKV  370
Mm_bGlc18     FSESGDYPPVMRSRIDVMSAEEGLKTSRLPYFSKEEIELIRGSADFLGLNHYTASLCSSEFKE-----LPSRPSYTSDTGANCYQPDYW-EPTGVSQFKV  371
Mm_bGlc19v    FSEEGDFPEIVKTRVKQISDVIGYSNNRLPVFTAEEIEYIHGTSDFLGFNHYSTDLCKAADDA-----SLVHPSNKGDTGADCQKSDDW-ESAASSWVKV  382
Mm_bGlc2      YSETGDYPPVLRERVDALSAEQGYARSRLPYFTEEEIELIRGSFDFLGLNHYTTYLCYAATYD------LQPPSQWGDIGAAYYQLDEW-EGSGSSWLKV  367
Mm_bGlc3      YSETGDYPPILRERVDALSAEQGYARSRLPYFTEEEIELIRGSSDFLGLNHYTSLLCSFSTYD------LSPPSQWADTGVACYQSDEW-EGSGSSWLKV  366
Mm_bGlc5      YSDAGGYPPSMVSAIHALSTEEGFPRSRLPPFTEEEIANLKDTSDFLGLNHYGAYLCRPLSET----DEVISPSHVKDIGTYYYLSEDW-DQSASDWFAV  370
Mm_bGlc6      YHEDGDWPEIMKERIYQLSMDAGFVRSRLPTFTSDEIEEIKGSYDFFGLNHYTTLICTPTDSDSNEIDAEIVTSYENDVGTSCTVNENW-DETANGW-RV  380
B. brassicae  Y--KGDYPPIMKKWVDQKSKEEGLPWSKLPKFTKDEIKLLKGTADFYALNHYSSRLVTFGSDP--------NPNFNPDASYVTSVDEAWLKPNETPYIIP  349
C. populi     F--LGGWPPEAEKRIAERSRLANLTTSRLPAFSPEEIAFINGTYDFFGVNIYTGLLVENLPEY-----NKSISSYEADVHLELSFDPNW--TLEAGYFAF  366
P. striolata  S--KGDWPDIVKKRIAHRSQLEGRSSSRLPQITPEDIRLMKGAQDYIAINHYFTIMAANAEEA-----SYDEINYKHDVRVNLFNDPSW-KISVLGW-AI  372
T. molitor    I--NGNYPEVMIERVKANSLAEGYPQSRLPEFTADEQEMMKGTFDFLGLNHYSSDKVYFAEDG-----AGDHPSHWADTGVIGYQDASW-PGSASSWLKV  371

Mm_bGlc1      VPWGFRKLLNWIKNEYNNPEVIITENGFSDSP--GDVNDCRRVNYYNQYLESLLQAIHEDDCYVTGYTAWSLMDNFEWLMGYTARFGLYMVDF-DDEDRP  461
Mm_bGlc11     TPWGLRNILNWIKEEYGNPEVIITESGFTDSS--NDTRDCGRVNYYNTYLESLLEAIHEDECNVSGYTAWSIIDLFEWQFGYTYRFGMHYVDF-DDPDRP  465
Mm_bGlc14     HPSGLTSILNWIREKYDNPEVIITENGFSEAGEVNDLNDCDRVSYYNGYLYAVLDAL-EDGCQVSGYMAWSLMDNFEWMRGYSERFGLYYVDF-EDEDRP  467
Mm_bGlc15     VPWGLRRLLNWIKNEYGNPEVIITENGFSDT---GELNDCRRVNYYNSYLTAVLEAILEDGCNVSGYTAWSFMDNFEWLMGYTERFGIHYVDF-EDPDRP  465
Mm_bGlc16     IPWGLRKMLNWIKEEYNNPEIIITENGYSDNE--VILNDCRRINYYNSYLTELLNAIYEDNCNVTGYTAWSFMDNFEWRMGYSEKFGLYSVDF-NDPDRT  464
Mm_bGlc17     VPWGLRRLLNWIKENYDNPEVLITENGVSDNT--GILNDCRRINFYNTYLTAVLEAIHVDGCNVVGYTAWSFMDNFEWMQGYTERFGLYHVDF-NDTDRT  467
Mm_bGlc18     TPWAFGKLLVWIKNEYNNPEVIVTENGYSDNT--GDLYDCRRVYYYNSYLTELLHVVNDEGCRITGYMAWSFMDNFEWGNGYTAKFGIVNVDF-NDADRP  468
```

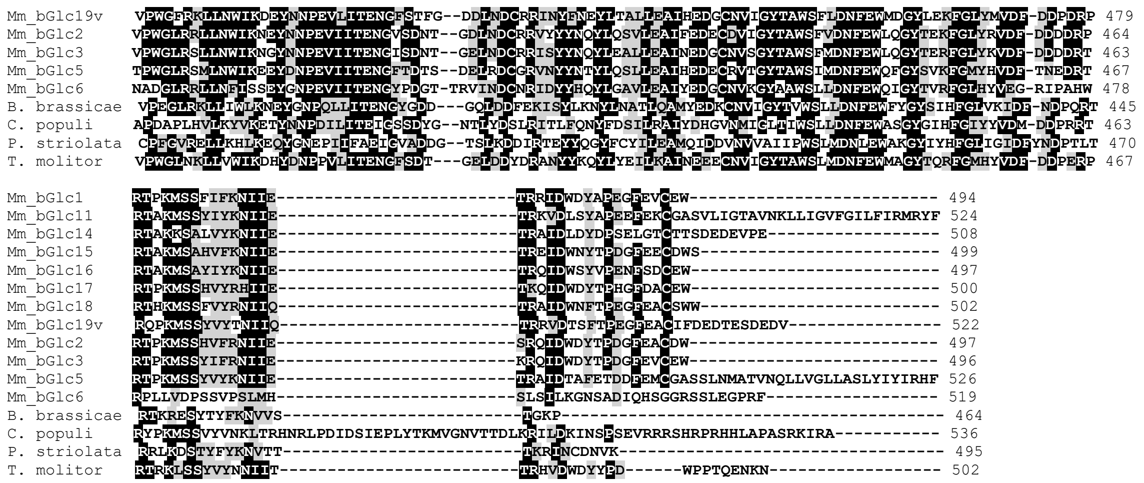

**Text S2.**

*>Mm_bGlc1*

ATGAAGCATCTACTATTATTTTTAATTTTGGCGGTTTTTGCAAATGCGCAGAATTATTCATTCCCTGATGGTTTTCTATT
TGGTGTTTCTACAGCATCGTACCAGGTTGAGGGAGCTTGGAATATAAGTGGTAAGGGAGAAAATATATGGGACAGACTCG
TACATTCAACTCCAGAACGTGTAGTAGATATGAGTAATGGAGACGTAGCTTGCGATGCTTATCATAAAACAGAAAAAGAC
GTCCAGCTACTGAAAAACCTCAACGTAAACTTTTACAGATTTTCCATCTCCTGGTCCAGAATCCTCCCATCAGGTTACGT
TAACGTAATTAACCCCGATGGCATTCGATATTACAACGAATTAATTGACGAACTGTTGGCGAATGGTATTGAGCCCTTTG
TAACGATGTACCACTGGGATTTGCCCCAACCCCTGCAAGAAATCGGTGGCTGGGCTAATCCCTTAATATCCGATATATTT
GCTGATTATGCTGAACTTCTTTACTCGCAGTTCGGTGATAGGGTGAAAAATTGGATAACGTTTAACGAACCTCCTGTTGT
GTGTAGTGGGTATGCTGGAATTATGCCACCTGGGTATGATCAAGGAGGTATCGGTGATTATTTATGTGCACACCATCTGC
TTATTTCACACGGAAAGGCATATCGAATTTATGACGAAGAATACAGGAATGCTCAACAAGGCAGAGTTGGTATAACTATA
GATACTAGTTGGTATGAACCTGAATCAGATTCAGATTTGGACATTTCTGAAAGGGTACTGCAAATGAAGTATGGTCTATA
CGTACATCCAATTTTTTTCCGAAACAGGGGATTATCCGCCGATACTAAGAAAAAGAGTGGACGATTTAAGTGTAGAACAAG
GTTACGCTCGTTCCAGACTACCTCATTTTACTCCAGATGAAATTGAATTCCTTCATGGTTCATCCGATTTCCTTGGTTTG
AACCATTACACATCTTATCTATGTTCTTCTTCTTCATACGATTTGTTACCTCCATCACAGTGGGCCGATACTGGTGCCGT
TTGTTATCAGTCAGACGAGTGGGAAAGTTCGTCTTCTACATGGTTGAAAGTCGTACCTTGGGGCTTTAGAAAGTTGCTAA
ACTGGATAAAAAATGAATACAACAATCCTGAAGTAATAATAACAGAAAACGGATTTTCTGATTCTCCTGGGGATGTTAAT
GATTGCAGAAGAGTTAATTATTATAATCAATACCTAGAAAGCCTGTTACAAGCAATTCATGAAGACGACTGTTACGTTAC

GGGTTACACAGCTTGGAGTTTAATGGATAATTTCGAATGGCTAATGGGTTATACCGCGAGATTTGGACTCTACATGGTCG
ATTTCGATGACGAAGACAGACCTAGAACGCCAAAAATGTCTTCGTTTATCTTCAAAAATATTATCGAAACTAGGCGAATT
GATTGGGATTACGCGCCGGAAGGATTCGAAGTTTGTGAATGG

> *Mm_bGlc2*

ATGAAACCAATCGGTTTAATTGTTTTAGTTTTCTTGGCCGCTGCCGACGCACAAAATTATTCCTTTCCCGATGATTTTAT
ATTCGGTGTTGCTACAGCGTCATATCAAGTTGAGGGAGCTTGGAATCTAAATGGTAAAGGAGAAAATATATGGGATAGAT
TGACGCATTCAACTCCAGAACTTATAGCAGATAACAGCAATGGAGACGTAGCCTGTGATACGTATCATAAAACTGAGGAT
GATGTGCAACTACTTAATAACATCGGAGTAAATTTCTACAGATTTTCCATATCGTGGTCCAGAATTCTTCCAACAGGTTA
CGTAAATGAAATTAATTCAGACGGAATTAGATATTATAACGAGTTAATCGATGAACTCTTGGCCAATGATATCCAGCCTC
TTGTTACGATGTTCCACTGGGATTTGCCCCAGCCTTTGCAGGAAATTGGTGGATGGACAACTCCTTTGATATCCGATTTA
TACGCTGATTATGCTGATATTCTTTATTCAGAGTTCGGTGATAGAGTGAAGAATTGGATAACCTTCAATGAGCCTAATGC
TGTTTGCGCTGGTTATGCTGGAAGCATGGCACCAGGGTATTATATAGAAGGCATAGGTGACTACTTATGCGCACATCATT
TGCTCATTTCTCATGGGAAAGCTTACCAAATTTACAATGAGAAATATAGGGATGCCCAACAAGGTATAGTTGGAATAACT
GTGTATACTCCATGGTATGAACCAGCTTCAGATTCAGAAGACGATATTCAAGCTGCTGAAAGAGCTATGCAAATGACGTA
TGGCATATACGTACATCCAATTTATTCAGAAACAGGTGATTATCCACCAGTATTGAGGGAAAGAGTAGATGCTTTAAGCG
CGGAACAAGGTTATGCGAGATCTAGATTGCCCTATTTTACCGAAGAGGAAATAGAACTCATTCGTGGCTCGTTTGATTTT
CTGGGTCTAAATCACTACACGACTTATTTATGCTATGCTGCCACTTACGATTTGCAACCTCCGTCACAATGGGGAGATAT
TGGTGCTGCCTACTACCAGTTAGATGAATGGGAAGGTTCTGGTTCTTCTTGGTTGAAGGTCGTACCTTGGGGTCTGAGAC
GTTTATTAAATTGGATTAAAAATGAATACAACAATCCCGAAGTGATAATAACAGAAAACGGAGTTTCTGATAACACTGGG
GATTTGAATGATTGCAGAAGAGTTTATTATTACAATCAATACTTGCAATCCGTTCTAGAAGCAATTTTTGAAGACGAATG
TGATGTAATAGGTTATACAGCTTGGAGTTTCGTAGACAATTTTGAATGGTTACAAGGTTATACTGAAAAATTCGGTCTTT
ATAGAGTTGATTTTGACGATGATGACAGGCCTAGAACTCCGAAAATGTCTTCTCATGTCTTCAGGAATATTATCGAAAGT
AGACAGATTGACTGGGATTACACACCAGATGGATTTGAAGCTTGCGATTGG

> *Mm_bGlc3*

ATGAAATCACTAATTTTAATTTTAGTTTTGGCGGTCTCAGTCAATGGGCAAATATACTCGTTTCCCGAAGACTTTCAATT
CGGCGTCGCTACAGCGTCCTATCAAGTTGAGGGAGCTTGGAATGTAAGCGGTAAAGGAGAAAACATATGGGACAGGTTAA
CACATTCAACTCCAGAACGTATCGTAGATAACAGTAATGGTGACGTAGCTTGTGATGCTTACCATAAAACCGCAGAAGAC
GTGCAACTGCTCAAAGACATCGGAGTCGATTTCTACAGATTTTCAATTTCATGGTCCAGAATCCTTCCAACCGGTTACGT
AAACGAAATTAATCCCGATGGCATTCGATATTACAACGAACTAATTGACGAACTATTGGCTAATAATATCGTACCTCTAG

TTACGATGTTCCATTGGGACTTGCCTCAGTCTTTGCAGGATATCGGTGGATGGCCAAATCCATTAATATCTGACCTATTCGCCGATTATGCTGAAATTCTTTACTCACAGTTTGGCGACAGGGTGGCGAACTGGATAACGTTCAATGAACCTCCTGTTATTTGTGGTGGTTACGCGGGGGGTTTAGCGCCCGGGTATAATCAAGATGGCATAGGCGATTATTTATGCGCACACAATCTCCTAATCTCACACGCAAGAGCGTATCGAATTTATGAAGAGAAATATAAGGACGTTCAAGAAGGTAGAGTTGGTATTACTGTGAATACTAGATGGTACGAACCGGCTTCAGATTCAGCAGAAGATGCCGCAGCTGCCGAGAGAGCTATACAAATGATTTATGGTATATACATACATCCAATTTATTCAGAAACAGGAGATTATCCACCAATATTAAGAGAAAGAGTAGATGCATTGAGTGCAGAACAAGGTTACGCGAGATCTAGACTACCTTATTTTACAGAAGAAGAAATCGAACTCATTCGGGGTTCGTCCGACTTTTTGGGTTTAAATCATTACACCAGTCTCTTATGTTCCTTTTCAACATACGATTTAAGTCCTCCATCTCAGTGGGCTGATACGGGTGTAGCTTGTTACCAATCAGATGAGTGGGAAGGATCTGGTTCCTCTTGGTTAAAGGTTGTACCTTGGGGTCTGAGAAGTCTCCTAAATTGGATCAAAAACGGATATAACAATCCCGAAGTCATAATAACGGAAAACGGAATTTCTGATAATACTGGAGAACTAAACGATTGTAGACGAATCAGCTATTATAATCAATATTTGGAAGCACTTTTGGAAGCTATTAATGAAGACGGATGTAATGTTAGTGGATACACAGCTTGGAGTTTCATGGATAATTTCGAATGGCTGCAGGGCTATACTGAAAGATTTGGACTTTACAAAGTTGATTTCGATGATGATGATAGAACTAGAACTCCAAAAATGTCTTCTTACATCTTCAGGAATATTATTGAGAAAAGGCAAATCGACTGGGACTATACACCGGATGGATTCGAAGTCTGCGAATGG

> *Mm_bGlc5*

ATGAAAGCGATTATTATATTGGCTTTACTGTGTTCTGTAACCAAAGCGCAAGATTACGAATTTCCCCAAGACTTTCTTTTTGGTGTTGCTACAGCGTCTTATCAAATAGAAGGTGCTTGGAATGAAAGTGATAAAGGCGAAAATATATGGGATTATCTAACTCATAACGATCCAGATTTCGTGACTAATAGAGACAATGGTGATGACGCCTGTAAAGCTTACTATAAAACAGATGAGGATGTCCAAATGATTAAGGATGTAGGCGCTCATTTCTATAGATTTTCGCTGTCTTGGTCGAGACTCCTTCCAACAGGACACATTAATAAAGTCAGCGAAGATGGTGTTGCATATTATAACGATTTAATTGATCAATTATTAGCGAACGACATAATTCCATATGTGACAATATTTCATTGGGATTTACCCCAGCCTTTGCTGGAGTTGGGCGGATGGCCTAATCCAGCTTTGGCTGATATCTATGCAGCTTACGCTAACTTCGTATTCAACGAATTTGGAGATAGAGTACAAAATTGGATTACCTTCAATGAACCGTACCAAATTTGTGAACAGGGTTTTTCGGACGGTAGCTTAGCACCTGGATATGCTCAACAAGGAATTGGCGGTTATTTATGTGGTCATACTGTTCTCCTAGCTCATTCGAAAGCTTATCATATCTATGACGACCTCTATAGGGAAACACAAGGAGGTAGAGTCGGTATAGTCGTTCACGGTGCTTGGGGTGAACCAGAATCTGATTCAGATGAAGATGCACAAGCAGCTGAGAATTTCATTCAAATGAATTTCGGTTGGTTCTTACATCCAATTTACAGTGACGCTGGTGGATATCCACCCAGTATGGTTTCCGCTATACATGCACTAAGTACCGAAGAAGGGTTTCCCAGATCAAGATTACCGCCTTTTACAGAAGAAGAGATCGCCAACCTAAAGGATACGTCGGATTTTCTGGGATTAAATCATTACGGAGCTTATTTATGCAGACCCCTCAGTGAGACAGACGAAGTGATATCACCTTCACATGTTAAGGATATCGGAACTTATTATTATCTGTCCGAAGATTGGGATCAGAGTGCTTCGGATTGGTTTGCGGTGACACCTTGGG

GTTTGAGGAGCATGTTAAATTGGATCAAAGAAGAGTACGACAATCCCGAAGTAATCATTACAGAAAATGGTTTTACGGATACTAGTGACGAGTTGCGTGATTGCGGTAGAGTTAATTACTATAATACATACTTGCAATCACTTCTGGAAGCCATACACGAAGATGAATGCAGAGTTACCGGTTACACTGCTTGGAGTATCATGGACAACTTCGAATGGCAATTTGGTTACTCGGTGAAATTTGGAATGTACCACGTTGATTTCACCAATGAAGATAGGACCAGAACACCAAAAATGTCGTCATACGTATACAAGAATATAATTGAAACAAGGGCAATCGATACCGCCTTCGAAACTGACGATTTTGAAATGTGTGGGGCGTCCTCGTTGAATATGGCCACAGTAAATCAATTACTTGTAGGTCTTCTTGCCAGTTTGTACATTTATATTAGGCATTTT

> *Mm_bGlc6*

ATGAGACGCGTCTTAATATTAATCTTATTAGCGTGCCTGTGTGCGGCTCAAGATGAAGGGGAAGAAACAGATAACACTGTGTTCCCAGATGACTTTCAGTTCGGTGTTGCCACAGCTGCTTATCAAACAGAAGGTGGTTGGGATGACGGTGGGAAAGGTGAAAATATTTGGGACTACTTGCTCCACAATACCGATAATTTTACTGTCGATGGAAGTAGTGGCGATGATGCGTGCAAAAGTTATTATAAAACTGAGGAAGATGTCGAATTATTGAAGAACATTGGTGTAAATTTCTATCGGTTCTCAATCTCGTGGTCTAGAATTCTGCCGACTGGACATTCGTACAGCGTAAATCAAGTAGGCGTCAACTACTATAATGACTTGATTAACCGACTAGTAGCTAATGGAATAGAACCAATGGTTACGATGTACCATTGGGATTTACCGCAACCCATGCAAGAATTGGGTGGTTGGCCCAACCCCCGTTTGGCAGAATATTTCGTAGATTATGCCGATGTACTCTTCAGACTTTTCGGAGACAGAGTCAAAACTTGGGTCACTATCAACGAACCGTATGAATTTTGCCATCGGGGATATGGAACGGGCGGGTTAGCACCTGGATACACGCAAGATGGAATTGGAGATTATTTATGTGCTTATACTGTCCTTCTAGCTCATGCTAGAACTTATAGAATGTACCAGGAGAATTACTTAGAAGATCAAGGAGGAAGAATCGGAATATCCCTCAACAGTGATTGGTATGAAGGTGCAGATGTTAATGCTGAAGATACGGCGGCAGCTGAAACCAGTAACCAAATGATGTTGGGCTGGTTCGCACACCCGATTTACCATGAAGATGGTGACTGGCCTGAAATAATGAAAGAGAGAATTTATCAATTAAGTATGGACGCAGGATTTGTGAGAAGTCGACTTCCCACTTTCACTTCAGACGAAATAGAAGAAATAAAAGGCAGTTATGACTTCTTTGGCCTTAATCACTATACCACACTTATTTGTACTCCAACTGATAGTGATAGTAACGAAATTGATGCCGAAATTGTGACTTCATATGAAAATGATGTTGGTACATCCTGTACTGTAAACGAGAATTGGGATGAAACGGCTAATGGATGGAGAGTCAATGCTGATGGTTTGCGAAGATTGTTGAATTTTATTAGTAGCGAGTACGGTAACCCGGAAGTAATCATAACGGAAAATGGATACCCCGACGGTACAACTCGCGTTATAAATGACTGCAATAGAATTGATTATTACCACCAATACCTAGGAGCAGTACTGGAAGCAATATACGAGGATGGATGTAATGTTAAAGGATATGCAGCTTGGAGTTTATTGGATAACTTTGAATGGCAAATTGGCTATACGGTGAGGTTTGGTCTCCATTACGTAGAAGGGCGAATTCCAGCACACTGGCGGCCGTTACTAGTGGATCCGAGCTCGGTACCAAGCTTGATGCATAGCTTGAGTATTCTAAAGGGCAATTCTGCAGATATCCAGCACAGTGGCGGCCGCTCGAGTCTAGAGGGCCCGCGGTTCTAA

> *Mm_bGlc11*

ATGAAAGTGCAGGTTGTATTAATTTTATGCATTTATTTATGTAATGCGCAAGAGTATGAATTCCCCGAAGGTTTTATATT

CGGCGTAGGTACCGCGTCATATCAAATAGAAGGTGCTTGGAATGAAAGCGATAAAGGAGAAAATATATGGGATTACCTAA
CTCATAACTATCCTGAATTTGTAATGGATAGCAGTAACGGTGACGTGGCAAGTGATGGTTATCACAAGCTAGATCAAGAT
ATCCAATTAATGAAGAATGTTGGTGTAGATTTCTATAGGTTTTCGCTTTCATGGTCGAGGATCCTTCCCACCGGTCACAT
AAATAAAGTCAGCGATGATGGTGTTCGCTATTACGATGAGCTAATCGATAAATTATTAGCGAATGACATACTTCCATACG
TAACGATATTTCATTGGGACTTACCTCAGCCCTTGTTGGAGATTGGTGGATGGCCTAACCCTGCTTTAGCCGACATTTTC
GCAAATTATGCCGATCTAGTGTTTCGTTTATTTGGAGATCGTGTTAAAAATTGGATCACTATTAACGAGCCGTATCAGAT
TTGCGAGGAAGGATTTTCAGAAGGGATATTGGCACCTGGTTATGCTCAACATGGAATCGGAGGTTACTTGTGCGGTCATA
CGGTATTGCTAGCCCATGCTAAAGCGTTTAAGATTTACAACGACAAGTACAGAGAGGAACAAGATGGCAGAGTTGGGATA
GTAGTTCATGGTGCGTGGGCTGAACCAAGTTCGGAATCAGAAGAGGACAAATCAGCTGCTGAAACATGGCAACAAATGAA
TTTTGGATGGTTTTTGCATCCAATTTACTCAGAATCTGGAGATTATCCATCAATTATGAAAGAAAGAATAGAAGATCTAA
GCGAATTAGAAGGATTTCCGCGATCCAGATTACCTGTATTCAGTCAGGAAGAGATCGAATTGCTAAAAGATTCTTCCGAT
TTCTTAGGTCTTAATCACTATGGATCCTTTCTATGTAAACCACTGGAGGAATCACAGATTATTCCATCACATGAGAATGA
TATTGGCAGCGAATGTTATTTATCGGATGATTGGGAGCCGAGCGCATCTCCATGGTTTAGCGTGACTCCGTGGGGTTTGA
GGAACATTCTAAATTGGATTAAGGAAGAATATGGTAATCCAGAAGTGATAATAACAGAAAGCGGTTTTACAGATTCGAGT
AATGACACACGTGATTGTGGAAGGGTTAATTATTATAACACTTACTTGGAATCATTGTTGGAAGCCATACACGAAGACGA
GTGCAACGTAAGCGGTTACACTGCTTGGAGCATTATCGATCTTTTTGAATGGCAATTTGGTTACACATATCGTTTTGGTA
TGCATTATGTGGATTTTGACGACCCAGATAGACCTAGAACGGCAAAAATGTCTTCCTATATTTACAAGAACATCATCGAA
ACGAGGAAAGTTGATTTGAGCTATGCCCCGGAAGAATTCGAAAAATGTGGAGCATCAGTTTTAATTGGAACGGCGGTGAA
CAAATTACTTATTGGCGTCTTTGGGATATTATTTATTCGTATGAGATATTTT

> *Mm_bGlc14*

ATGAGACGAATCATTTTCCTTTTGCTTCTAGGATGCTTATGCAAGGCACAGGATGGAGACCGCTTGTTTCCAGAAGACTT
TAAATTTGGCGTTGCCACAGCGTCTTATCAGATAGAAGGAGCATGGCAGGCTGACGGTAAAGGAGAGAATATCTGGGACT
ATCTAACCCACAACAAAGACACATTAATAGACGACGGCAGTAATGGCGACGTTGCATGCAATTCTTATAACAAAGTTGAT
GACGACGTGGAATTATTAGTTGAATTGAGAGTTCAATATTATAGATTCTCTATATCATGGTCTAGAATACTTCCAACCGG
TCATGCAAATGAAGTTAATGAGGCGGGAGTGCAATATTATAGTGACTTAATCGATAAACTTCTTGAAAATGGGATAACCC
CTTTGGTAACGATGTATCATTGGGATTTACCTCAACCGTTACAAGAAATTGGAGGCTGGACCAGCCCAGTACTAGTTGAC
CTCTTCGTAGATTATGCTAATGTTCTCTTTACAAGTTTTGGCGACAGAGTGACGAATTGGATTACTTTCAATGAACCCTA
TCAGATTTGTCAACAAGGATATTCAGAAGGCACTAAGGCACCGGCTTACACACAAGATGGTGTCGGTGGATATCTTTGTG
GTCATACTTTGCTGCTGGCCCATGCTTACACTTATCACCTTTATCACGATGACTATGCAGGAACGGATGGTAGAATTGGT

ATAACAGTTAATGGAGTGTGGGCTGAAGCAGCATCTGAAGATTCTGTAGATCAAACCGCTGCTGACGATTATATAGAATTTCATTTTGGTTGGTTCTTTAATCCAATTTATATCGGAGACTATCCTCAAATCATGAAGGATAATATACAGGCCAGAAGTGAGCTCGAAGGTTTTCCAGATACGAGATTACCTGAGTTCACCGAAACAGAAATTGCACTCTTACGTAGCTCTTCAGACTTTCTCGGTCTAAACCATTACACAACCTATCAATGCACACCATTAGAAGATGTAGATTCGCTTGAACAACCTTCATTTAAAGTAGACAGCGGCGTTAATTGCTGGTCACCTACTGATTGGGAAGGTGGTGCTTCATCTTGGTTGAAGGTCCATCCTTCCGGTCTTACTAGCATATTGAATTGGATCAGGGAAAAATACGATAATCCCGAAGTTATAATAACTGAAAATGGTTTTTCTGAGGCTGGTGAGGTGAATGACCTAAACGACTGTGATAGAGTTAGTTACTATAATGGATACTTATATGCCGTTTTGGATGCCTTGGAAGATGGATGCCAAGTATCCGGGTATATGGCTTGGAGCCTCATGGATAACTTTGAATGGATGAGAGGCTACAGCGAACGATTTGGATTATACTATGTCGATTTCGAAGATGAAGACAGGCCCAGAACTGCAAAGAAATCTGCCCTCGTTTATAAAAATATAATTGAAACTAGAGCAATAGATTTGGATTATGATCCATCAGAATTGGGAACTTGTACTACTTCTGACGAGGACGAAGTGCCAGAG

> *Mm_bGlc15*

ATGAAGCTCGTAATTTTCGCTCTGATTTTAACATACGTTAGCGCCCAAGATACTCGATTTCCCGATGGATTTAAATTTGGGGTTGCTACGGCATCGTATCAGATAGAAGGCGCTTGGGATGCTAACGGTAAAGGAGAAAATATATGGGATAGGTTGACGCATACAAGACCTGAACTAATCGCAGACGGCAGCAATGGTGATATCGCATGCGATGCTTACCACAAAACTGAAGAAGACGTTCAGTTGCTGAAAGGTCTGGGAGTTGATTTCTATCGTTTCTCCATCTCATGGTCGAGAATTCTTCCCACTGGTTATGCCGACGAAATAAATGAAGATGGTATTCGCTATTACAACGAATTAATTGATGAACTTCTGGCAAATGATATCACGCCATATGTGACAATGTATCATTGGGATTTGCCGCAGCCGTTGCAGGAGATAGGTGGCTGGCTAAACTCTTCTTTAGCTGATATTTTTGTTGATTACGCAGATGTGCTTTATGAGCAATTCGGTGACAGAGTGAAAGATTGGATAACGTTTAATGAACCCATCCAGGTTTGCGAAGCAGGTTATTCGAATGCTGGAAAAGCTCCAGCTTATACAATGGCAGGAGTTGGAGGTTATTTGTGTGGTCATACTTTGCTTATTGCACATGGAAAGACTTATCGACTTTACAATGAAAAATATAGGGAAAGTCAACAAGGTAGGGTTGGTATAACTATCGATGCTGGTTGGTATGAACCAGTTTCGGACTCAGATACAGATATAGAAGCTGCCGAACGTAGCATGCAAATGAACTATGGTTGGTATGCACATCCAATCTTTTCTGAAACGGGGGATTACCCTGAAATTATGAAGGAAAGGGTGGCAGAATTAAGTGAATTAGAAGGCTTCGAAACTTCTAGATTACCTTCCTTTACTCAAGAAGAAATCGAATTCATACGAGGCTCGTCCGACTTCCTAGGTCTTAATCATTACAGTAGTTCCTTGTGTACATCGATACCAGAAGAATGGGCTTTAACTGGACCTAGTCAATGGATAGACGTTGGTAGTTTGTGCTTACCCAGTCCCGAGTGGGAAGTGGCTGCTTCTAGTTGGTTGTATGTTGTTCCTTGGGGATTAAGAAGACTGTTAAATTGGATTAAGAACGAATATGGCAATCCCGAGGTTATAATTACTGAGAATGGATTTTCTGATACCGGTGAATTAAATGATTGTCGAAGAGTGAATTATTATAATTCATATTTAACGGCAGTTCTAGAGGCAATCTTAGAAGATGGCTGCAACGTTTCAGGATATACGGCTTGGAGTTTTATGGATAATTTTGAATGGTTGATGGGTTACACGGAACGATTTGGAA

TTCACTATGTCGATTTCGAGGATCCCGACAGACCAAGAACTGCTAAAATGTCCGCTCATGTCTTTAAAAATATTATAGAA
ACAAGGGAAATCGATTGGAATTACACTCCAGATGGTTTCGAAGAGTGCGATTGGTCATAA

> *Mm_bGlc16*

ATGAAGGTTCTAGTTATACTTTGTACAGTAATATATTTATGTAAAGCCCAAGCTTACGAATTTCCAGACGATTTTCTATT
TGGCGTTGCTACGGCAGCTTATCAGATAGAAGGTGCTTGGGATACTAATGGAAAAGGAGAAAATATATGGGATAATTTGA
CGCATACCTATCCACACTTGGTTGTAGATGGTTCTAATGGAGATGTTGCATGCGATGCCTACAGTCATACTGCAGAAGAC
GTCCAACTGCTCAAAAATTTAGGCGTCGACTTTTATAGGTTTTCCTTCTCTTGGTCGAGAGTCTTACCCACTGGAAAGAC
TGATTACATAAACCCAGATGGAATACGCTATTATAATGAACTAATTGACACTTTATTAGAGAATAATATAGAACCTATGG
CCACGATGTATCACTTTGATTTACCACAACCTCTTGAAGACGAAGGAGGTTTCCTGAACATAGTCATCGCTGATTATTTC
GAAGATTACGCGGAAGTTCTGTACGAGAATTTTGGTGATCGCATCAAACAATGGATAACGTTTAATGAACCGTCTCCTAT
ATGTGAAAATGGATACGGCGCAGACAGTATGGCACCTTTGGCAAATCAACCGGGAGTTGGTGGTTATATCTGTAGTCGAA
CTTTATTAGTGGCCCATGCTAAAGCTTATCATCTCTACAATGATCGTTACAAGGGAAGTCAAGGAGGTCGAGTGGGTATT
ACTATAAACAACTTTTGGTATGAACCTTATGCTGATACTACAGACTCAGCAGTAGAGTTAGCATTACAGGTTGCGTTTGG
TTGGTTTACTCATCCCATATATTCTAAAGAAGGCGATTTCCCACCAGCCATGAAAGAAAGAATAGCTAGATTAAGTGAAG
AGGAAGGCTTTTCTGCATCCAGACTACCAGAACTTACAACGGAAGAAATAGAATTAATAAAAGGATCATCTGACTTTTTC
GGTTTGAATCACTACACCACTCACTTCTGTTCCGAGTCTGGTATCGACTCGATATCACGACCATCCCATAATTACGACAT
GGGTGTAACATGCATGCCAGATTATAGTTACGAAGTCGCCGGATCGTTTTGGTTGAGCGTTATTCCTTGGGGGTTGAGGA
AGATGCTTAATTGGATCAAAGAAGAATACAACAATCCGGAAATTATAATCACTGAAAATGGATACTCCGATAATGAAGTT
ATTTTGAACGATTGCCGCAGAATTAATTACTATAATTCTTATTTAACAGAATTGTTAAATGCTATTTACGAAGATAACTG
TAACGTCACTGGTTACACAGCGTGGAGTTTTATGGATAATTTTGAATGGAGAATGGGTTATAGCGAAAAATTTGGTTTAT
ACTCAGTAGATTTTAATGATCCCGATAGGACAAGGACAGCTAAAATGTCAGCATACATATACAAAAATATTATTGAAACA
AGGCAAATTGACTGGAGTTATGTGCCCGAAAATTTTAGCGACTGTGAATGG

> *Mm_bGlc17*

ATGAAGAGACTAATTCTTATTTTCGTATTGGCTTGTTTAGCCAAAGCACAAGATTATCAATTTCCTGAAGGGTTTAAATT
TGGTGTCGCTACAGCCTCGTATCAAGTGGAGGGCGCTTGGAACGAAAATGGCAAAGGCGAAAACATATGGGACAGATTAA
CACACACCCAACCTGATTTAATCGCTGATAATAGCAATGGTGACATCGCTTGTGATGCCTACCACAAAACCGAAGAAGAC
GTTCAATTATTAAAGAACCTTGGAGTCAATTTCTACCGTTTCTCGATATCGTGGTCGAGAATTCTTCCAACAGGTTACAC
GAATGTCGTGAACGAAGATGGTATTCGATACTATAACGCATTAATCGACGCTCTTTTGGAAAACGGTATTACTCCGCTTG
TAACGATGTTCCATTGGGATTTACCACAGCCGTTGCAAGAAATCGGTGGCTGGCCCAATCCGCTTCTGGCTGATATATTT

GCAGATTATGCTGACATACTTTATCGAGAATTCGGAGATAGGGTGAAGGATTGGCTAACTTTCAACGAACCCACTCCCAT
TTGTATTGGAGGTTATTCGGAGGGTTGGATGGCGCCGGCTTATGAATTACAAGGTGTTGGAGGTTACCTGTGTGGTCATA
CATTACTTATTGCTCATGGGAAAGCTTATCGGCTTTACAACGAAAAGTATAGAGATACTCAGGAAGGTAGAGTTGGTATA
ACAATTGACAGCGGTTGGTATGAACCTGCTTCAGAATCAGAAGAAGACATTCAAGCAGCTGAAGACAGTGTACATATAAA
GTACGGTTGGATGGTCCATCCGATCTATTCAGAGACCGGTGATTATCCCCCTGTATTGAGAGAAAGGGTAGACGCATTGA
GTGCTGAAGAAGGCTTTGCGAGGTCCAGATTGCCTATCTTTACGGAGGAGGAAATCGAACTTATTAAAGGCTCTAGTGAT
TTTCTGGGTTTGAATCACTATACTACCAATCTCTGCACCCCCATTCCGGAAGAATGGGGTGTGGTGGGTCCTTCACATTA
CGTAGATAGTGGTGCTAATTGTTACCAAGATCCCTCGTGGGAAGGTTCGGGCTCGTCTTGGTTAAAGGTTGTACCTTGGG
GTTTGAGGAGATTGTTGAATTGGATTAAGGAAAATTACGACAATCCCGAAGTATTAATAACGGAGAATGGAGTTTCTGAT
AATACTGGTATTTTGAATGATTGCAGGAGGATCAATTTCTATAACACATATTTAACGGCAGTTCTTGAAGCTATTCACGT
GGACGGTTGCAATGTGGTTGGATATACAGCTTGGAGTTTCATGGACAACTTCGAATGGATGCAGGGATATACTGAACGAT
TTGGACTTTACCACGTCGATTTTAACGACACCGACAGGACGAGAACTCCAAAAATGTCATCCCATGTTTACAGACACATT
ATTGAGACTAAGCAAATAGACTGGGATTATACGCCTCATGGATTTGATGCCTGCGAGTGG

> *Mm_bGlc18*

ATGGGATACTTTGAACCATTAATAGTTTTCTTGATATCTTTATGGCGAGCCCAGAGTTATGTTTTTCCGGAAGAATTTTT
GTTTGGTGTTGCTACGTCCACGTATCAAGTCGAAGGTGCCTGGAATCTAAGTGGTAAAGGTGTAAATATATGGGACCACT
TGACCCATACAAATCCTGAGTTCACGGCGGACGGCAGTAATGGTGATGTAGCTTGCGATGCCTATCACAAAACTAAAGAA
GATGTCCAACTTATGGTGGATATGGGTCTTGACGTTTATCGTTTTTCCCTATCATGGACGAGAATTCTTCCAAACGGTTA
CAATAATTACACGAATCCCGATGGTGTTCGCTACTATAACGAATTGATCGATACTTTATTGGCGAATGGCATAAAGCCAT
TAGTCACGATGTTTCATTGGGACATACCGCAAGTATTTCAAGACGATTACGGTGGATGGTTAGGCTCGGATATGGTAGAT
ATTTTCGTCGATTATGCTGATGTCGCTTTTTCGTTGTTTGGCGATAGAGTGAAGGATTGGATTACTTTTAATGAACTTCA
TATATTTTGTGAACTTGGACATTCCATGGATATTATGGCACCGGCTCTTGGATTGAGCGGAGTTGGTGGTTATTTATGCG
CCCATAATATATTAATAGCTCATGGAAAGACTTATAAATTGTACAATGAGAAGTACAGGGATATTCAGAAAGGTAGAATT
GGTATCACAATTGATGGTGAATGGAAAGAACCGGCATCCACTTGTCCTGAAGATATAGAAACGGCAGAACGAGCTCTACA
AATGGAGTTTGGTTGGATAGCGCACCCAATCTTCTCAGAATCCGGCGATTATCCACCCGTTATGAGGAGCAGAATTGACG
TGATGAGTGCCGAAGAAGGTCTGAAGACATCAAGATTGCCCTATTTTTCCAAAGAGGAGATCGAATTAATAAGAGGATCG
GCTGATTTCCTAGGTTTAAATCACTACACGGCTTCGTTGTGTTCGTCAGAATTCAAAGAGTTACCAAGCAGGCCTTCGTA
TACTAGTGACACTGGAGCTAATTGTTATCAACCCGATTATTGGGAACCTACTGGCGTGTCGCAATTTAAAGTTACGCCAT
GGGCTTTTGGAAAGTTGTTGGTTTGGATCAAAAACGAATATAACAATCCGGAAGTTATCGTTACAGAGAATGGATATTCT

GATAATACGGGTGATTTATATGATTGTAGAAGAGTATATTATTACAATTCTTACCTTACTGAACTATTACATGTTGTTAACGATGAAGGATGTAGGATTACTGGTTACATGGCTTGGAGTTTCATGGATAATTTCGAATGGGGTAATGGTTACACAGCTAAATTTGGGATTGTTAACGTCGACTTTAATGATGCTGACAGACCGAGAACTCACAAGATGTCCTCGTTCGTTTACAGAAATATAATCCAGACGAGAGCGATCGATTGGAACTTTACTCCAGAGGGATTTGAAGCATGTTCGTGGTGG

> *Mm_bGlc19v*

ATGTCGTCATATGTATACAAAAACATTATAGAAACTAAAACAATCGACGAAGATTTTGTACCTGAAGGATTTGCTGCTTGCAGTTCTAGTGAATACACCTTCCCTGACGGTTTCATATTTGGTGTCGCTACTTCAGCTTATCAAGTGGAAGGTGCTTGGGAAGATGATGGAAAGGGAGAGAATATCTGGGACCATTTAACACACACAAGACCGACCGCCATAACTGATGAGAGTAACGGTGACATCGCTTGCGACACGTATCACAAGACGGTGGAAGATGTTCAGTTACTAAACACATTGGGCGTAGATTTTTACCGTTTTTCACTATCTTGGTCAAGACTTCTCCCTACCGGTCACGCCAACACTGTAAATCCTTTAGGCGTTGCTTACTATAACGAATTAATAGACGAGTTGATAGCAAATGATATAACACCATTGGTTACACTATTTCATTGGGATCTGCCACAACCACTGCAAGAAATTGGCGGTTGGCCTAATCCACTTCTTGTGGATTTATATGCCGATTATGCTAACATTGTTTTCAGTCTGTTTGGTGATCGCGTTAAAGATTGGATTACGTTCAATGAGCCTTATCAGATATGTCAAGAGGCTTATTCGGAGGCAAATAAAGCTCCTGCATACAATCAAGATGGCACTGGTGGCTATTTGTGCGCTTACACAGTATTACTTGCACATGCAAGAGCGTATCGTTTGTACGAGACAACATATAAGGAAGCACAACAAGGTAGAGTTGGAATTACCGTGCACGGAATATGGGCAGAACCTGAAACCGCAACTGATGCTGATATAGCTGCCGCTGAAAGTTATCAACAATTCCACTTTGGTATATATCTTCATCCAATATTCTCAGAAGAGGGCGATTTTCCGGAAATTGTAAAAACTAGAGTGAAGCAGATAAGTGACGTCATAGGCTACTCCAACAATCGCTTGCCAGTATTCACGGCTGAAGAAATCGAGTATATCCATGGTACCTCCGATTTCCTGGGCTTCAATCATTATTCGACAGACCTATGTAAAGCAGCTGACGACGCTTCGTTAGTGCATCCATCCAATAAGGGTGATACTGGGGCTGATTGCCAAAAGTCCGATGACTGGGAAAGCGCCGCTTCTTCGTGGGTGAAAGTTGTTCCGTGGGGTTTCAGAAAATTGTTAAACTGGATTAAAGACGAATATAATAATCCAGAAGTACTGATTACTGAAAATGGTTTCTCCACATTTGGTGATGATTTGAACGACTGCAGAAGAATTAACTACTTCAACGAATATCTAACCGCCCTTTTGGAAGCAATACACGAGGACGGATGCAACGTTATAGGATACACAGCTTGGAGCTTCCTTGATAATTTCGAATGGATGGATGGTTATTTAGAGAAGTTCGGTTTATACATGGTCGACTTTGACGATCCAGATAGACCGAGACAACCAAAAAATGTCTTCCTATGTTTACACAAATATCATCCAGACACGAAGAGTCGACACGAGCTTCACCCCTGAAGGTTTTGAAGCTTGCATATTTGATGAAGATACCGAGTCAGATGAAGATGTTTAA

**Text S3.** Fragment nucleotide sequence of *tubulin*, *GFP*, *Mm_bGlc16*, *Mm_bGlc17*, *Mm_bGlc18* in *pCR®2.1-TOPO® and pIB/V5-His-TOPO®* vectors. The T7-Promotor sequence for *in vitro* RNA synthesis are highlighted in yellow.

*Tubulin forward in pCR®2.1-TOPO®*

GTT CAT CCA TGC CAT GTG TAA TCC CAG CAG CTG TTA CAA ACT CAA GAA GGA CCA TGT GGT CTC TCT TTT CGT TGG GAT CTT TCG AAA GGG CAG ATT GTG TGG ACA GGT AAT GGT TGT CTG GTA AAA GGA CAG GGC CAT CGC CAA TTG GAG TAT TTT GTT GAT AAT GGT CTG CTA GTT GAA CGC TTC CAT CTT CAA TGT TGT GTC TGG TTT TGA AGT TAA CTT TGA TTC CAT TCT TTT GTT TGT CTG CAG TGA TGT AGA CCT TGT GGC TGT TGT AGT TGT ATT CCA ACT TGT GGC CGA GGA TGT TTC CGT CCT CCT TGA AAT CGA TTC CCT TAA GCT CGA TCC TGT TGA CGA GGG TGT CTC CCT CAA ACT TGA CTT CAG CAC GTG TCT TGT AGT TCC CGT CGT CCT TGA AGA AGA TGG TCC TCT CCT GTA CGT ATC CCT CAG GCA TGG CGC TCT TGA AGA AGT CGT GCT GCT TCA TAT GAT CTG GGT ATC TTG CAA AGC

*Tubulin rev in pCR®2.1-TOPO®*

GCT TTG CAA GAT ACC CAG ATC ATA TGA AGC AGC ACG ACT TCT TCA AGA GCG CCA TGC CTG AGG GAT ACG TAC AGG AGA GGA CCA TCT TCT TCA AGG ACG ACG GGA ACT ACA AGA CAC GTG CTG AAG TCA AGT TTG AGG GAG ACA CCC TCG TCA ACA GGA TCG AGC TTA AGG GAA TCG ATT TCA AGG AGG ACG GAA ACA TCC TCG GCC ACA AGT TGG AAT ACA ACT ACA ACA GCC ACA AGG TCT ACA TCA CTG CAG ACA AAC AAA AGA ATG GAA TCA AAG TTA ACT TCA AAA CCA GAC ACA ACA TTG AAG ATG GAA GCG TTC AAC TAG CAG ACC ATT ATC AAC AAA ATA CTC CAA TTG GCG ATG GCC CTG TCC TTT TAC CAG ACA ACC ATT ACC TGT CCA CAC AAT CTG CCC TTT CGA AAG ATC CCA ACG AAA AGA GAG ACC ACA TGG TCC TTC TTG AGT TTG TAA CAG CTG CTG GGA TTA CAC ATG GCA TGG ATG AAC

*GFP forward in pCR®2.1-TOPO®*

GTT CAT CCA TGC CAT GTG TAA TCC CAG CAG CTG TTA CAA ACT CAA GAA GGA CCA TGT GGT CTC TCT TTT CGT TGG GAT CTT TCG AAA GGG CAG ATT GTG TGG ACA GGT AAT GGT TGT CTG GTA AAA GGA CAG GGC CAT CGC CAA TTG GAG TAT TTT GTT GAT AAT GGT CTG CTA GTT GAA CGC TTC CAT CTT CAA TGT TGT GTC TGG TTT TGA AGT TAA CTT TGA TTC CAT TCT TTT GTT TGT CTG CAG TGA TGT AGA CCT TGT GGC TGT TGT AGT TGT ATT CCA ACT TGT GGC CGA GGA TGT TTC CGT CCT CCT TGA AAT CGA TTC CCT TAA GCT CGA TCC TGT TGA CGA GGG TGT CTC CCT CAA ACT TGA CTT CAG CAC GTG TCT TGT AGT TCC CGT CGT CCT TGA AGA AGA TGG TCC TCT CCT GTA CGT ATC CCT CAG GCA TGG CGC TCT TGA AGA AGT CGT GCT GCT TCA TAT GAT CTG GGT ATC TTG CAA AGC

*GFP rev in pCR®2.1-TOPO®*

GCT TTG CAA GAT ACC CAG ATC ATA TGA AGC AGC ACG ACT TCT TCA AGA GCG CCA TGC CTG AGG GAT ACG TAC AGG AGA GGA CCA TCT TCT TCA AGG ACG ACG GGA ACT ACA AGA CAC GTG CTG AAG TCA AGT TTG AGG GAG ACA CCC TCG TCA ACA GGA TCG AGC TTA AGG GAA TCG ATT TCA AGG AGG ACG GAA ACA TCC TCG GCC ACA AGT TGG AAT ACA ACT ACA ACA GCC ACA AGG TCT ACA TCA CTG CAG ACA AAC AAA AGA ATG GAA TCA AAG TTA ACT TCA AAA CCA GAC ACA ACA TTG AAG ATG GAA GCG TTC AAC TAG CAG ACC ATT ATC AAC AAA ATA CTC CAA TTG GCG ATG GCC CTG TCC TTT TAC CAG ACA ACC ATT ACC TGT CCA CAC AAT CTG CCC TTT CGA AAG ATC CCA ACG AAA AGA GAG ACC ACA TGG TCC TTC TTG AGT TTG TAA CAG CTG CTG GGA TTA CAC ATG GCA TGG ATG AAC

*Mm_bGluc 16 forward in pIB/V5-His-TOPO®*

TTT AAT ACG ACT CAC TAT AGG GAG AAC AGT ATG GCA CCT TTG GCA AAT CAA CCG GGA GTT GGT GGT TAT ATC TGT AGT CGA ACT TTA TTA GTG GCC CAT GCT AAA GCT TAT CAT CTC TAC AAT GAT CGT TAC AAG GGA AGT CAA GGA GGT CGA GTG GGT ATT ACT ATA AAC AAC TTT TGG TAT GAA CCT TAT GCT GAT ACT ACA GAC TCA GCA GTA GAG TTA GCA TTA CAG GTT GCG TTT GGT TGG TTT ACT CAT CCC ATA TAT TCT AAA GAA GGC GAT TTC CCA CCA GCC ATG AAA GAA AGA ATA GCT AGA A

*Mm_bGluc 17 reverse in pIB/V5-His-TOPO®*

TTT AAT ACG ACT CAC TAT AGG GAG ATC TAG CTA TTC TTT CTT TCA TGG CTG GTG GGA AAT CGC CTT CTT TAG AAT ATA TGG GAT GAG TAA ACC AAC CAA ACG CAA CCT GTA ATG CTA ACT CTA CTG CTG AGT CTG TAG TAT CAG CAT AAG GTT CAT ACC AAA AGT TGT TTA TAG TAA TAC CCA CTC GAC CTC CTT GAC TTC CCT TGT AAC GAT CAT TGT AGA GAT GAT AAG CTT TAG CAT GGG CCA CTA ATA AAG TTC GAC TAC AGA TAT AAC CAC CAA CTC CCG GTT GAT TTG CCA AAG GTG CCA TAC TGT A

*Mm_bGluc 17 forward in pIB/V5-His-TOPO®*

TTT AAT ACG ACT CAC TAT AGG GAG AAC CCA CTC CCA TTT GTA TTG GAG GTT ATT CGG AGG GTT GGA TGG CGC CGG CTT ATG AAT TAC AAG GTG TTG GAG GTT ACC TGT GTG GTC ATA CAT TAC TTA TTG CTC ATG GGA AAG CTT ATC GGC TTT ACA ACG AAA AGT ATA GAG ATA CTC AGG AAG GTA GAG TTG GTA TAA CAA TTG ACA GCG GTT GGT ATG AAC CTG CTT CAG AAT CAG AAG AAG ACA TTC AAG CAG CTG AAG ACA GTG TAC ATA TAA AGT ACG GTT GGA TGG TCC ATC CGA TCT ATT CAG AGA CCG GTG ATT ATC CCC CTG TAT TGA GAG AAA GGG TAG ACG CAT TGA GTG CTG AAG AAG GCT TTG CGA GGT CCA GAT TGC CTA TCA

*Mm_bGluc 17 reverse in pIB/V5-His-TOPO®*

TTT AAT ACG ACT CAC TAT AGG GAG AGA TAG GCA ATC TGG ACC TCG CAA AGC CTT CTT CAG CAC TCA ATG CGT CTA CCC TTT CTC TCA ATA CAG GGG GAT AAT CAC CGG TCT CTG AAT AGA TCG GAT GGA CCA TCC AAC CGT ACT TTA TAT GTA CAC TGT CTT CAG CTG CTT GAA TGT CTT CTT CTG ATT CTG AAG CAG GTT CAT ACC AAC CGC TGT CAA TTG TTA TAC CAA CTC TAC CTT CCT GAG TAT CTC TAT ACT TTT CGT TGT AAA GCC GAT AAG CTT TCC CAT GAG CAA TAA GTA ATG TAT GAC CAC ACA GGT AAC CTC CAA CAC CTT GTA ATT CAT AAG CCG GCG CCA TCC AAC CCT CCG AAT AAC CTC CAA TAC AAA TGG GAG TGG GTA

*Mm_bGluc 18 forward in pIB/V5-His-TOPO®*

TTT AAT ACG ACT CAC TAT AGG GAG ATT CAT ATA TTT TGT GAA CTT GGA CAT TCC ATG GAT ATT ATG GCA CCG GCT CTT GGA TTG AGC GGA GTT GGT GGT TAT TTA TGC GCC CAT AAT ATA TTA ATA GCT CAT GGA AAG ACT TAT AAA TTG TAC AAT GAG AAG TAC AGG GAT ATT CAG AAA GGT AGA ATT GGT ATC ACA ATT GAT GGT GAA TGG AAA GAA CCG GCA TCC ACT TGT CCT GAA GAT ATA GAA ACG GCA GAA CGA GCT CTA CAA ATG GAG TTT GGT TGG ATA GCG CAC CCA ATC TTC TCA GAA TCC GGC GAT TAT CCA CCC GTT ATG AGG AGC AGA ATT GAC GTG ATG AGT GCC GAA GAA GGT CTG AAG ACA

*Mm_bGluc 18 reverse in pIB/V5-His-TOPO®*

TTT AAT ACG ACT CAC TAT AGG GAG AGT CTT CAG ACC TTC TTC GGC ACT CAT CAC GTC AAT TCT GCT CCT CAT AAC GGG TGG ATA ATC GCC GGA TTC TGA GAA GAT TGG GTG CGC TAT CCA ACC AAA CTC CAT TTG TAG AGC TCG TTC TGC CGT TTC TAT ATC TTC AGG ACA AGT GGA TGC CGG TTC TTT CCA TTC ACC ATC AAT TGT GAT ACC AAT TCT ACC TTT CTG AAT ATC CCT GTA CTT CTC ATT GTA CAA TTT ATA AGT CTT TCC ATG AGC TAT TAA TAT ATT ATG GGC GCA TAA ATA ACC ACC AAC TCC GCT CAA TCC AAG AGC CGG TGC CAT AAT ATC CAT GGA ATG TCC AAG TTC ACA AAA TAT ATG AAA

**SI Figures**

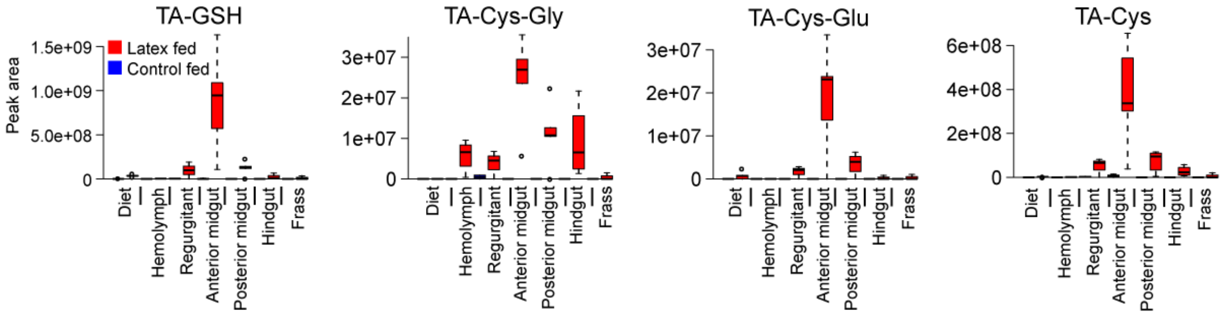


**Fig S1.** Relative quantification of TA-glutathione conjugates in *Melolontha melolontha* larvae feeding on diet with and without *Taraxacum officinale* latex. N = 5. TA = taraxinic acid ; gsh = glutathione; cys = cysteine; gly = glycine; glu = glutamate.

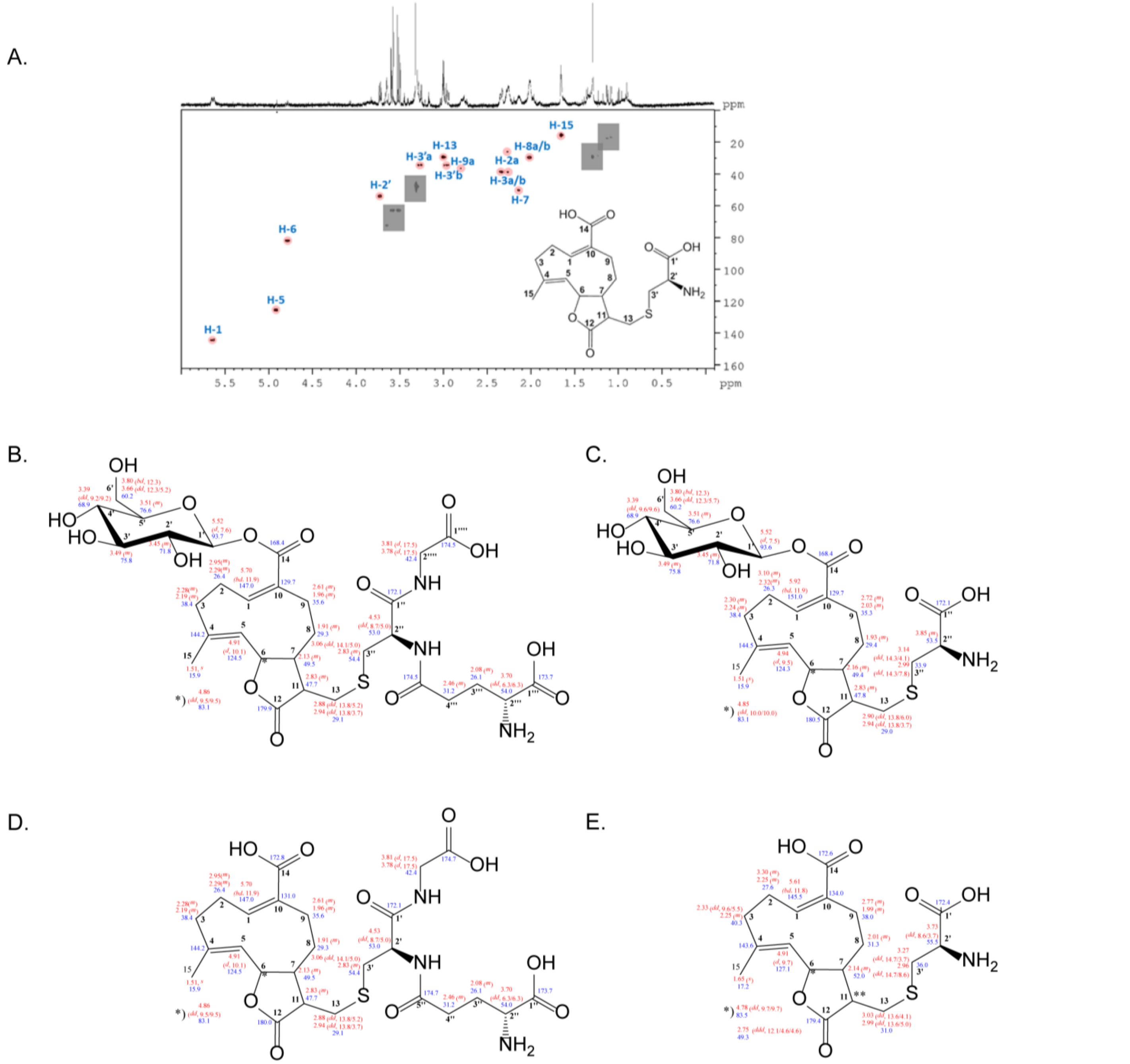


**Fig S2.** A. 500 MHz $^{1}$H-$^{13}$C HSQC NMR spectrum of the partially purified *M. melolontha* midgut extract. Intensity level was adjusted to suppress noise. Grey rectangles mask impurities for clarity. Resonances above noise level are highlighted by pink circles. Missing resonances in the noise level are not considered in the picture. Sample was measured in MeOH-*d*$_4$. B. Structure of synthesized TAG-GSH with chemical shifts (500 MHz, in MeOD-*d*$_4$). C. Structure of synthesized TAG-Cys with chemical shifts (500 MHz, in MeOD-*d*$_4$). D. Structure of synthesized TA-GSH with chemical shifts (500 MHz, in MeOD-*d*$_4$). E. Structure of synthesized TA-Cys with chemical shifts (500 MHz, in MeOD-*d*$_4$).

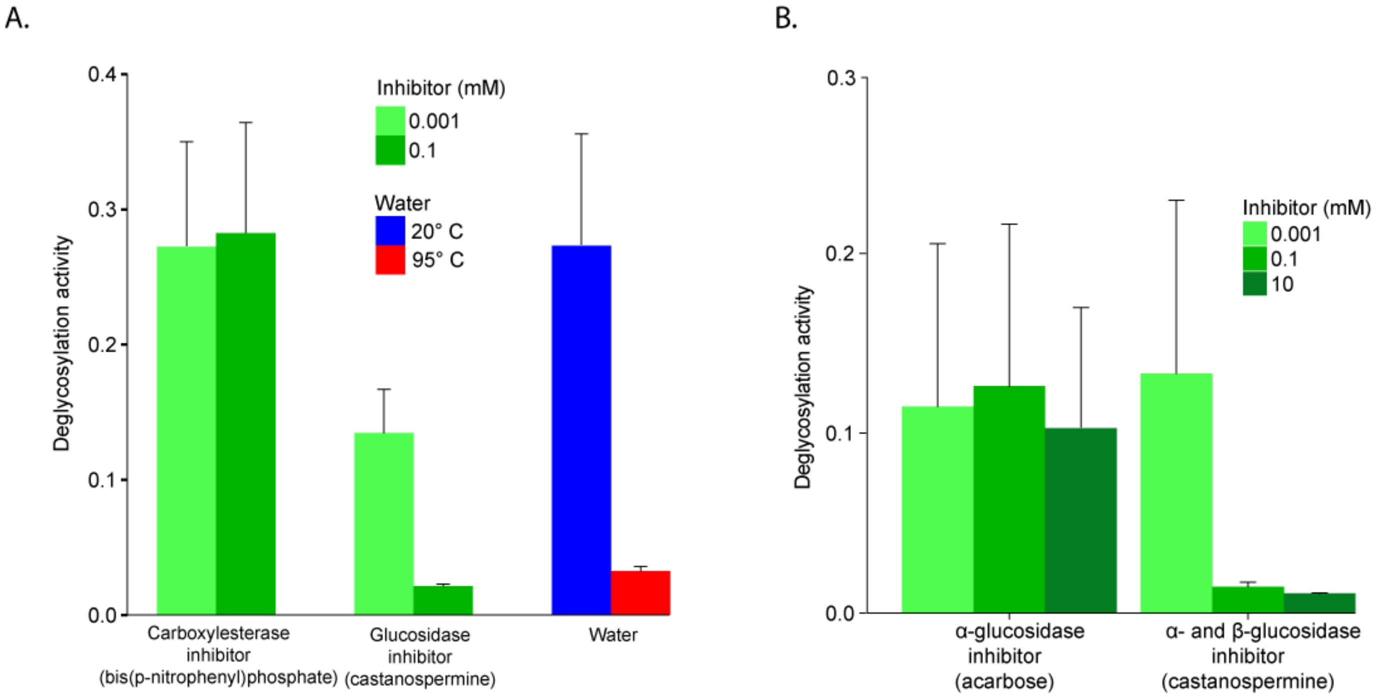


**Fig S3.** A. TA-G deglucosylation activity (TA/(TA+TA-G)) of *Melolontha melolontha* anterior midgut samples in the presence of either the carboxylesterase inhibitor bis(p-nitrophenyl)phosphate or the α- and β- glucosidase inhibitor castanospermin. Only the glucosidase inhibitor reduced deglucosylation of TA-G. N = 6. Error bars denote SEM. B. TA-G deglucosylation activity (TA/(TA+TA-G) of *M. melolontha* anterior midgut samples in the presence of acarbose, an α-glucosidase specific inhibitor, or castanospermine, an α- and β-glucosidase inhibitor. Only castanospermine reduced deglucosylation of TA-G. TA-G = taraxinic acid β-D-glucopyranosyl ester. TA = taraxinic acid. N = 3. Standard errors denote SEM.

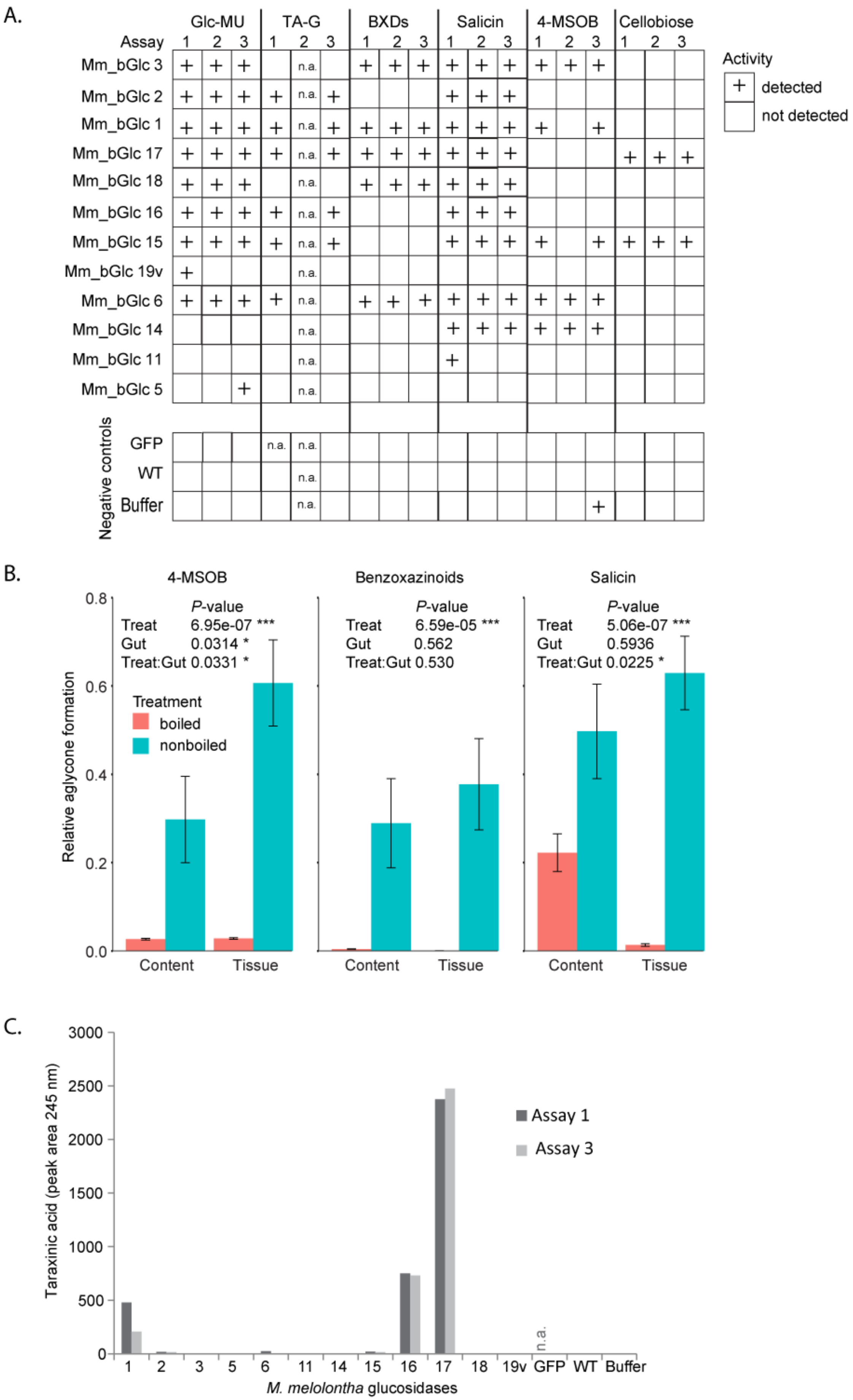
A.
Glc-MU
TA-G
BXDs
Salicin
4-MSOB
Cellobiose
Assay 1 2 3
Mm_bGlc 3
Mm_bGlc 2
Mm_bGlc 1
Mm_bGlc 17
Mm_bGlc 18
Mm_bGlc 16
Mm_bGlc 15
Mm_bGlc 19v
Mm_bGlc 6
Mm_bGlc 14
Mm_bGlc 11
Mm_bGlc 5
n.a.
Negative controls
GFP
WT
Buffer
Activity
+ detected
not detected
B.
4-MSOB
Benzoxazinoids
Salicin
P-value
Treat 6.95e-07 ***
Gut 0.0314 *
Treat:Gut 0.0331 *
Treat 6.59e-05 ***
Gut 0.562
Treat:Gut 0.530
Treat 5.06e-07 ***
Gut 0.5936
Treat:Gut 0.0225 *
Treatment
boiled
nonboiled
Relative aglycone formation
Content
Tissue
C.
Taraxinic acid (peak area 245 nm)
Assay 1
Assay 3
n.a.
1 2 3 5 6 11 14 15 16 17 18 19v GFP WT Buffer
M. melolontha glucosidases

**Fig S4.** A. Activity of heterologously expressed *Melolontha melolontha* β-glucosidases and negative controls (GFP = green fluorescent protein; WT = non-transformed wild type; buffer) towards plant defensive glycosides, cellobiose and the standard substrate 4-methylumbelliferyl-β-D-glucopyranoside (Glc-MU), which fluoresces upon deglucosylation. The glucosidase activity of three deglucosylation assays was categorized according to relative activity. Both total protein levels and catalytic activity may contribute to the overall activity. N.a. = not available. B. Deglucosylation of defensive glycosides by boiled and non-boiled *M. melolontha* anterior midgut extracts *in vitro*. Aglycone formation was normalized to the maximal peak area of all samples. *P*-values of two-way ANOVAs are shown for each metabolite separately. N = 10. Error bars denote SEM. TA-G = taraxinic acid β-D-glucopyranosyl ester; BXDs = benzoxazinoids; 4-MSOB = 4-methylsulfinylbutyl glucosinolate. C. Taraxinic acid aglycone formation of the heterologously expressed *M. melolontha* β-glucosidases and negative controls (GFP = green fluorescent protein; WT = non-transformed wild type; buffer) of two deglucosylation assays. Both total protein levels and catalytic activity may contribute to the aglycone formation. N.a. = not available

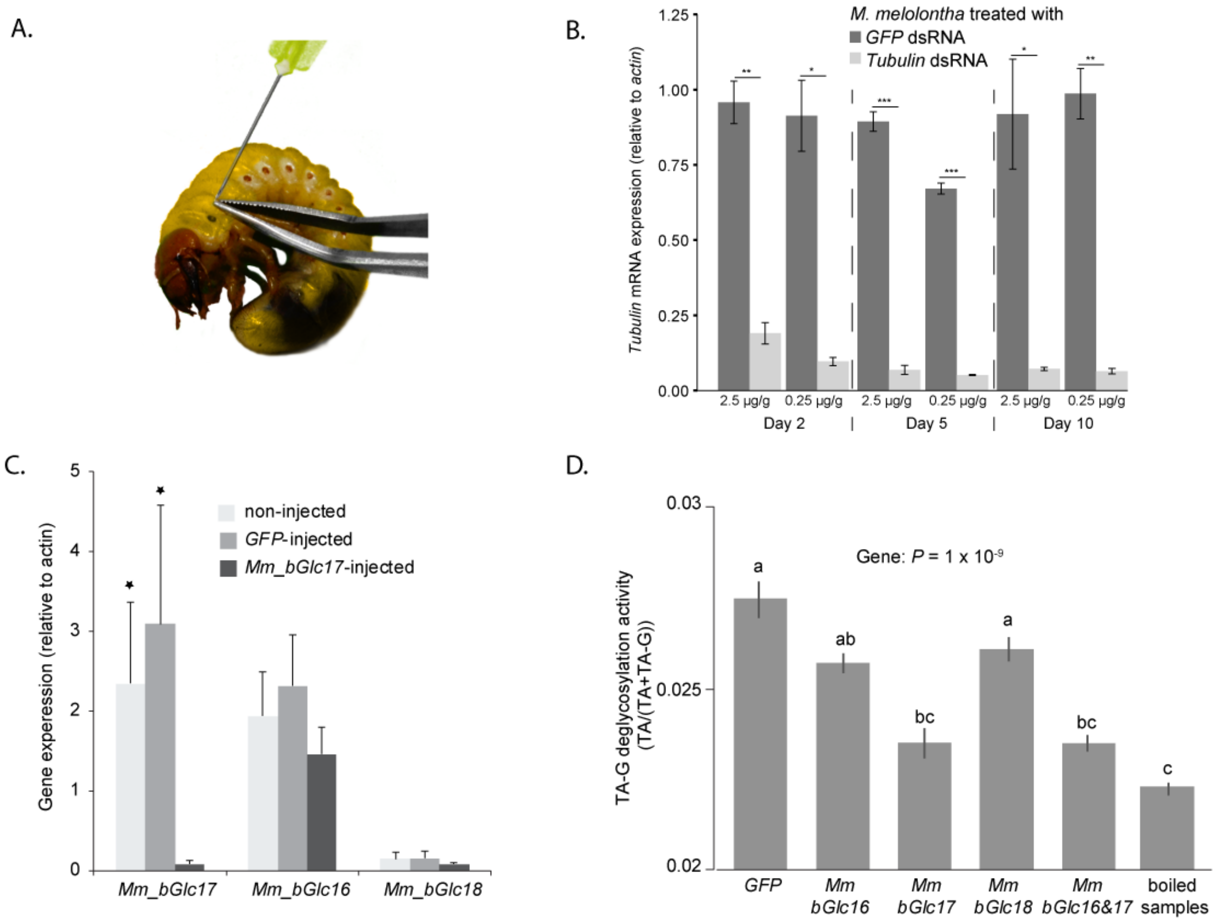


**Fig S5**. A. Injection of dsRNA by a sterile syringe between the second and third segment of *Melolontha melolontha*. B. *Tubulin* mRNA expression in *M. melolontha* larvae treated with 2.5 µg or 0.25 µg *GFP* or *tubulin* dsRNA per g larval mass 2, 5 and 10 days after injection. N=3. Error bar denote SEM. Asterisks indicate significant differences in relative *tubulin* expression between *GFP* and *tubulin* dsRNA treated larvae according to a two-tailed Student's *t*-test (* $P < 0.5$; ** < 0.01; *** < 0.001). C. Gene expression (relative of actin) of *Mm_bGlc16*, *Mm_bGlc17* and *Mm_bGlc18* in *Mm_bGlc17* dsRNA injected larvae 2 days after dsRNA application. Asterisks indicate significant differences in the relative gene expression between non-injected and *Mm_bGlc17* injected, as well as between *GFP*-injected and *Mm_bGlc17* injected larvae according to two-tailed Student's *t*-tests (* $P < 0.5$). Error bars denote SEM. N = 6-8. D. TA-G deglycosylation activity (TA/(TA+TA-G)) of different RNAi silenced *M. melolontha* β-glucosidases and *GFP* as a negative control. TA-G deglycosylation of *Mm_bGlc16*, *Mm_bGlc18* and *GFP*, but not of *Mm_bGlc17* and *Mm_bGlc16&17* RNAi silenced larvae was higher than the boiled control samples. N = 9-15. *P*-value of a one-way

ANOVA is shown. The same lower case letters indicate no difference according to a Tukey's honest significance test. Error bars = SEM.

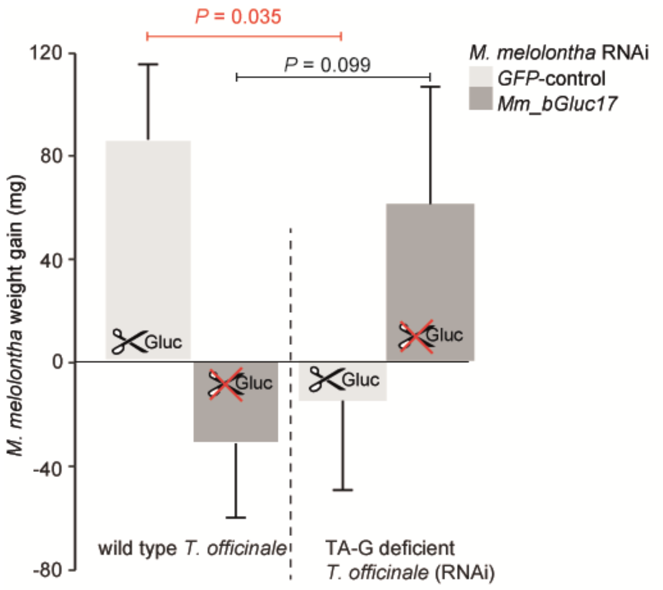


**Fig S6.** Weight gain of *Mm_bGlc17*-silenced and *GFP*-control *M. melolontha* larvae growing on transgenic TA-G deficient or control *T. officinale* lines. N = 11-15. *P*-values refer to Student's *t*-tests. Error bars = SEM.

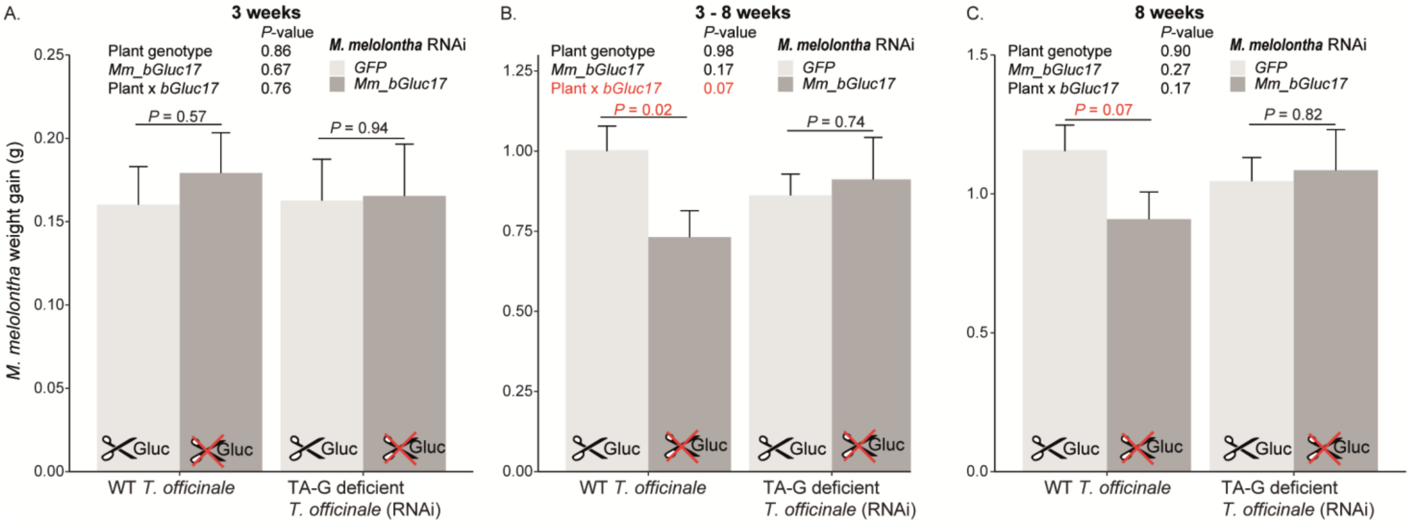

**Fig S7.** Weight gain of *Mm_bGlc17*-silenced and *GFP*-control *M. melolontha* larvae growing on transgenic TA-G deficient or control *T. officinale* lines within the first three weeks (A), three-eight weeks (B) and eight weeks (C). N = 12-20. *P*-values refer to two-way ANOVAs and Student's *t*-tests. Error bars = SEM.

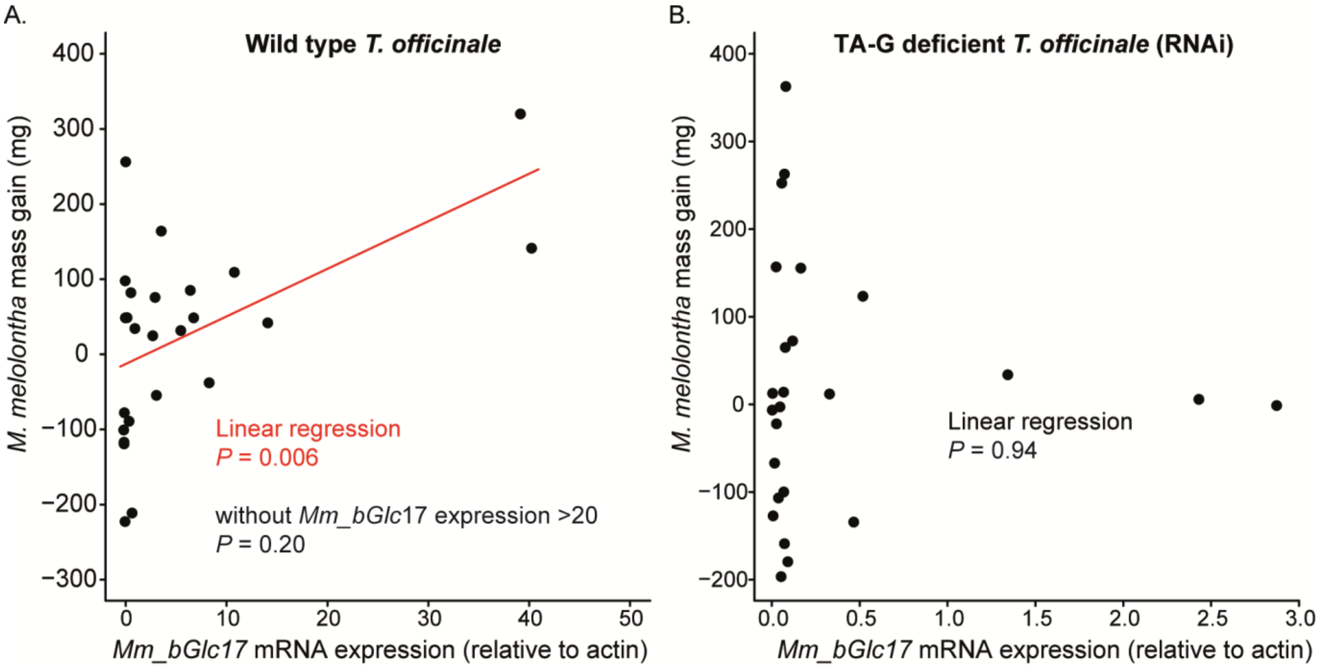


**Fig S8**. Correlation between *Mm_bGluc17* mRNA expression (relative to actin) and *M. melolontha* weight gain of *GFP*-control larvae growing for three weeks on TA- G containing wild type (A) or TA-G deficient transgenic *T. officinale* lines (B). *P*-values refer to linear regression analyses. The *P*-value of a separate linear model in which the two data points with *Mm_bGlc17* mRNA expression > 20 were excluded is also displayed.

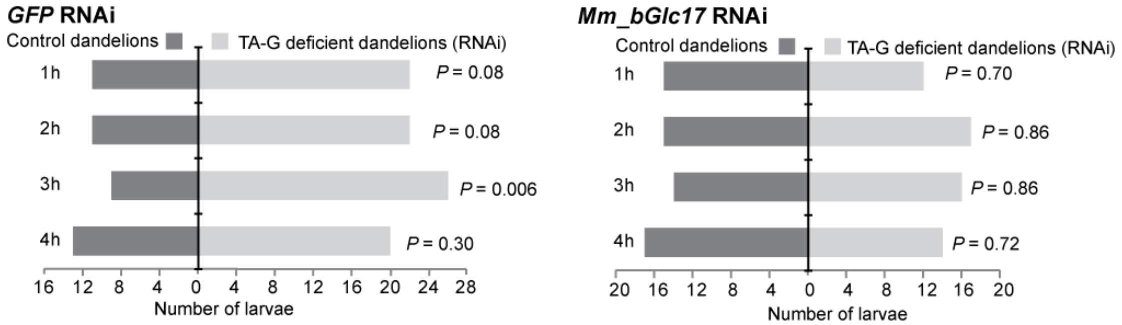


**Fig S9.** Choice of *M. melolontha* larvae that were treated with either *GFP* (left panel) or *Mm_bGlc17* (right panel) dsRNA between TA-G deficient transgenic and wild type *T. officinale* plants one to four hours after start of the experiment. *P*-values refer to binomial tests. Larvae that did not choose were excluded from the analysis. GFP = green fluorescent protein.

**SI tables**

**Table S1 Summary statistics of two-way ANOVA from Fig 2B.**

| | **Df** | **F value** | ***P*-value** |
|---|---|---|---|
| Gut section | 5 | 12.58 | 2.50e-08 *** |
| Heat treatment | 1 | 50.46 | 1.81e-09 *** |
| Gut:Treat | 5 | 13.52 | 8.63e-09 *** |

**Table S2. Summary statistics two-way ANOVA of Fig 2C.** TA-G: taraxinic acid β-D-glucopyranosyl ester; TA = taraxinic acid; GSH = glutathione; Cys = cysteine.

| | **Df** | **F value** | ***P*-value** |
|---|---|---|---|
| TA-G | | | |
| Plant enzyme | 1 | 2.517 | 0.141 |
| Tissue | 1 | 47.651 | 2.58e-05 *** |
| Plant enzyme*tis | 1 | 1.039 | 0.330 |
| TA | | | |
| Plant enzyme | 1 | 0.789 | 0.393 |
| Tissue | 1 | 0.973 | 0.345 |
| Plant enzyme*tis | 1 | 0.023 | 0.883 |
| TA-GSH | | | |
| Plant enzyme | 1 | 0.161 | 0.6960 |
| Tissue | 1 | 9.432 | 0.0106 * |
| Plant enzyme*tis | 1 | 0.038 | 0.8488 |
| TA-Cys | | | |
| Plant enzyme | 1 | 0.954 | 0.349621 |
| Tissue | 1 | 30.014 | 0.000192 *** |
| Plant enzyme*tis | 1 | 0.397 | 0.541695 |

**Table S3. Summary statistics of one-way ANOVA of Fig 4A.**

| | Df | F value | *P*-value |
|---|---|---|---|
| Silenced gene | 4 | 8.177 | 5.7e-05 *** |

**Table S4. Primer information.**

| Primer name | Primer sequences (5‘ -> 3‘) | Target gene |
|---|---|---|
| Primers to amplify full-length seuqences of *M. melolontha* β-glucosidases. | | |
| 1-pIB-fwd | GTAATGAAGCATCTACTATTATTTTTAAT | Mm_bGlc1 |
| 1-pIB-rev | CCATTCACAAACTTCGAATCCTTC | Mm_bGlc1 |
| 2-pIB-fwd | GTAATGAAACCAATCGGTTTAATTGTT | Mm_bGlc2 |
| 2-pIB-rev | CCAATCGCAAGCTTCAAATCCAT | Mm_bGlc2 |
| 3(2)-pIB-fwd | GTAATGAAATCACTAATTTTAATTTTAGTT | Mm_bGlc3 |
| 3(2)-pIB-rev | CCATTCGCAGACTTCGAATCCAT | Mm_bGlc3 |
| 5-pIB-fwd | GATATGAAAGCGATTATTATATTGGCT | Mm_bGlc5 |
| 5-pIB-rev | AAAATGCCTAATATAAATGTACAAAC | Mm_bGlc5 |
| 6neu-pIB-fwd | GTAATGAGGCGCGTCTTAATATTAATC | Mm_bGlc6 |
| 6neu-pIB-rev | TAGAATACTCAAGCTATGCATCAAG | Mm_bGlc6 |
| 11-pIB-fwd | GATATGAAAGTGCAGGTTGTATTAAT | Mm_bGlc11 |
| 11-pIB-rev | AAAATATCTCATACGAATAAATAATA | Mm_bGlc11 |
| 14-pIB-fwd | GATATGAGACGAATCATTTTCCTTTTG | Mm_bGlc14 |
| 14-pIB-rev | CTCTGGCACTTCGTCCTCGTCA | Mm_bGlc14 |
| 15-pIB-fwd | GTA ATGAAGCTCGTAATTTTCGCTCTG | Mm_bGlc15 |
| 15-pIB-rev | TGACCAATCGCACTCTTCGAAACC | Mm_bGlc15 |
| 16-pIB-fwd | GATATGAAGGTTCTAGTTATACTTTG | Mm_bGlc16 |
| 16-pIB-rev | CCATTCACAGTCGCTAAAATTTT | Mm_bGlc16 |
| 17-pIB-fwd | GATATGAAGAGACTAATTCTTATTTTC | Mm_bGlc17 |
| 17-pIB-rev | CCACTCGCAGGCATCAAATCCA | Mm_bGlc17 |
| 18-pIB-fwd | GATATGGGATACTTTGAACCATTAATA | Mm_bGlc18 |
| 18-pIB-rev | CCACCACGAACATGCTTCAAAT | Mm_bGlc18 |
| 19v-pIB-fwd | GATATGTCGTCATATGTATACAAAAAC | Mm_bGlc19v |
| 19v-pIB-rev | AACATCTTCATCTGACTCGGTA | Mm_bGlc19v |
| 5‘ RACE and nested-PCR primer | | |
| Mm_6-5’-RACE | CAATTTCGGCATCAATTTCGTTACTATCACTA | Mm_bGlc6 |
| Mm_6-5’-Nest | CACAAATCCTGCGTCCATACTTAATTGATA | Mm_bGlc6 |
| 3‘ RACE und Nested-PCR Primer | | |
| Mm_15-3’-RACE | CTGATACCGGTGAATTAAATGATTGTCGAAGAG | Mm_bGlc15 |
| Mm_15-3’-Nest | CAGTTCTAGAGGCAATCGTAGAAGATG | Mm_bGlc15 |

| | | |
|---|---|---|
| Mm_6-3'-RACE | TTATACTGTCCTTCTAGCTCATGCTAGAAC | Mm_bGlc6 |
| Mm_6-3'-Nest | TGCAGATGTTAATGCTGAAGATACGGC | Mm_bGlc6 |
| SMARTer RACE cDNA Amplification Kit Primer | | |
| Long Universal Primer | CTAATACGACTCACTATAGGGCAAGCAGTGGTATCAACGCAGAGT | |
| Short Universal Primer | CTAATACGACTCACTATAGGGC | |
| Nested Universal Primer | AAGCAGTGGTATCAACGCAGAGT | |
| Primers for dsRNA biosynthesis with and without the T7 promoter sequence (highlighted in green). | | |
| Mm-Tubulin-fwd | CACGCATACGACTTTGGA | |
| Mm-Tubulin-rev | GAGGCCGTGATTGAAGAT | |
| GFP-RNAi_fwd | GCTTTGCAAGATACCCAG | |
| GFP-RNAi_rev | GTTCATCCATGCCATGTG | |
| Mm_bGlc_16_fwd_T7 | TAATACGACTCACTATAGGGAGAACAGTATGGCACCTTTGGC | |
| Mm_bGlc_16_rev_T7 | TAATACGACTCACTATAGGGAGATCTAGCTATTCTTTCTTTCATGG | |
| Mm_bGlc_16_fwd | ACAGTATGGCACCTTTGGC | |
| Mm_bGlc_16_rev | TCTAGCTATTCTTTCTTTCATGG | |
| Mm_bGlc_17_fwd_T7 | TAATACGACTCACTATAGGGAGAACCCACTCCCATTTGTATTGG | |
| Mm_bGlc_17_rev_T7 | TAATACGACTCACTATAGGGAGAGATAGGCAATCTGGACCTCG | |
| Mm_bGlc_17_fwd | ACCCACTCCCATTTGTATTGG | |
| Mm_bGlc_17_rev | GATAGGCAATCTGGACCTCG | |
| Mm_bGlc_18_fwd_T7 | TAATACGACTCACTATAGGGAGATTCATATATTTTGTGAACTTGGAC | |
| Mm_bGlc_18_rev_T7 | TAATACGACTCACTATAGGGAGAGTCTTCAGACCTTCTTCGG | |
| Mm_bGlc_18_fwd | TTCATATATTTTGTGAACTTGGAC | |
| Mm_bGlc_18_rev | GTCTTCAGACCTTCTTCGG | |
| Primers for RT-qPCR | | |
| qPCR_Mm _bGlc_16_fwd | GACTCAGCAGTAGAGTTAGCATTACAGG | |
| qPCR_Mm _bGlc_16_rev | CGTTGTAAGTTCTGGTAGTCTGGATGC | |
| qPCR_Mm _bGlc_17_fwd | GAAGACATTCAAGCAGCTGAAGACAGTG | |
| qPCR_Mm _bGlc_17_rev | CCTCCGTAAAGATAGGCAATCTGGAC | |
| qPCR_Mm _bGlc_18_fwd | TACAAATGGAGTTTGGTTGGATAGCG | |
| qPCR_Mm _bGlc_18_rev | GGCACTCATCACGTCAATTCTGC | |
| qPCR_Mm _actin-fwd | CTGCCAGCTCAAGTTCCC | |
| qPCR_Mm _actin-rev | GAACAGAGCTTCTGGGCA | |
| qPCR_Mm_Tubulin_fwd | CACGCATACGACTTTGGAACAC | |

| | |
|---|---|
| qPCR_Mm_Tubulin_rev | GAGGCCGTGATTGAAGATACG |
